# Epigenetic-focused CRISPR/Cas9 screen identifies ASH2L as a regulator of glioblastoma cell survival

**DOI:** 10.1101/2022.08.17.504245

**Authors:** Ezgi Ozyerli-Goknar, Ezgi Yagmur Kala, Ali Cenk Aksu, Ipek Bulut, Ahmet Cingöz, Sheikh Nizamuddin, Martin Biniossek, Fidan Seker-Polat, Tunc Morova, Can Aztekin, Sonia H.A. Kung, Hamzah Syed, Nurcan Tuncbag, Mehmet Gonen, Martin Philpott, Adam P. Cribbs, Ceyda Acilan, Nathan A. Lack, Tamer T. Onder, H.T. Marc Timmers, Tugba Bagci-Onder

**Author notes:** Corresponding author Koç University School of Medicine, Koç University Research Center for Translational Medicine, Rumelifeneri Yolu, 34450, Sarıyer, Istanbul, Türkiye.

## Abstract

Glioblastoma is the most common and aggressive primary brain tumor with poor prognosis, highlighting an urgent need for novel treatment strategies. In this study, we investigated epigenetic regulators of glioblastoma cell survival through CRISPR/Cas9 based genetic ablation screens using a customized sgRNA library EpiDoKOL, which targets critical functional domains of chromatin modifiers. Screens conducted in multiple cell lines revealed *ASH2L*, a histone lysine methyltransferase complex subunit, as a major regulator of glioblastoma cell viability. *ASH2L* depletion led to cell cycle arrest and apoptosis. RNA sequencing and greenCUT&RUN together identified a set of cell cycle regulatory genes, such as *TRA2B, BARD1, KIF20B, ARID4A and SMARCC1* that were downregulated upon *ASH2L* depletion. Mass spectrometry analysis revealed the interaction partners of ASH2L in glioblastoma cell lines as SET1/MLL family members including SETD1A, SETD1B, MLL1 and MLL2. We further showed that glioblastoma cells had a differential dependency on expression of *SET1/MLL* family members for survival. The growth of *ASH2L*-depleted glioblastoma cells was markedly slower than controls in orthotopic *in vivo* models. TCGA analysis showed high ASH2L expression in glioblastoma compared to low grade gliomas and immunohistochemical analysis revealed significant ASH2L expression in glioblastoma tissues, attesting to its clinical relevance. Therefore, high throughput, robust and affordable screens with focused libraries, such as EpiDoKOL, holds great promise to enable rapid discovery of novel epigenetic regulators of cancer cell survival, such as *ASH2L*. Together, we suggest that targeting *ASH2L* could serve as a new therapeutic opportunity for glioblastoma.

## INTRODUCTION

Glioblastoma represents 45.6% of malignant primary brain tumors and occurs with 3.1/100 000 incidence per year^1^. Presence of necrotic foci, high proliferation rate, invasiveness, and highly angiogenic feature are hallmarks of glioblastoma, contributing to its very high lethality^2^. Median survival with standard care for patients is 14.6 months; and only 5.5% of patients can survive past 5 years after diagnosis^3^. Therefore, deeper understanding of genetic and epigenetic vulnerabilities of glioblastoma is crucial to design more effective therapies.

Epigenetic control of tumor progression has been widely studied, and the de-regulation of the expression of genes associated with tumor suppression or progression through epigenetic changes have been reported^4,5^. DNA methylation, histone modifications and chromatin remodeling are major epigenetic alterations, which broadly affect cell phenotype. Histone proteins are prone to a variety of dynamic posttranslational modifications (e.g. phosphorylation, acetylation, methylation, ubiquitination)^6^, which either modulate their affinity for the DNA or form new binding sites for protein modules, supporting the euchromatin (active) or heterochromatin (repressed) state^7^. These modifications are written, read or removed by unique proteins to regulate gene transcription, DNA repair, replication and chromosome condensation^8,9^. Abnormal DNA methylation and distinct histone modification patterns due to aberrant activity of epigenetic modifiers are frequently encountered in tumor cells and have effects on their drug response and growth^10,5^. Therefore, identifying epigenetic vulnerabilities of cancer cells can provide excellent therapeutic intervention points for various types of cancers.

CRISPR/Cas9 system is a widely used genome editing tool based on the delivery of gene– complementary synthetic guide RNA (sgRNA) with 3′-protospacer-associated motif (PAM) into the cell^11^. Cas9 endonuclease binds sgRNA and initiates site-specific cleavage of DNA, which ultimately results in insertions and/or deletions within targeted genes. CRISPR/Cas9-based functional genetic screens were shown to be more potent than shRNA screens^12,13^. Genome-wide CRISPR knockout screens with multiplexed sgRNAs enabled the identification of essentiality genes^13^ and vulnerabilities to drugs such as TRAIL^14^, ATR inhibitors^15^ and Ras inhibitors^16^ in various cancers. However, labor-intensive and expensive nature of genome-wide screens with low signal/noise ratio increased attention for focused libraries^17,18^. Additionally, different gene targeting approaches have been developed, since targeting the 5′ exons as default practice of most CRISPR applications, may produce in-frame functional variants of proteins. Therefore, as an alternative approach “domain specific targeting” was developed^19^, where the genomic regions encoding the functional protein domains were targeted by sgRNAs leading to higher proportion of null mutations.

Here, we introduce a customized <u>Epi</u>genetic <u>Do</u>main-specific <u>K</u>nock <u>O</u>ut <u>L</u>ibrary (EpiDoKOL), which targets functional domains of key epigenetic modifiers. Using EpiDoKOL, we performed drop-out screens on multiple cell lines and identified *ASH2L* as an indispensable gene for glioblastoma cell survival. ASH2L is a trithorax group family member that functions within the SET1/MLL family methyltransferase complexes specific for histone H3 lysine 4 (H3K4) methylation including MLL1, MLL2, MLL3, MLL4, SET1A and SET1B; and acts as a cofactor, supporting active gene transcription^20,21,22,23^. Here, we show that ASH2L directly regulates cell cycle-related genes and facilitates tumor cell survival both *in vitro* and *in vivo*. Together, our results suggest suitability of domain-targeted chromatin-focused CRISPR library screens for the identification of novel and druggable epigenetic vulnerabilities in glioblastoma.

## MATERIALS & METHODS

### Reagents and cell lines

All reagents and cell lines are described in **Supplementary Information**. RRID numbers for used cell lines, antibodies, organisms, plasmids, or tools are given in the related main or Supplementary method sections.

### Generation of EpiDoKOL

<u>Epi</u>genetic <u>Do</u>main-specific <u>K</u>nock <u>O</u>ut <u>L</u>ibrary (EPIDOKOL) targeting 251 critical chromatin modifier enzymes was developed by pooling sgRNAs designed against functional domains of target genes using CCtop tool^24^. The library consists of 1628 gene-targeting sgRNAs in total, in addition to 80 non-targeting sgRNAs retrieved from GECKO Library as negative control. 5 sgRNAs were picked for each domain of the targeted chromatin modifier genes. Designed sgRNAs were cloned to pLentiGuide (RRID:Addgene_52963) and plentiCRISPRv2 (RRID:Addgene_52961) backbones^25^ by GIBSON assembly. To determine sgRNA distribution in plasmid pools, next generation sequencing was performed on MISeq at MIT BioMicro Center. Library composition, sgRNA design, cloning and sequencing procedures are detailed in **Supplementary Information**. Sequences of sgRNAs of EpiDoKOL are available in **Supplementary Table 1**.

### Viral packaging and transduction

Lentiviral particles from plentiCRISPRv2 and retroviral particles from pBabe-hygro-GFP (RRID:Addgene_61215) were produced in 293T cells as described^26,27^. For stable cell line generation, cells were seeded at 1.5×10^6^ cells per plate in 10cm plates and were transduced with virus containing media supplemented with protamine sulfate (10µg/ml). Transduced cells were selected by antibiotics. Details are described in **Supplementary Information**.

### Cas9 activity assay

pBabe-hygro-GFP expressing T98G and U373 cells were seeded to cell culture plates and transduced with control sgRNAs g-NT1 and g-NT2 and GFP-targeting sgRNAs g-T1 and g-T2 separately. sgRNA sequences are listed in **Supplementary Table 2**. Percentage of GFP(+) cells were assessed by Flow cytometry. Details are described in **Supplementary Information**.

### Essentiality screen with EpiDoKOL

Screen was performed as two biological replicates. U373 and T98G cells were seeded and transduced with EpiDoKOL virus with MOI=0.4 to ensure that each cell takes up a single sgRNA. After transduction, a proportion of cells were pelleted to serve as a reference point for baseline sgRNA distribution while remaining cells were selected with puromycin. At day 30, cells were counted and pelleted. Collected pellets were stored at −80°C and genomic was DNA isolated using MN Nucleospin Tissue kit (Macherey-Nagel, Germany) according to manufacturers’ protocol. Isolated DNA was used for nested PCR. Sequences of external and internal PCR primers are listed in **Supplementary Table 3**. Samples were sent to Illumina Hiseq2500 RAPID sequencing to Vincent J. Coates Genomic Sequencing Laboratory of University of Berkeley. NGS results were analyzed using Python programming language and Model-based Analysis of Genome-wide CRISPR-Cas9 Knockout (MAGeCK) (version 0.5.8)^28^. p<0.05 cutoff was applied to gene-level analysis to identify significantly depleted genes. EpiDoKOL screen sequencing data are deposited to the NCBI GEO database with the accession number GSE201657. Detailed information on screening procedure, sequencing and statistical analysis are provided in **Supplementary Information**.

### sgRNA cloning for validation of hits

To knock-out candidate essentiality genes, top depleted sgRNAs sequences for *ASH2L, RBX1, SSRP1* were derived from EpiDoKOL and individually cloned to pLentiCRISPRv2 plasmid. All sgRNA sequences used for validation experiments are listed in **Supplementary Table 2**. Cloning procedure is detailed in **Supplementary Information**.

### siRNA transfection

Glioblastoma cells (5×10^4^) were seeded on 6-well plates. After 16h, cells were transfected with siRNA when 50-70% confluency was reached. For transfection *MLL1* (Ambion, 107890) and *WDR5* (Ambion, 136959) siRNAs were utilized. Transfection procedure is detailed in **Supplementary Information**.

### Colony formation assay

Control cells (g-NT infected) and cells carrying sgRNAs against candidate genes were seeded as 750 cells/well in triplicates in 6-well plates 7 days post-transduction. Colonies were grown for 14 days with fresh medium and then stained with Crystal Violet. Number of colonies were counted using ImageJ Software (NIH Image, Bethesda, MD, USA). Staining procedure is detailed **Supplementary Information**.

### GFP competition assay

7 days post-transduction, cells were seeded in 6-well plates together with their GFP-stable counterparts at equal proportion (150,000 cells each) and at final density of 300,000 cells per well for the GFP competition assay. At initial seeding day (day 0) and on days 2, 5, 7 and 9, the cells were detached with trypsin and the GFP-positive and negative cell ratio was determined by flow cytometry. Cells were analyzed by BD Accuri C6 (BD Biosciences, USA) (excitation 488 nm, emission 530/575 nm) recording 10,000 events per sample.

### Cell viability and apoptosis assays

Cell viabilities and Caspase 3/7 activities were measured via Cell Titer-Glo (CTG) Luminescent Cell Viability Assay (Promega, USA) or Caspase-Glo® 3/7 (Promega, USA) respectively, according to manufacturer’s instructions using a plate reader (BioTek’s Synergy H1, VT, USA). Annexin V/PI staining was performed with Muse® AnnexinV&Dead Cell Kit (Luminex, MCH100105) according to the manufacturer’s instructions and analyzed by Muse Cell Analyzer (Merck, Darmstadt, Germany). Western Blots for apoptotic markers; cleaved PARP (Abcam, ab74290, USA) and caspase3 (Cell Signaling, 5A1E) were also performed. Details of cell viability, caspase activation assays, Annexin V/PI staining, and Western Blotting are described in **Supplementary Information**.

### Histone extraction

Histone were acid-extracted, and protein concentration was determined with Pierce BCA Protein Assay Kit (Thermo Scientific, 23225). Histone extraction and western blotting details are described in **Supplementary Information**.

### Cell cycle assay

Cells were collected 14 days post-transduction, fixed with cold 70% Ethanol, stained with Muse Cell Cycle Reagent (The Muse® Cell Cycle Kit, Luminex, MCH100106) and analyzed with Muse Cell Analyzer (Merck, Darmstadt, Germany). Staining and cell cycle analysis procedure is detailed in **Supplementary Information**.

### Quantitative RT-PCR

List of primers can be found in **Supplementary Table 4**. Experimental details are available in **Supplementary Information**.

### TCGA

Gene expression profiles of “glioblastoma” (GBM) and “brain lower grade glioma” (LGG) tumors were preprocessed by the pipeline of The Cancer Genome Atlas (TCGA) consortium. Classical, Mesenchymal, Neural, and Proneural subgroups were identified. The log2-transformed gene expression values were compared against each other using the Wilcoxon rank sum test. Detailed explanation is available in **Supplementary Information**.

### Immunohistochemistry

Brain Glioblastoma tissue microarray (US Biomax-GL806f) was stained for ASH2L. Aperio AT2 Scanner (Leica Biosystems) was used for imaging slides and Aperio ImageScope (Leica Biosystems) program with Positive Pixel Count Algorithm was used for digital scoring as further described in **Supplementary Information**.

### RNA sequencing

U373 cells were transduced with *ASH2L* or *NT* sgRNAs. Cell pellets were collected as triplicates at 14 days post transduction for RNA isolation. Library preparation and sequencing was performed at University of Oxford (Oxford, UK). Data is deposited to the NCBI GEO database with the accession number GSE201657. RNAseq and statistical analysis procedures are detailed in **Supplementary Information**.

### Quantitative mass spectrometry of the ASH2L interactome

Sample preparation, GFP-affinity purification and data analysis were performed as reported^29,30^. Briefly, nuclear and cytoplasmic proteins were isolated from glioblastoma cells harboring ASH2L-GFP and used for GFP-affinity purification with GFP-Trap agarose beads (#gta-200, Chromotek). Purified proteins were on-bead digested and tryptic peptides were analyzed by nanoflow-LC-MS/MS with a high performance nanoflow-HPLC Orbitrap based ms/ms system (Thermo Fisher Scientific). The raw data files were analyzed with MaxQuant software (version 1.5.3.30). The obtained protein files were analyzed by Perseus software (MQ package, version 1.6.12). Detailed protocol is available in **Supplementary Information**. All mass spectrometry data have been deposited to the ProteomeXchange Consortium via the PRIDE partner repository (PXD033358).

### Genome localization experiments by greenCUT&RUN

Genome localization analysis of GFP-tagged ASH2L was performed by greenCUT&RUN with the combination of enhancer-MNase and LaG16-MNase, as described^31^. Sequencing libraries were prepared as described^29^ and sequenced in Illumina, HiSeq 3000 platform. Detailed experimental procedure and bioinformatic analyses of NGS are available in **Supplementary Information**. greenCUT&RUN sequencing datasets have been deposited to the Sequence Read Archive (SRA) portal of the NCBI with accession ID PRJNA828380.

### In vivo tumor growth

All *in vivo* experiments were approved by the institutional ethical committee of Koç University. 6-8-week-old non-obese diabetic/severe combined immunodeficiency (NOD/SCID) mice were used for orthotopic tumor models as described in **Supplementary Information**. Tumors were monitored using IVIS Lumina III (Perkin Elmer, USA). Quantification of tumor progression was performed with GraphPad PRISM software (San Diego, CA, USA).

## RESULTS

### Generation of EpiDoKOL and validation of the performance of the sgRNA library

To identify epigenetic modifier genes indispensable for cell survival, we generated a customized epigenome-wide domain-targeted pooled sgRNA library. In <u>Epi</u>genetic <u>Do</u>main-specific <u>K</u>nock<u>o</u>ut <u>L</u>ibrary (EpiDoKOL), sgRNAs were designed against functional catalytic domains of genes encoding chromatin modifiers. EpiDoKOL consists of 1750 sgRNAs in total, targeting 250 different genes, and 80 non-targeting controls **(Figure 1A)**. Molecular cloning of sgRNAs for pooled libraries into two different backbones (pLentiGuide and pLentiCrisprv2) was achieved by gibson assembly, leading to 800X coverage and successful lentiviral packaging **(Figure 1B, Supplementary Figure 1A)**.

**Figure 1.**
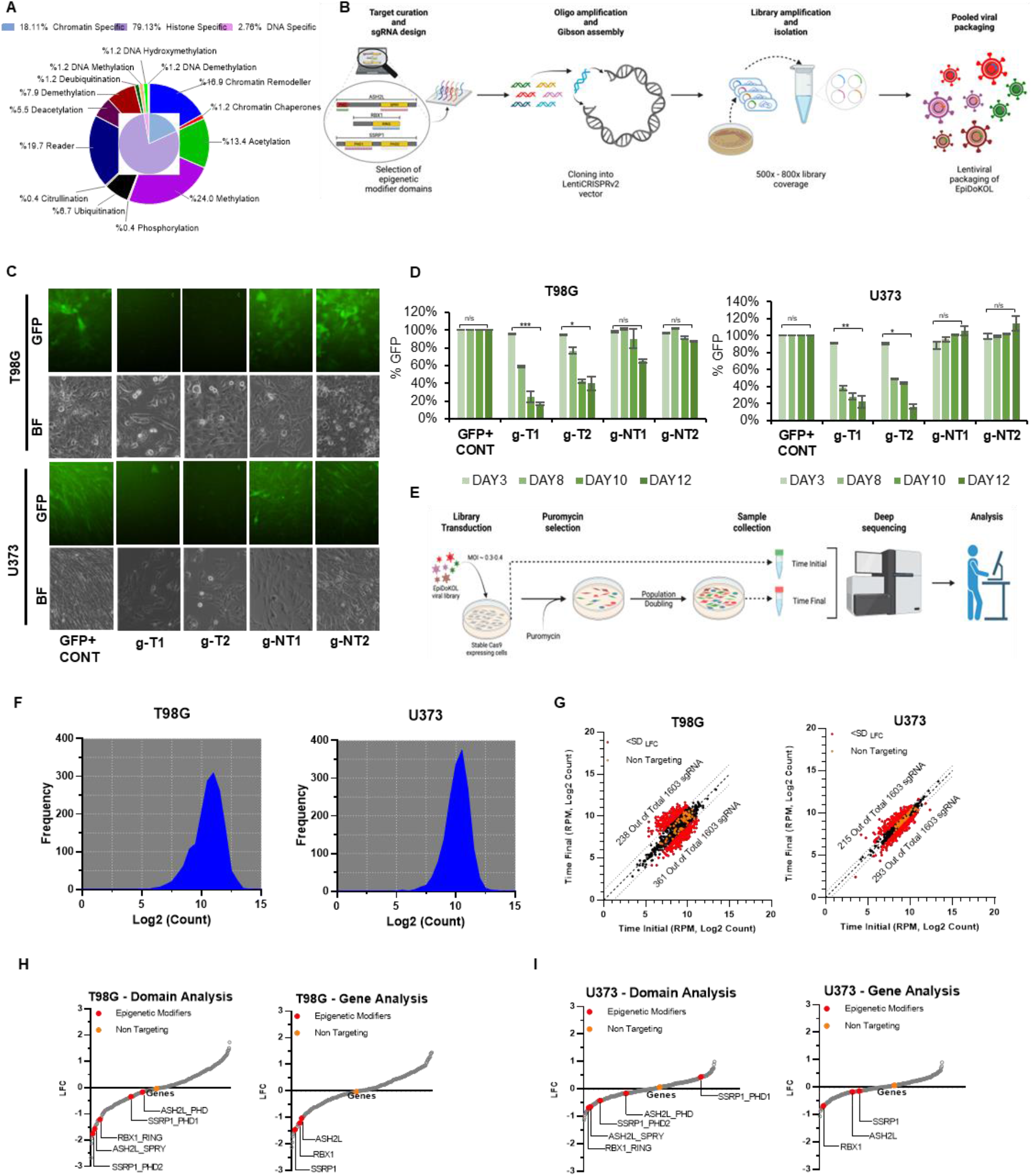
<u>Epi</u>genetic <u>Do</u>main-specific <u>K</u>nock <u>O</u>ut <u>L</u>ibrary (EpiDoKOL) screen identified common essentiality genes in T98G and U373 cells. **A**. Composition of EpiDoKOL in terms of target molecules, their functions and related epigenetic modifications. **B**. Library generation procedure. Figure created with BioRender.com **C**. Cas9 activity assay of U373 and T98G cells. Microscopic images of T98G and U373 cells transduced with indicated sgRNAs 12 days post-transduction. **D**. Flow cytometric analysis of U373 and T98G cells transduced with indicated sgRNAs (g-NT1, g-NT2: non-targeting sgRNAs; g-T1 and g-T2: sgRNAs targeting GFP) **E**. Schematic of EpiDoKOL screening procedure. Figure created with BioRender.com **F**. sgRNA density plots from cells transduced with pLentiCRISPRv2 plasmid containing EpiDoKOL. Cell pellets collected before puromycin selection. **G**. Log2 counts of sgRNAs at initial and final time points in T98G and U373 cells. **H**. Waterfall plots for Log2fold changes of genes after screening T98G cells with EpiDoKOL for a month. Domain and gene-based analysis revealed overlapping essentiality hits. **I**. Waterfall plots for Log2fold changes of genes after screening U373 cells with EpiDoKOL for a month. Domain and gene-based analysis revealed overlapping essentiality hits. Mageck p values were calculated with negative binomial model fitting. Other p values were determined by two-tailed Student’s t-test *P<0.05, **P<0.01, ***P<0.001.

As Cas9-based genetic ablation may be suboptimal in certain tumor cell lines due to chromosomal aberrations and mutational burden^32^, we first examined the efficiency of Cas9-mediated cleavage efficiency in glioblastoma cell lines. We first transduced U373 and T98G with GFP-encoding lentiviruses and then introduced g-NT1, g-NT2 (non-targeting sgRNAs), g-T1 and g-T2 (sgRNAs targeting GFP) along with Cas9 in a single lentiviral vector. Flow cytometry indicated that GFP+ cells were reduced to 20% of the total population at 12 days post-transduction in g-T1 or g-T2 transduced cells. No change in GFP signal was observed with g-NT1 and g-NT2 vectors, indicating that Cas9-activity is optimal in glioblastoma cells within two weeks (**Figure 1C-D**). We then continued with EpiDoKOL screen in both U373 and T98G cells. To identify common regulators of glioblastoma cell survival, cells were transduced with EpiDoKOL at a low multiplicity of infection (MOI=0.4) with 500x coverage to ensure single sgRNA intake per cell and proper sgRNA representation (**Figure 1E)**. Following antibiotic selection, transduced cells were cultured for 30 days by maintaining the sgRNA coverage at each population doubling. Using standard library preparation protocols, deep sequencing and analysis^33^, we compared the sgRNA composition in initial and final timepoints. Histogram of median normalized read counts for all sgRNA plotted for U373 and T98G cells confirmed normal distribution of log2-transformed normalized counts, indicating that no bias was introduced during cloning or transduction steps (**Figure 1F**). We then compared the depletion scores of gene-targeting versus non-targeting gRNAs to evaluate the overall efficiency of the screen. Essential gene targeting gRNAs were depleted significantly, whereas no change in non-targeting control sgRNA abundance was observed (**Figure 1G**). Altogether, these analyses illustrated that EpiDoKOL preserves normal distribution of sgRNAs in infected cells and reveals potential essential genes for cell survival.

### EpiDoKOL screen in T98G and U373 cells identified common regulators of glioblastoma cell survival

As EpiDoKOL was composed of multiple sgRNAs targeting a functional domain of a single gene, it was possible that one gene was targeted by more than 10 sgRNAs in total, depending on the domain composition. Therefore, we undertook two different analysis approaches; at domain level or gene-level and determined common depleted genes using median normalization scores (**Figure 1H-I**). Accordingly, several epigenetic modifiers were found to be significantly depleted in both glioblastoma cell lines. Along with epigenetic modifiers that were previously implicated in cancer cell fitness, such as *CHD1, CHD4, DNMT3B, ELP3, SUPT16H, SUV39H2*; novel hits *ASH2L, RBX1* and *SSRP1* were discovered as essential genes for both U373 and T98G suggesting common regulatory roles across various glioblastoma lines **(Supplementary Figure 1B)**.

To validate the function of these novel hits in glioblastoma, we individually introduced selected sgRNAs and assessed their effects with several functional experiments **(Figure 2A)**. Accordingly, colony forming abilities of U373 and T98G cells with individual knockouts of *RBX1, ASH2L, SSRP1* genes markedly decreased compared to control cells that received g-NT (**Figure 2B**). To assess the effect of ablation of hit genes in a heterogenous population, we performed GFP competition assays, in which GFP-negative cells were transduced with sgRNA against a hit gene and mixed in a 1:1 ratio with gNT-transduced GFP-positive cells. Monitoring both the GFP+ and GFP-fraction over 16 days, we observed gradual decreases in the cells transduced with selected sgRNA in U373 and T98G cells (**Figure 2C)** as well as U87MG cells (**Supplementary Figure 2A)**. Since the most significant phenotype was observed for U373 cells, we further delineated the effects of *RBX1, ASH2L, SSRP1* ablation on glioblastoma cell fitness using this cell line. Depletion of *ASH2L, RBX1* or *SSRP1* genes slowed down the growth of cells as gauged by cell viability assays (**Figure 2D**); and induced apoptosis as revealed by elevated caspase 3/7 activity (**Figure 2E**) and Annexin V and PI positivity **(Figure 2F**). Thus, cell death is likely to contribute to the reduced cell numbers observed upon *ASH2L, RBX1* and *SSRP1* ablation. These proof-of-principle experiments illustrated that EpiDoKOL is a practical functional genomics tool that enables identification of epigenetic modifiers important for cancer cell fitness.

**Figure 2.**
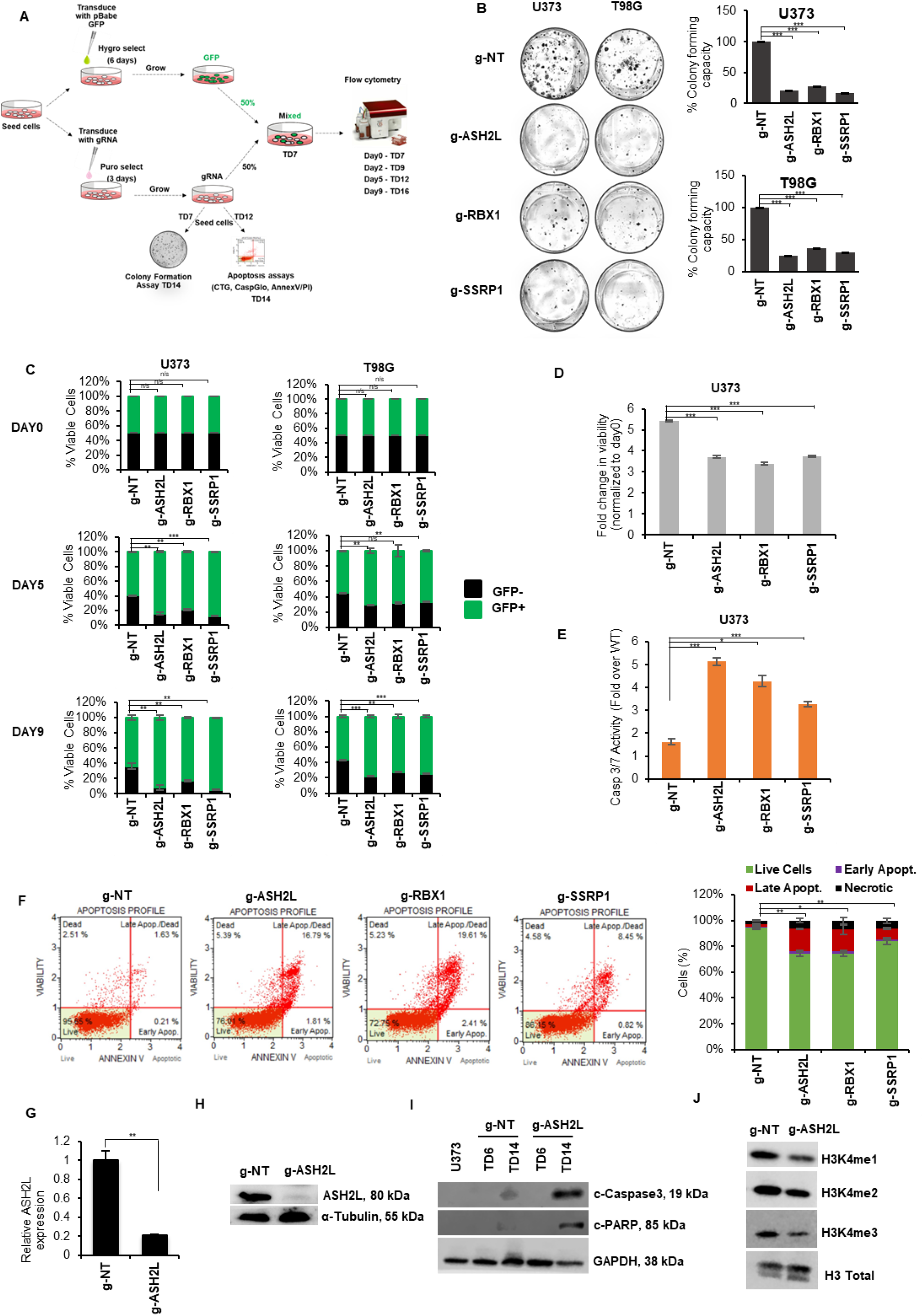
Effects of candidate genes on glioblastoma cell fitness were validated with functional assays in vitro. **A**. Scheme of validation experiments for novel EpiDoKOL essentiality hits. TD: post-transduction day **B**. Representative images of long-term clonogenic assay of cells infected with sgRNAs against selected hits and statistical analysis. Quantification of colonies was performed by ImageJ software. **C**. Results of GFP competition flow cytometric assay for selected hits. **D-F**. Cell viability (D), Caspase 3/7 activity (E) and AnnexinV & PI analysis (F) conducted on U373 cells upon depletion of ASH2L, RBX1 and SSRP1 genes. **G**. qPCR analysis for ASH2L mRNA levels upon transduction of U373 cells with g-ASH2L or g-NT. **H**. Western blot analysis of ASH2L protein levels upon transduction of U373 cells with g-ASH2L or g-NT. **I**. Western Blot analysis for cleaved Caspase3 and PARP in U373 cells at day 6 and 14 post-transduction with g-NT control or g-ASH2L. **J**. Western Blot analysis for H3K4 mono and trimethylation levels in U373 cells 14 days post-transduction with g-NT control or g-ASH2L. P values determined by two-tailed Student’s t-test *P<0.05, **P<0.01, ***P<0.001.

### ASH2L is essential for glioblastoma cell survival and regulates histone methylation and transcription

As the most pronounced phenotype was observed with *ASH2L* gene ablation, we further investigated its role in glioblastoma. To this end, we knocked out *ASH2L* in U373 and U87MG cells and showed successful depletion of *ASH2L* both in protein and mRNA level **(Figure 2G-H, Supplementary Figure 2B**). Consistent with previous results, extensive cleavage of caspase-3 and PARP was observed indicating potent induction of apoptosis upon ASH2L depletion **(Figure 2I**). ASH2L functions as a cofactor within MLL family of methyltransferase complexes to trigger histone H3 lysine 4 (H3K4) methylation^20,21,22,23^, and its loss was previously associated with reduction of H3K4 trimethylation. Consistently, we observed reduced mono and trimethylation of H3K4 upon *ASH2L* depletion in U373 and U87MG cells **(Figure 2J, Supplementary Figure 2C**). Since H3K4me3 is a well-known euchromatin mark acting globally, we investigated gene expression changes upon *ASH2L* knock-out to gain a mechanistic understanding for its essential role. We performed RNA-seq comparing U373 cells transduced with control and *ASH2L* sgRNAs (**Figure 3A)**. *ASH2L* knockout resulted in 461 up-regulated (FDR<0.05, p-value<0.05 and log2-fold-change ≥1) and 1076 down-regulated (FDR<0.05 and log2-fold-change≤-1) genes. Number of downregulated genes was higher than upregulated ones, consistent with the association between H3K4 trimethylation and gene activity (**Figure 3B)**. GSEA revealed several negatively and positively enriched pathways with significant normalized enrichment scores (**Figure 3C, Supplementary Figure 3A)**. Downregulated genes upon *ASH2L* depletion significantly overlapped with cell cycle and mitosis-associated gene sets, whereas upregulated genes enriched for metabolism, glycolysis, and hypoxia pathways. Expression of representative downregulated hits were validated by qRT-PCR (**Supplementary Figure 3B**).

**Figure 3.**
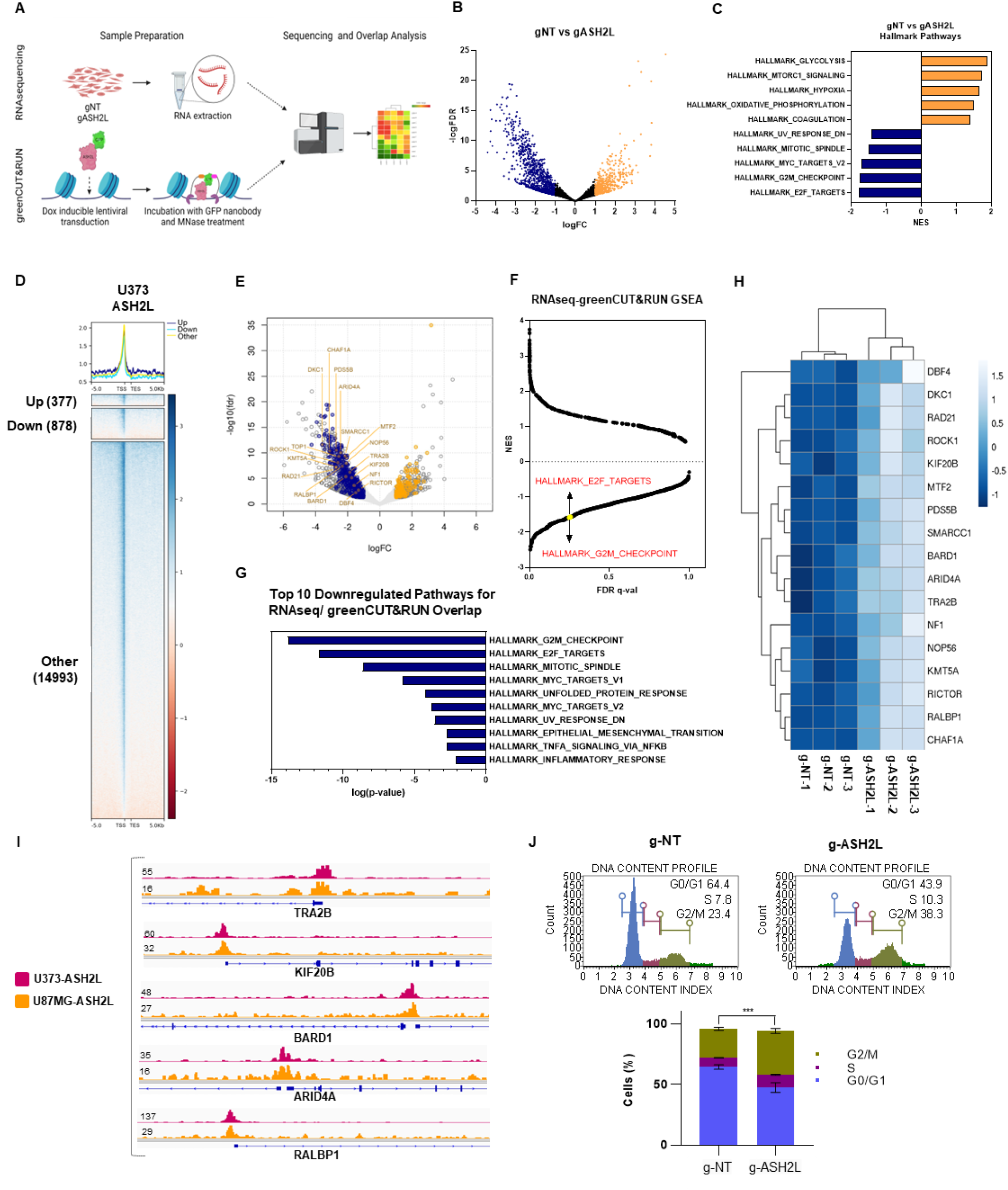
ASH2L transcriptionally regulates cell cycle via direct promoter interactions with G2/M-checkpoint and E2F target gene sets. **A**. Schematic of RNAseq and greenCUT&RUN experiments. **B**. Volcano plot of RNAseq data showing differentially expressed genes (DEGs) in ASH2L depleted U373 cells compared to controls on the 14 day post-transduction **C**. Normalized enrichment score results of gene set enrichment analysis (GSEA) for all gene sets available from MSigDB v7.5. Top 5 biological processes enriched in downregulated or upregulated genes upon ASH2L knockout are shown. P values were calculated by hypergeometric test. **D**. greenCUT&RUN analysis in U373 cells revealing genomic localization of ASH2L at differentially expressed gene promoters **E**. Venn diagram and volcano plot of overlapped RNAseq and greenCUT&RUN data to show proportion of differentially expressed genes bound by ASH2L in U373 cells. Genes bound by ASH2L and differentially expressed upon ASH2L depletion are now called as critical genes, denoted by blue (for downregulated) and yellow (for upregulated) dots on volcano plot. **F**. Normalized enrichment score and FDR-qval results of GSEA on critical genes for all gene sets available from MSigDB v7.5. Some of the negatively enriched pathways related to cell cycle are highlighted. **G**. Top 10 biological processes enriched in regulatory genes upon ASH2L knockout. **H**. RNAseq heatmaps of critical genes **I**. greenCUT&RUN representative igv plots of critical genes. **J**. Flow cytometric cell cycle analysis of ASH2L-depleted U373 by PI staining on post-transduction day 14 and its statistical analysis. P values determined by two-tailed Student’s t-test *P<0.05, **P<0.01, ***P<0.001.

Regulation of the transcriptome by ASH2L prompted us to analyze its localization on chromatin using CUT&RUN in both U373 and U87MG cells. Using the default criteria of the HOMER (FDR<0.0001 for calling the peaks and fold changes≥4 and p value < 0.001 against control for filtering the peaks), 13360 and 1964 chromatin peaks were called in U373 and U87MG cells, respectively (**Figure 3D**). To find the commonalities between these two cell lines, initially all the peaks were merged, and coverage was compared. In total, 13782 merged peaks were identified, of which, 117 (0.85% of total merged peaks, 5.96% of the U87MG peaks) and 4493 (32.6% of total merged peaks and 33.6% of the U373 peaks) were found unique for the respective cell lines. Conversely, 94.1% of U87MG peaks and 66.4% of U373 peaks were common indicating direct targets of ASH2L in glioblastoma cells (**Supplementary Figure 3C**).

### ASH2L is critical for cell cycle regulation

We explored genes bound by ASH2L on chromatin and differentially expressed upon ASH2L depletion as major targets of ASH2L. By overlapping RNAseq and greenCUT&RUN results, we generated a list of genes that may be directly linked with the essential function of ASH2L. When ASH2L peaks on promoter-TSSs were overlapped with DEGs from RNA-seq data, a total 150 of up-regulated and 350 down-regulated genes were identified, which together constitute a “critical gene list” (**Figure 3E)**. GSEA conducted on this list led to the identification of G2/M checkpoint, E2F targets and mitotic spindle hallmark gene sets as enriched (**Figure 3F-G)**. Heatmap for differential expression of cell cycle and mitosis related genes as well as greenCUT&RUN IGV tracks for selected genes are shown as examples **(Figure 3H-I, Supplementary Figure 3D)**. Based on our finding of cell cycle regulatory genes as ASH2L targets, we conducted cell cycle analysis on *ASH2L*-depleted U373 and U87MG cells. Propidium iodide staining revealed decreased number of cells in the G0/G1 and accumulation at S and G2/M phases of cell cycle upon ASH2L knockout, resembling mitotic arrest (**Figure 3J, Supplementary Figure 2D)**. Collectively, these results suggest that ASH2L plays a critical role in maintenance of glioblastoma cell fitness through regulating cell cycle progression.

### Glioblastoma cells have differential dependency on SET1/MLL family of transcription factors interacting with ASH2L

To address the essential role of ASH2L for glioblastoma cell survival further, we focused our attention to chromatin modifying complexes, specifically, SET1/MLL family of histone methyltransferases SET1 (SETD1A), SET1B (SETD1B), MLL1 (KMT2A), MLL2 (KMT2B), MLL3 (KMT2C) and MLL4 (KMT2D)^34^. Each of these methyltransferases can form distinct multi-subunit complexes with diverse functions. All SET1/MLL families contain four core subunits, WDR5, RBBP5, ASH2L and DPY30, which are collectively named as the WRAD module^35^. Other subunits of SET1/MLL family complexes are specific to one or a few SET1/MLL families and involved in chromatin recruitment^36^. Being part of WRAD module, ASH2L acts as a cofactor to support trimethylation^20,21,22,23^. To assess which SET1/MLL family complexes ASH2L is mainly part of in glioblastoma cells, we performed quantitative mass spectrometry (MS) of the ASH2L interactome for U373 and U87MG cell lines. Cells were transduced with inducible N-terminal GFP fusions of *ASH2L* gene encoding N-GFP-ASH2L fusion protein. Nuclear and cytoplasmic proteins were extracted, GFP-affinity purified and subjected to tandem MS (**Figure 4A)**. Accordingly, we identified common members of MLL-family and SETD1A/B complexes (DPY30, WDR5, ASH2L, RBBP5, HCFC1); as well as with MLL-specific (KMT2A, KMT2B, KMT2C, KMT2D, NCOA6, KDM6A, MEN1, PSIP1, PAGR1, PAXIP) and SETD1A/B-specific (SETD1A, SETD1B, WDR82, BOD1, CXXC1) complex members in nuclear fraction. Based on LFQ values, we determined the stoichiometry of interacting proteins (**Figure 4B-D)**. While in cytoplasmic fraction, interaction was evident only with SETD1A/B complex (**Supplementary Figure 4A-B**); in nuclear fraction ASH2L interacted with MLL family and SETD1A/B members with similar stoichiometry (**Figure 4B-D)**. To assess the functional importance of SET1/MLL family complex members for glioblastoma cell viability, we performed siRNA mediated knock-down of WDR5 and KMT2A (MLL1) in U373 and U87MG cells and checked their colony forming ability (**Figure 4E)**. Downregulation of MLL1 significantly reduced colony forming potential of both U373 and U87MG cells, whereas WDR5 silencing affected only U87MG cells (**Figure 4F)**. Taken together, these results implicate ASH2L-MLL1 complex as a specific dependency in GBM cell proliferation.

**Figure 4.**
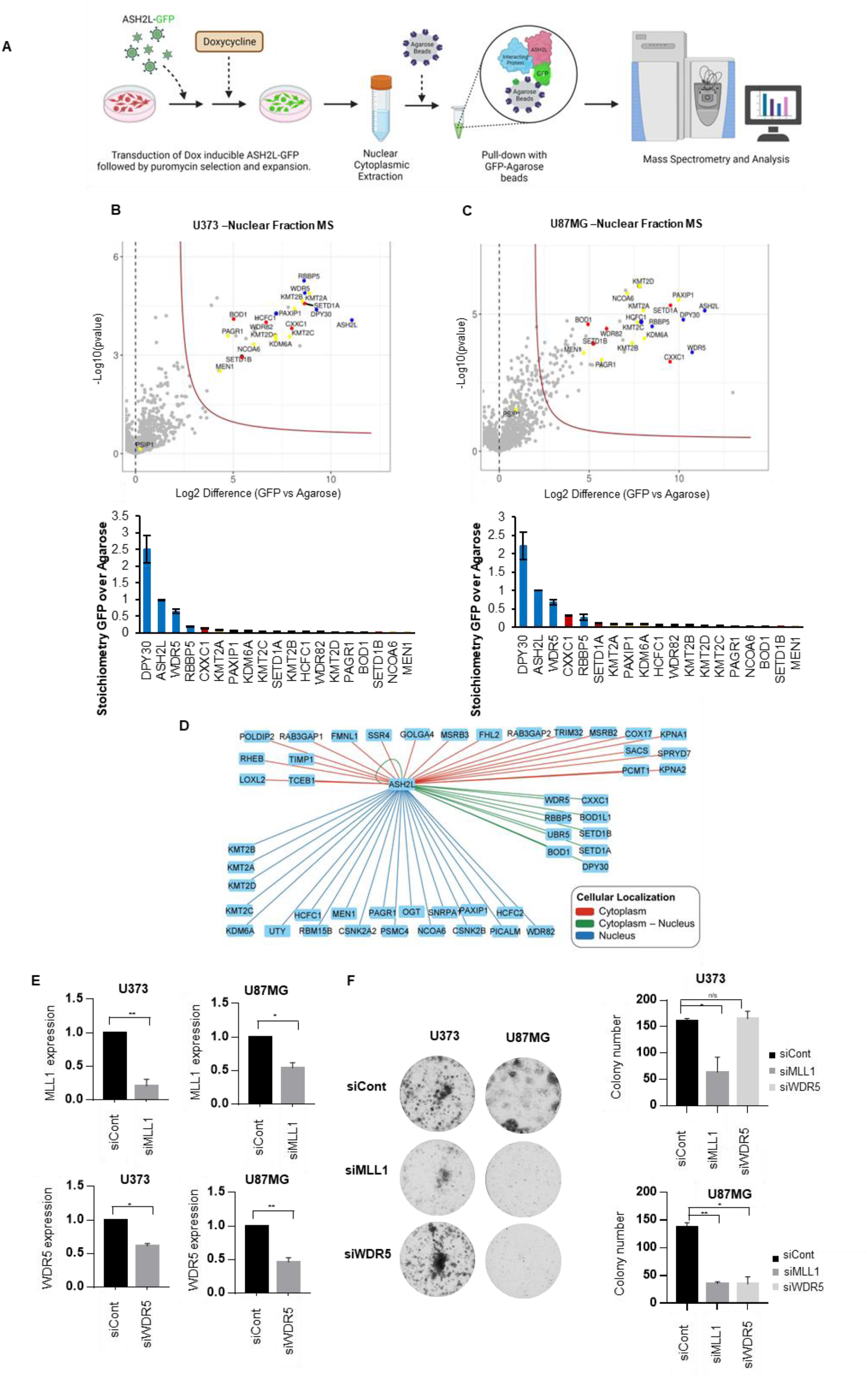
Glioblastoma cells have differential dependency to SET1/MLL family of transcription factors interacting with ASH2L. **A**. Scheme of tandem mass spectrometry experiments performed on glioblastoma cells. **B-C**. Proteomic analyses of GFP-tagged ASH2L in U373 (B) and U87MG (C) cells. Volcano plots of significant interactors of GFP-ASH2L isolated from nuclear extracts are shown. Stoichiometry plots of bound SET1/MLL family members (MLL family specific: yellow, SETD1A/B specific: red, MLL & SETD1A/B common members: blue) are depicted. All interactors are normalized to the GFP-ASH2L bait. Results shown represent Intensity Based Absolute Quantification with standard deviations. **D**. Distribution of ASH2L common interactors in terms of subcellular localization in U373 and U87MG cells. **E**. qPCR analysis for mRNA levels upon siRNA mediated knockdown of WDR5 and KMT2A genes in U373 and U87MG cells. **F**. Representative images of long-term clonogenic assay upon siRNA mediated knockdown of WDR5 and KMT2A genes and statistical analysis. Quantification of colonies was performed by ImageJ software. P values determined by two-tailed Student’s t-test *P<0.05, **P<0.01, ***P<0.001.

### ASH2L is essential for tumor progression *in vivo*

To assess the role of ASH2L in glioblastoma growth *in vivo*, we utilized orthotopic xenograft models using Luciferase (Fluc)-expressing U87MG cells, stably transduced with control sgRNA or ASH2L sgRNA. Following transduction, cells were intracranially injected and tumor growth was monitored until 33 days post-implantation **(Figure 5A)**. Accordingly, U87MG cells with ASH2L knockout could not form tumors as compared to the control cells. **(Figure 5B)**. Striking block of tumor forming capacity by lack of ASH2L encouraged us to check available patient data. Indeed, ASH2L expression was higher in glioblastoma patients in comparison to low grade gliomas; ASH2L was expressed in all glioblastoma subtypes, but enriched for proneural subtype, based on TCGA **(Figure 5C-D)**. We next examined ASH2L protein in patient samples by immunohistochemistry utilizing a glioblastoma tissue microarray with 80 cores representing 40 different glioblastoma cases. 70% of glioblastoma specimens had strong ASH2L expression, whereas nonmalignant tissue specimens had none/low expression **(Figure 5E, Supplementary Figure 5)**. Collectively, these clinically relevant results illustrate that ASH2L plays a critical role in glioblastoma progression.

**Figure 5.**
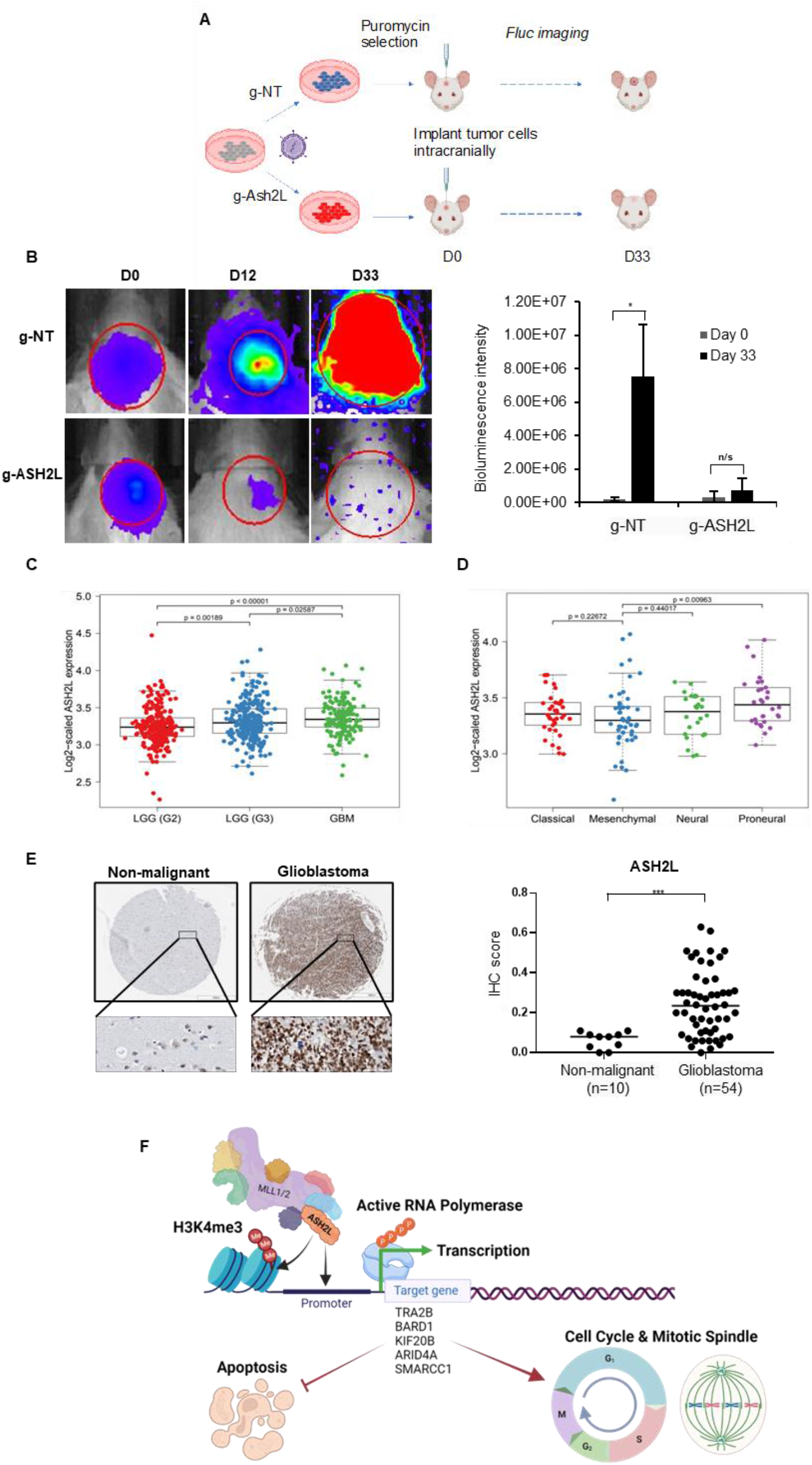
ASH2L is essential for glioblastoma tumor growth *in vivo*. **A**. Scheme for in vivo validation of ASH2L essentiality for GBM via intracranial injection of g-ASH2L or g-NT transduced U87MG cells (n=5 per group) and bioluminescence detection. **B**. Representative images of 3 mice taken during bioluminescence measurements at days 0-12-33 were illustrated. Bioluminescence signal of tumors formed by g-ASH2L or g-NT transduced U87MG cells were compared 33 days post-injection. **C**. The boxplots displaying the ASH2L gene expression ratio (RSEM normalization values) in low grade gliomas (LGG) and GBM based on TCGA **D**. The boxplots displaying the ASH2L gene expression ratio in GBM subtypes based on TCGA **E**. Representative core images from Brain Glioblastoma tissue microarray (TMA) stained with anti-ASH2L antibody. Scale bar 500 μm. **F**. Percentage of ASH2L-positive cores were shown (n=80 cores). P values determined by two-tailed Student’s t-test *P<0.05, **P<0.01, ***P<0.001 **G**. Model of ASH2L essentiality for glioblastoma cell survival. ASH2L together with DPY30, RBBP5 and WDR5 forms WRAD module, which acts as cofactor of SET1/MLL family transcription factors (MLL1/2, MLL3/4, SETD1A, SETD1B) to bind promoters of target genes and induce methylation of H3K4, a mark for euchromatin state. Active RNA polymerase binds to open chromatin to initiate transcription of genes involved in cell cycle progression, regulation of mitotic spindles and survival of cells (e.g TRA2B, BARD1, KIF20B, ARID4A, SMARCC1). Deregulation of ASH2L levels interferes with cell cycle and leads to cell cycle arrest and apoptosis.

## DISCUSSION

Epigenome-directed treatment strategies interfering with abnormal DNA methylation, acetylation or chromatin remodeling patterns of glioblastoma are heavily investigated. For example, clinical outcome of conventional treatment with Temozolomide (TMZ) is tightly dependent on DNA tumors’ methylation status. Because patients with methylated MGMT gene have survival advantage^37^, synthetic inhibitors of MGMT were tested in glioblastoma patients (NCT00613093)^38^. HDAC inhibitors also hold promise for better clinical outcome^39,40^. Valproic acid with TMZ and radiotherapy (NCT00302159) gave promising results^41,42^. Belinostat combined with TMZ is under clinical investigation (NCT02137759), with encouraging results in terms of recurrence delay and decreased psychological symptoms^43^. Inhibitors of chromatin remodeling complex, namely Oliparib and Veliparib are FDA-approved and clinically investigated for glioblastoma as single agents or in combination with chemo/radio therapy (NCT01390571, NCT03212274, and NCT02152982, NCT03581292, NCT01514201 respectively) with promising results^44,45^.

Based on the ever-growing importance of the epigenetic mechanisms in tumor progression and therapy response, we investigated epigenetic regulators of glioblastoma cell survival through a functional screen approach using our custom-generated targeted library, EpiDoKOL designed against functional domains of epigenetic modifier proteins. sgRNAs targeting essential domains generate the strongest lethality phenotypes and thus provide a strategy to rapidly define the protein domains required for cancer dependence^46^. Targeting functional domains results in higher proportion of null mutations as opposed to conventional CRISPR approaches targeting 5’ exon of genes which might create truncated proteins that retain functionality. We identified *ASH2L, CHD1, CHD4, DNMT3B, ELP3, RBX1, SSRP1, SUPT16H, SUV39H2* as essential regulators of the viability of multiple cell lines. Some of the gene hits from EpiDoKOL screen were in concordance with previous reports. To exemplify, suppression of *SSRP1* decreases proliferation of malignant glioma through modulation of *MAPK* signaling^47^. *SSRP1* also facilitates hepatocellular carcinoma malignancy^48^. *SSRP1* and *SUPT16H* are components of FACT complex and responsible for quick malignant transformation in various cancer types^49^. Inhibition of FACT complex eliminates tumor initiating cells and increases survival in preclinical studies^50^. *RBX1* is E3 ubiquitin ligase and its repression attenuates tumor proliferation through cell cycle arrest, programmed cell death and senescence^51^. Essentiality of chromatin remodeling factor *CHD1* is demonstrated in *PTEN*-deficient cancer^52^. *CHD4* is an essential gene in breast cancer^53^ and an important regulator of colorectal cancer malignancy. *CHD4* has oncogenic functions in initiating and maintaining epigenetic silencing of multiple tumor suppressor genes^54^. An essential role for DNA methyltransferase *DNMT3B* in cancer cell survival has previously been illustrated^55^. *SUV39H2* is a potential oncogene in lung adenocarcinoma^56^ and osteosarcoma^57^. *ELP3* can initiate tumorigenesis and regeneration in the intestine^58^. Finally *WDR5* upregulation promotes proliferation, self-renewal and chemoresistance in bladder cancer via H3K4 trimethylation^59^. Identification of these already-established cancer fitness genes by EpiDoKOL attests to the power and utility of our screening strategy.

Our experiments focused on the characterization of the role of *ASH2L*, one of the four well-conserved core subunits of SET1/MLL family complexes since it has not been previously studied in glioblastoma. Emerging evidence implicates ASH2L in regulating cell proliferation and fate. *ASH2L* is crucial for euchromatin state and pluripotency in ESC^60^. *ASH2L* enhances Tbx1, Pax7, or Mef2-mediated transcriptional activity in embryogenesis and stem cell differentiation by increasing H3K4me3 levels^61,62,63^. *ASH2L* has also oncogenic properties^64^. ASH2L interacts with MYC to enhance MYC-mediated gene transcription and cooperates with activated H-RAS to transform rat embryo fibroblasts^64,65^. It participates in the crosstalk between H2B ubiquitination and H3K4 methylation to modulate gene transcription^66^. Additionally, ASH2L has been shown to play important roles in the progression of multiple tumors. ASH2L upregulates GATA3-induced transcription of ESR1 in breast cancer cells^64^. ASH2L is recruited to the promoter region of apoptosis-related genes, co-activating p53 to promote apoptosis in colorectal cancer^67^. ASH2L is highly expressed in cervical cancer, and its depletion inhibits HeLa cell proliferation^64^. *ASH2L* is involved in promotion of endometrial cancer progression via upregulation of *PAX2* transcription^68^. *ASH2L* drives proliferation and sensitivity to bleomycin and other genotoxins in Hodgkin’s lymphoma and testicular cancer cells^69^. Low expression of ASH2L protein correlates with a favorable outcome in acute myeloid leukemia^70^.

In line with previous evidence, we observed the following in glioblastoma cells upon *ASH2L* depletion: 1) colony formation and proliferation of the cells were reduced; 2) apoptosis was induced; 3) transcription dynamics were altered; 4) cell cycle kinetics of cells changed, and G2/M transition of cell cycle was blocked; and 6) tumor forming capacity of glioblastoma cells was inhibited. The RNA-seq data of *ASH2L*-depleted cells showed a high number of downregulated genes consistent with the necessity of the WRAD core complex to establish H3K4me1 and H3K4me3 at active enhancers and promoters, respectively^71,72,73^. We observed downregulation of *MYC* target genes (*SMARCC1, NOP56*) upon *ASH2L* depletion consistent with previous reports. Additionally, cell cycle, mitotic spindle and E2F transcription factor associated genes^74^ (e.g. *BARD1, PDS5B, RAD21, SMARCC1, KMT5A, ROCK1, NF1, RICTOR*) were downregulated upon *ASH2L* depletion. Together, we generated a model for involvement of *ASH2L* in glioblastoma cell survival (**Figure 5F**). Accordingly, the SET1/MLL family complex and H3K4 tri-methyltransferase have been reported to participate in the coordination of cell cycle progression and glioblastoma proliferation^75^. Similarly, knockout of *ASH2L* resulted in G2/M arrest accompanied with a proliferation defect in hematopoietic progenitor cells^76,77^. Moreover, greenCUT&RUN analysis revealed that ASH2L directly occupied the core promoter regions of these downregulated cell cycle regulatory genes implying a direct regulation. On the other hand, in RNAseq analysis a small but considerable number (337) of upregulated genes were identified upon ASH2L depletion and those genes were also direct targets of ASH2L. These upregulated genes were enriched in metabolism; EMT, glycolysis and hypoxia related gene sets and needs further investigation to decipher their surprising negative regulation by ASH2L.

Quantitative mass spectrometry of the ASH2L interactome for multiple cell lines revealed ASH2L interaction with MLL family and SETD1A/B members with similar stoichiometry in nucleus. We observed that glioblastoma cells had differential dependency to SET1/MLL family of epigenetic factors, since cells showed differential response to *MLL1* and *WDR5* ablations. Essentiality of WDR5 together with ASH2L for glioblastoma cells emphasizes the importance of WRAD module for cancer cell fitness. *RBBP5*, another component of the module, has recently been classified as a novel oncogene involved in gliomagenesis. Amplification and overexpression of the *RBBP5* gene were found in patient-derived tumor samples^78^.

*ASH2L*-ablated glioblastoma cells had reduced tumor forming capacity *in vivo* and ASH2L expression was high in glioblastoma tissues, attesting to ASH2L’s clinical relevance. Though we illustrated the essentiality of ASH2L for glioblastoma cell fitness, whether this essentiality stems from proteins’ function in SET1/MLL family complexes and to what extent each family member contributes to cell fitness remain elusive and will be subject to further investigation.

Despite the complexity and heterogeneous nature of cancer, epigenetic therapies hold great promise for improved survival of patients due to their potential of resetting the cancer epigenome. Detection of epigenetic factors modulating tumor survival via high throughput, robust and affordable screens such as EpiDoKOL holds great potential to enable rapid discovery of novel cancer-related mechanisms and development of effective therapies.

## ACKNOWLEDGEMENTS

Financial support was obtained from The Scientific and Technological Research Council of Turkey (TUBITAK) (1003-216S461). The authors acknowledge the use of the services and facilities of the Koç University Research Center for Translational Medicine (KUTTAM), funded by the Presidency of Strategy and Budge, Türkiye. We also thank to Nareg Pınarbaşı-Değirmenci and Özlem Yedier-Bayram for their assistance with experimental analysis and image design.

## AUTHOR CONTRIBUTIONS

Study design: TBO, EOG, FSP, CA, NAL, TTO; data generation: EOG, EYK, ACA, IB, AC, SN, MB, SHYK, MP, APC; data analysis: EOG, EYK, ACA, IB, APC, SN, TM, HS, NT, MG; data interpretation: TBO, EOG, HTMT; supplied reagents: CAA, TTO, HTMT; manuscript draft: EOG and TBO; approved final manuscript: all authors.

