## Supplementary information for "Epigenetic-focused CRISPR/Cas9 screen identifies ASH2L as a regulator of glioblastoma cell survival"

#### 1- SUPPLEMENTARY FIGURES AND LEGENDS

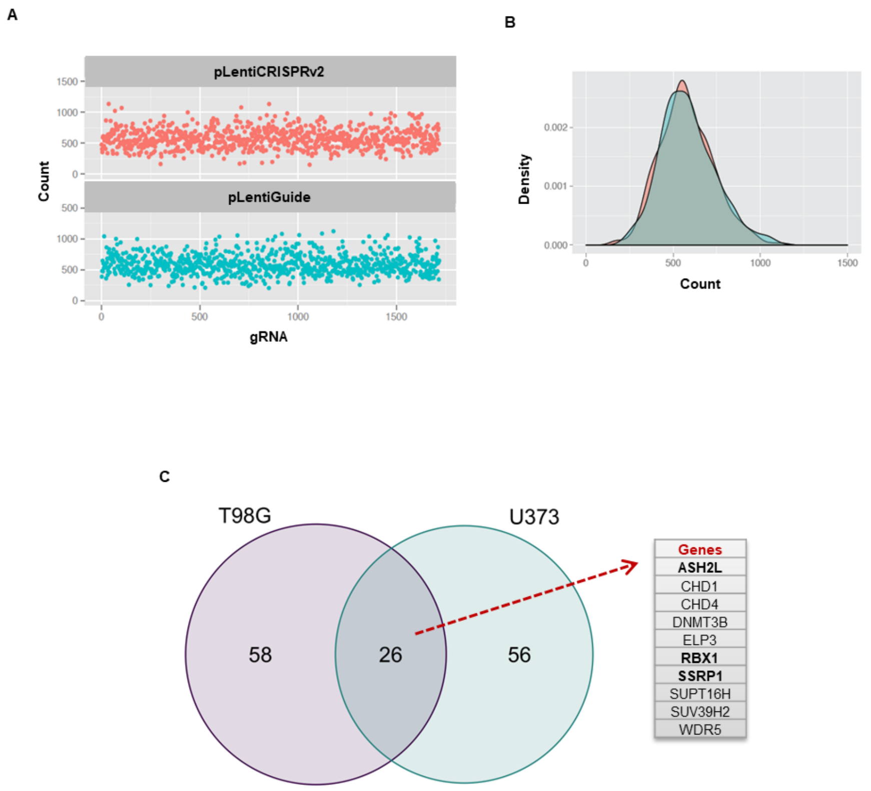

**Supplementary Figure 1. Quality Check of EpiDoKOL in plasmid level.**
**A.** sgRNA count and density plots for EpiDoKOL in pLentiGuide or pLentiCRISPRv2 backbones **B.** Common hits of EpiDoKOL screens on T98G and U373 cell lines identified by LFC-SD comparison. 26 genes were identified as common.

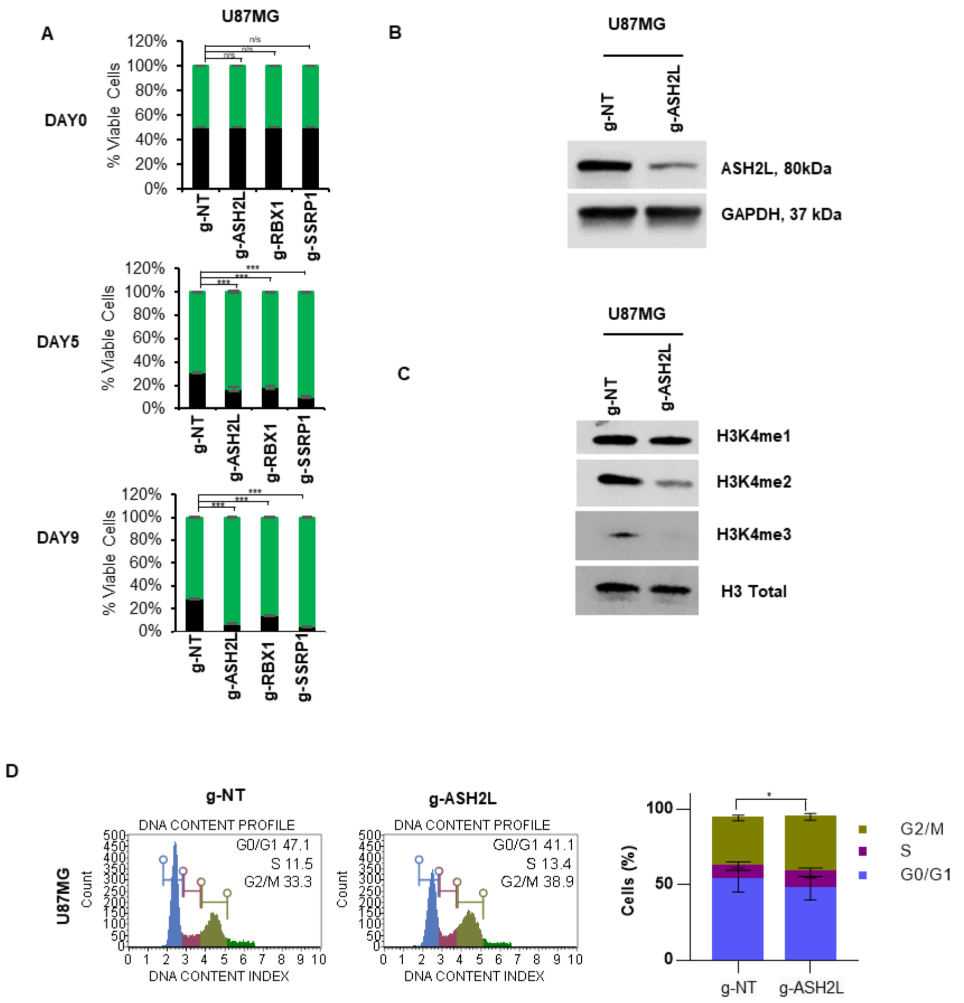

**Supplementary Figure 2. Effects of candidate genes on U87MG cell fitness.**
**A.** Results of GFP competition flow cytometric assay for selected hits in U87MG cells. **B.** Western blot analysis of ASH2L protein levels upon transduction of U87MG cells with g-ASH2L or g-NT. **C.** Western Blot analysis for H3K4 mono and trimethylation levels in U87MG cells 14 days post-transduction with g-NT control or g-ASH2L. **D.** Flow cytometric cell cycle analysis of ASH2L-depleted U87MG by PI staining on post-transduction day 14 and its statistical analysis. P values determined by two-tailed Student's t-test *P<0.05, **P<0.01, ***P<0.001.

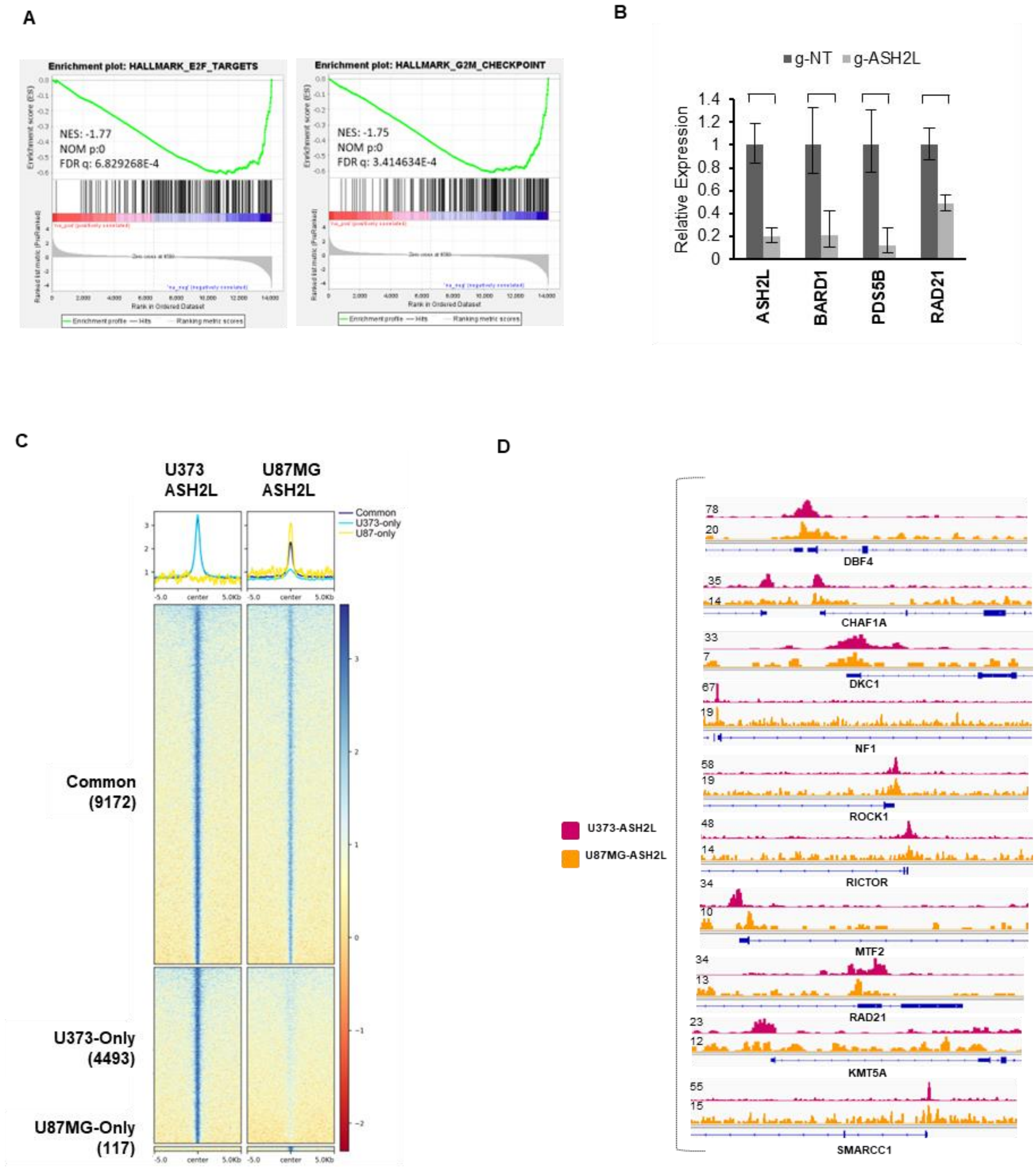

**Supplementary Figure 3. RNAseq and greenCUT&RUN analyses in glioblastoma cells.**
**A.** Enrichment plots of cell cycle related gene sets are depicted for U373 cells **B.** qPCR analysis for mRNA levels of representative critical genes upon g-NT control or g-ASH2L transduction of U373 cells. **C.** ASH2L greenCUT&RUN analysis in U87MG and U373 cells reveals overlapping genomic localization of ASH2L. **D.** greenCUT&RUN representative igv plots of critical genes

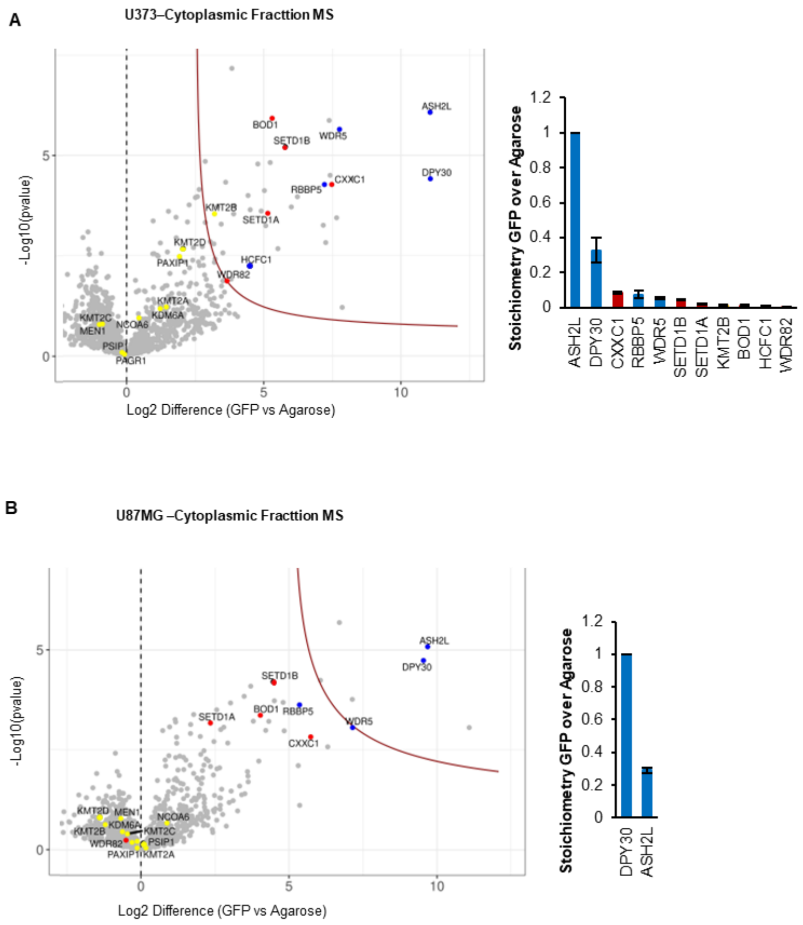

**Supplementary Figure 4. Cytoplasmic interactions of ASH2L in glioblastoma cells.** Volcano plots of significant interactors of GFP-ASH2L isolated from cytoplasmic extracts of **A.** U373 and **B.** U87MG cells are shown. Stoichiometry plots of bound SET1/MLL family members (MLL family specific: yellow, SETD1A/B specific: red, MLL & SETD1A/B common members: blue) are depicted. All interactors were normalized to the GFP-ASH2L bait. Results shown represent Intensity Based Absolute Quantification with standard deviations

A

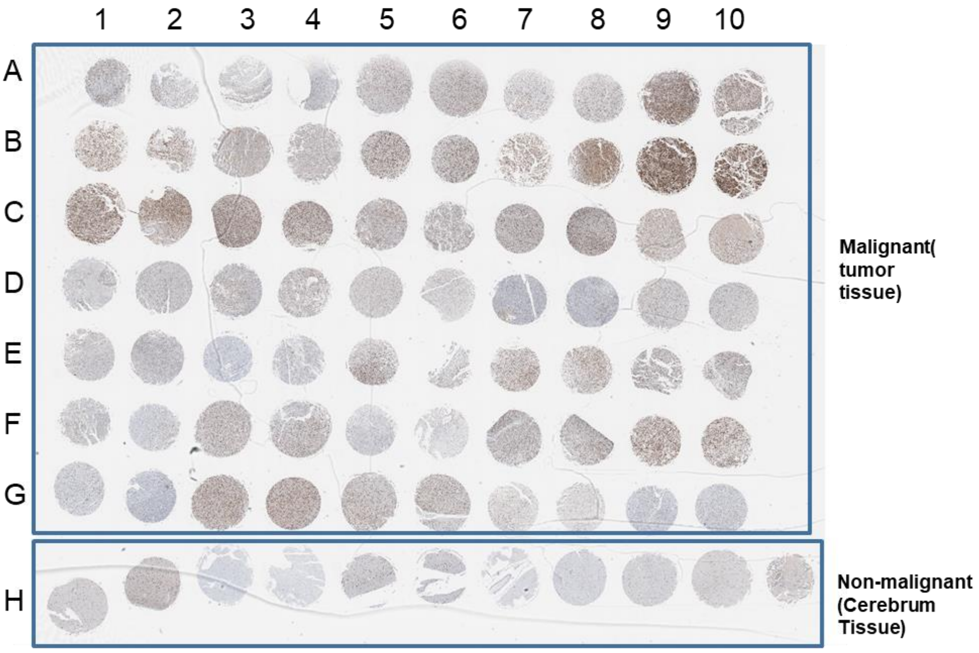

B

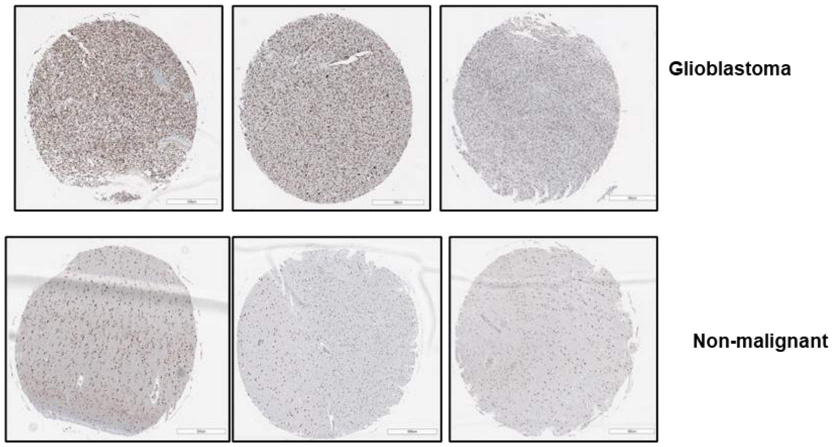

**Supplementary Figure 5. ASH2L immunohistochemistry in glioblastoma.**
**A.** Scanned brain glioblastoma tissue microarray (TMA) slides stained with ASH2L antibody. A1-G10 (malignant tissue), H1-1H10 (non-malignant tissue). **B.** Representative tissue cores, scale bar 500 μm

### 2-SUPPLEMENTARY METHODS

**sgRNA composition of EpiDoKOL:** We generated an Epigenetic Domain-specific Knock Out Library (EPIDOKOL) targeting 251 critical chromatin modifier enzymes by pooling guide RNAs designed against functional domains of target genes using CCtop tool[1]. The library consists of 1628 gene-targeting sgRNAs in total, in addition to 80 non-targeting sgRNAs serving as negative control. sgRNAs were designed against functional catalytic domains of chromatin modifier genes. Information on protein sequences of conserved functional domains was retrieved from NCBI Unigene software. Protein sequences of the domains were traced back to coding exonic sequences by UCSC Blat software and then given as input to CCTop to yield candidate sgRNA sequences[1]. Four-step criteria was followed to choose appropriate sgRNA for efficient targeting; 1) sgRNAs with non-exonic targets were ignored, 2) Exonic off-targets of sgRNA should have more than 3 mismatches, 3) Mismatches between sgRNA and its target gene sequence should be after 8 bp downstream from 5'-end and 4) sgRNA should not contain repetitive TTTTTT sequence. In case there was no sgRNA matching to the criteria mentioned above, we utilized GECKO library driven sgRNAs targeting the first coding exon[2]. In total 5 sgRNAs were picked for each domain of the targeted chromatin modifier genes. 80 non-targeting sgRNAs retrieved from GECKO Library were included in the library as negative controls.

**Cloning and sequencing of EpiDoKOL**: Designed sgRNAs were cloned to pLentiGuide and lentiCRISPRv2 backbones[3] by GIBSON assembly as described[4]. Briefly, all sgRNAs were synthesized as pooled oligonucleotides (LC Biosciences). Lyophilized oligonucleotides were resuspended and amplified by PCR with following conditions: For 50 µl of total PCR mix, 10 µl of 5X HF buffer (NEB, USA), 1 µl of dNTP mix (10 mM) (Thermo Fisher, USA), 2.5 µl of forward primer (10 µM), 2.5 µl of reverse primer (10 µM), 0.1 µl of resuspended oligomix, 0.5 µl of Phusion High-fidelity DNA polymerase (NEB, USA), 33.4 µl of dH2O were mixed. Thermal cycler conditions were set: 30s at 98ºC for denaturation, followed by 20 cycles of (10s at 98ºC, 20s at 63ºC, 15s at 72ºC), 3 min at 72ºC for final extension. Two tubes of 50 µl reaction were run on agarose gel and correct-sized bands were gel-extracted. Backbone vectors were digested by BsmB1 endonuclease (in Buffer3.1) at 60°C for 3h, run on 1% agarose gel, extracted from the gel and subjected to antarctic phosphatase (AP) treatment at 37°C for 1h followed by 15min 65°C enzyme inactivation. PCR-amplified oligo DNAs were ligated to pLentiGuide or lentiCRISPRv2 through GIBSON assembly (NEB, USA) in 1:5 ratio, incubated at 50ºC for 1h. Endura™ electrocompetent cells were transformed with the ligation product through seven electroporations in parallel. Transformed bacteria were seeded on large bioassay petri dishes (Corning). In total, ~$1.4 \times 10^6$ colonies were picked from seven plates to establish 800x library coverage. Plasmid pool was isolated by Maxi-prep Kit (Qiagen, USA). To determine sgRNA distribution in plasmid pools, plasmid DNAs were amplified by adding Illumina compatible sequences. 10 ng template DNA were mixed with 0.5 µl Phusion High-Fidelity DNA Polymerase (NEB, USA), 10 µl 5x GC Buffer (NEB, USA), 1 µl dNTP mix (10 mM each) (Thermo Fisher, USA), 2 µl Forward Stagger Mix (10 µM) and 2 µl Reverse Index Primer (10 µM) and Nuclease-free water (NEB,USA) up to 50 µl. Thermal cycler conditions were as follows: denaturation for 30s at 98ºC, followed by (10s at 98ºC, 15s at 63ºC, 20s at 72ºC) for 16 cycles, final extension for 4 mins at 72ºC. PCR amplicons were gel extracted using NucleoSpin Gel and PCR clean-up kit (Macherey-Nagel, Germany) according to the manufacturer's instructions and quantified using Nanodrop (ThermoFisher Scientific, USA). Next generation sequencing was performed on MISeq at MIT BioMicro Center with at least 10 million reads. Normal distribution of sgRNAs was verified.

**Cell culture:** U87MG, U373 and T98G cell lines were purchased from American Tissue Type Culture Collection (ATCC) and authenticated. 293T cells were kind gift of Dr. Tamer Onder (Koç University, Turkey). Cells were grown in DMEM medium (Gibco, USA) supplemented with 10% fetal bovine serum (Gibco, USA) and 1% Pen/Strep (Gibco, USA) in humidified incubator at 37ºC with 5% CO2 level.

**Viral packaging and transduction:** Viral packaging was performed as described[5,6]**.** Briefly, on day 0, 2.5x$10^6$ HEK 293T cells were seeded to 10 cm culture dish with DMEM supplemented with 10% FBS and 1% Pen/Strep. Viral plasmid DNA (2,5 µg), packaging plasmids Gag-Pol (2,250 µg of pUMVC or 8.2DeltaVPR for retroviruses and lentiviruses respectively) and VSVG (250 ng) were transfected to cells using FugeneHD Transfection Reagent (Promega, USA) in serum-free DMEM. Next day (minimum 8 hours after transfection), medium was changed and 48 and 72 hours post-transfection, media containing virus was collected and filtrated by 0.45 µm low protein binding filters. Viruses were concentrated using PEG-8000 (Sigma) and stored at -80ºC. For stable cell line generation, cells were seeded at 1.5 x $10^6$ cells per plate in 10 cm plates and were transduced with virus containing media supplemented with protamine sulfate (10 µg/ml). 16 hours post-transduction, viral medium was replaced by fresh media. Transduced cells were selected by Puromycin (1µg/ml for 3 days) or Hygromycin (300 µg/ml for 6 days) depending on the resistance cassette on the plasmid. After selection, cells were allowed to proliferate in fresh and antibiotic-free DMEM media.

**Cas9 activity assay:** Viral packaging of negative control sgRNAs g-NT1 and g-NT2 and GFP targeting sgRNAs g-T1 and g-T2 were performed as explained above. For Cas9 activity assay, pBabe-hygro-GFP expressing T98G and U373 cells were seeded to cell culture plates and transduced with indicated viruses separately. Percentage of GFP (+) cells are assessed by Flow cytometry starting from 2 days post-transduction untill GFP signal in g-T1/g-T2 transduced cells significantly diminishes. For flow cytometric analysis, cells were trypsinized and harvested. Cell pellets were washed two times with PBS, centrifuged at 1500 rpm for 5 min. Cell pellets were resuspended in 1x PBS supplemented with 1% FBS. BD Accuri C6 Flow Cytometer (BD Biosciences, USA) was used to analyze the samples according to the manufacturer's instructions.

**Essentiality screen with EpiDoKOL**: Screen was performed as two biological replicates. U373 and T98G cells were seeded and transduced with EpiDoKOL virus with low MOI (MOI=0.4) to ensure that each cell takes up a single sgRNA among 1718 sgRNAs in the library. 2.082.424 cells (calculated for 400 cell/sgRNA representation with MOI value in consideration) were seeded for viral transduction. After viral transduction, a proportion of cells were pelleted to serve as a reference point for baseline sgRNA distribution while remaining cells were selected with puromycin (2 mg/ml) for 3 days. Selected cells were regularly passaged and grown in cell culture for 18 days so that Cas9 gets active in glioblastoma cell lines. Cells were then separated into 2 sets. Set1 was composed of 687.200 cells (400 cell per guide representation) seeded in 6-well plate, Set2 was composed of 14 x $10^6$ cells seeded in 15 cm plates (8000 cell per guide representation). Two sets of cells were grown and passaged regularly for 12 more days. At day 30, cells were counted and pelleted. Collected pellets were stored at -80ºC until genomic DNA isolation.

**Genomic DNA isolation and Nested PCR**: Genomic DNA (gDNA) was isolated from pellets by MN Nucleospin Tissue kit (Macherey-Nagel, Germany) according to manufacturers' protocol. Isolated DNA was used for nested PCR, which amplifies sgRNA sequences and tag them with Illumina adaptor sequences for next generation sequencing. Input gDNA amount for PCR was calculated as 250x coverage of the EpiDoKOL, which corresponded to 13.2 µg per sample (assuming 6.6 pg DNA per cell). In external PCR gDNA was mixed with 1 µl Phusion High-Fidelity DNA Polymerase (NEB, USA), 20 µl 5x GC Buffer (NEB, USA), 2 µl dNTP mix (10 mM each) (Thermo Fisher,USA), 5 µl Forward External Primer (10 µM), 5 µl Reverse External Primer (10 µM) and Nuclease-free water (NEB, USA) up to 100 µl on ice. Thermal cycler conditions were set as follows: denaturation for 3 mins at 95ºC, followed by (25 s at 95ºC, 20 s at 65ºC, 15 s at 72ºC) for 17 cycles, final extension for 3 mins at 72ºC. For internal PCR, 5 µl from combined PCR products were used as a template with 5 µl Forward Stagger Mix (10 µM) and 5 µl Reverse Index Primer (10 µM) in a 100 µl reaction. Internal forward primers were composed of pooled staggered primers. Each sample was amplified by a distinct Index primer for barcoding. Thermal cycler conditions were the same as external PCR, except that amplification was carried out for 23 cycles instead of 17. Final amplicons from duplicate internal PCRs were gel extracted using NucleoSpin Gel and PCR clean-up kit (Macherey-Nagel, Germany) according to manufacturer's instructions and quantified using Nanodrop (ThermoFisher Scientific, USA).

**Next generation sequencing and statistical analysis**: Samples were sent to Illumina Hiseq2500 RAPID sequencing to Vincent J. Coates Genomic Sequencing Laboratory of University of Berkeley. NGS results were analyzed using Python programming language and Model-based Analysis of Genome-wide CRISPR-Cas9 Knockout (MAGeCK) (version 0.5.8)[7], which reports positively and negatively selected genes by applying robust rank aggregation (RRA) algorithm[8] to the rankings of genes in a pathway. Reads from R1 fastq files were counted at sgRNA level and normalized to library size. Biological replicates were presented as individual input files during sgRNA counting. Individual counts were combined as one output count for each sgRNA in every condition with median normalization to obtain gene-level log fold changes. $p<0.05$ cutoff was applied to gene-level analysis to identify significantly depleted genes. sgRNA counts were also normalized as Read Per Million (RPM) and converted to Log2 values[9,10]. Density plots of the Log2 transformed sgRNA counts were plotted with R using the geom_density function of the ggplot2 package to generate kernel density estimation (KDE) plots. Pearson correlations were calculated and plotted with R using the pairs.panels function in the psych package. Cumulative density plots were plotted with stat_ecdf function in ggplot2 package in R. Graphpad program was utilized to sketch plots of enriched/depleted sgRNAs based on their RRA scores and normalized read counts. EpiDoKOL screen sequencing data are deposited to the NCBI GEO database with the accession number GSE201657.

**sgRNA cloning for validation of hits:** Candidate essentiality genes were predicted by comparing initial and final-time point samples. To knock-out essentiality genes, top depleted sgRNAs sequences were derived from EpiDoKOL**.** For cloning, oligonucleotides encoding top and bottom strands of sgRNA against target genes were commercially purchased. For annealing reaction 1 µl from top and bottom strands (from 100 µM stock) was mixed with 6.5 µl nuclease-free water, 1 µL T4 ligase buffer with ATP (10X) and 0.5 µL T4 PNK. Mixture was run in PCR machine with thermal conditions: 37°C 30 mins, 95°C 4 mins, ramp down to 25°C (5°/min), infinite hold at 4°C. Annealed oligonucleotides were diluted 1/200 in nuclease free water and 1 µL of diluted sample was used for ligation reaction. Annealed samples were used for ligation into pLentiCRISPRv2 vector. Vector was digested with BsmB1 enzyme (in Buffer3.1) at 60°C for 3 h, run on 1% agarose gel, excised from gel and cleaned up from agarose and then treated with AP at 37°C for 1 h followed by 15 min 65°C enzyme inactivation. For ligation reaction the insert was mixed with 50 ng processed vector, 7.5 µL Quick Ligase buffer (2X), 1 µL Quick Ligase (Roche, Switzerland) and nuclease free water up to 20.5 µL reaction. Ligation was performed at RT for 15 minutes. Ligated vector was transformed to competent bacteria Stbl3 mixing 50 µL competent bacteria with ligation reaction, keeping on ice for 15 minutes, heat shocking bacteria at 42°C for 30 seconds, adding 150 µL LB without antibiotic and growing bacteria at 37°C 225 rpm for 1 hour. Transformed bacteria were spread on Ampicillin containing LB agar plates and colonies were grown overnight. Grown colonies were picked and grown in Ampicillin containing LB Broth for 16 hours to proliferate. Plasmids were isolated by MN miniprep kit and sent for sequencing by mixing with U6 forward sequencing primer (ACTATCATATGCTTACCGTAAC).

**siRNA experiments**: Glioblastoma cells ($5x10^4$) were seeded on 6-well plates. After 16h, cells were transfected with siRNA when 50-70% confluency was reached. 5 µL of Lipofectamine™ 3000 reagent was diluted in 200 µL of Opti-MEM™ and 100 pmol siRNA was mixed with 200 µL of Opti-MEM™ in separate tubes. After 5 min, diluted RNA and Lipofectamine 300 Reagent were mixed in 1:1 ratio and incubated at RT for 20 min. Following incubation, RNA and lipid complex was added to the wells dropwise. For transfection *MLL1* (Ambion, 107890), *MEN1* (Ambion, 143227) and *WDR5* (Ambion, 136959) siRNAs were utilized.

**Colony formation assay:** To assess relative cell fitness, control cells (g-NT infected) and cells carrying sgRNAs against candidate genes were seeded as 750 cells/well in triplicates in 6-well plates 7 days post-transduction. Colonies were grown for 14 days with fresh medium and then stained with Crystal Violet. For staining, media was aspirated and the plates containing the cells were washed 3 times with PBS slowly. Cells were fixed with 100% cold methanol for 5 minutes. After fixation, plates were washed slowly with PBS for 3 times and Crystal Violet dye was added on top and incubated for 5 mins until colonies are visible. Dye was removed and the plates were rinsed with water to remove excess staining. Number of colonies were counted using ImageJ Software (NIH Image, Bethesda, MD, USA).

**Cell viability, caspase activity assays:** Cells at 12 days post-transduction were seeded for cell viability and caspase activity experiments (5000 cells per well in a 96-well plate). Viability and caspase activity were measured 2 days later. Cell viability was detected by ATP based Cell Titer-Glo (CTG) Luminescent Cell Viability Assay (Promega, USA) according to the manufacturer's instructions using a plate reader (BioTek's Synergy H1, Winooski, VT, USA). 5.000 cells/well were seeded to 96-well plates (Corning Costar, clear bottom black side) as triplicates for each condition and incubated for 48 hrs. CTG reagent was added on top of cells (4 µl CTG reagent in 40 µl DMEM for each well) and luminescence was measured in plate reader at 560 nm after 2 minutes shaking period followed by 8 minutes incubation in dark. Viability data was normalized to control condition and t-test was performed to assess significance of viability changes among different conditions. Caspase 3/7 activity was measured by Caspase-Glo® 3/7 (Promega, USA) assays according to manufacturer's instructions, similarly.

**Annexin V/PI staining:** Annexin V staining was performed with Muse® Annexin V & Dead Cell Kit (Luminex, MCH100105) according to the manufacturer's instructions. Cells at day 12 post-transduction were seeded (200,000 cells per well in 6 well plates). Two days later, all cells (both live cells attached to the culture plate and free-floating dead cells) were centrifuged at 1200 rpm for 5 minutes and the pellets were washed in cold PBS, centrifuged and resuspended in PBS containing 500 µl 1% FBS. 75 µl of the cell suspension was collected and mixed with 75 µl of Annexin V & Dead Cell Assay (Millipore) reagent (Millipore). The mixture was incubated at room temperature in the dark for 20 minutes. Cells were analyzed by Muse Cell Analyzer (Merck, Darmstadt, Germany) at excitation 488 nm and emission 530/575 nm, with 10,000 events recorded for each sample.

**Histone extraction:** Cell pellets were washed twice with cold 1X PBS and resuspended with Triton Extraction Buffer (TEB) containing 0.5% Triton X 100 (v/v), 2 mM Phenylmethylsulfonyl Fluoride (PMSF) and 0.02% NaN3 (w/v) in 1X PBS, with a density of 107 cells/ml. Cell suspension was lysed on ice for 10 minutes while shaking gently. Following incubation, cell suspension was centrifuged for at 6,500 x g for 10 minutes at 4˚C. Supernatant was discarded and pellet was resuspended in half the volume of TEB from previous step and centrifuged as described. Pellet was resuspended in 0.2 N HCl with a density of 4x107 nuclei/ml. Acid extraction was carried out overnight at 4˚C. Later, 1 M NaOH was added as 1/5 volume of HCl and centrifuged as described. The protein concentration of the supernatant, which contains histone proteins, was determined with Pierce BCA Protein Assay Kit (Thermo Scientific, 23225).

**Western blot:** Cells were harvested, and pellets were dissolved in NP-40 lysis buffer in appropriate volume (50mM Tris Buffer pH 7.4, 250mM NaCl, 5mM EDTA, 50mM NaF, 1%NP40, 0.02% NaN3, 1mM PMSF) for 30 minutes on ice, sonicated and centrifuged at 10,000 rpm at 4 °C for 10 min. Supernatants were used for protein concentration determination by Pierce's BCA Protein Assay Kit (Thermo Scientific, 23225, USA). 30 µg protein for each sample were mixed with 4X Laemli sample buffer (Biorad, 1610747, USA) containing 10% beta-mercaptoethanol and incubated at 95°C for 10 minutes. 4-12% Mini Protean TGX Precast Gel (Bio-rad, 456-1044) was used to load the samples in 1x TGS Buffer. Transfer into PVDF membrane was performed using Bio-RadTrans-Blot® Turbo™ Transfer System (Bio-Rad, USA). Membrane was blocked for an hour at RT using 5% blocking powder dissolved in 1X TBS-T. Primary antibodies Histone H3 (Cell Signaling, 9715), H3K4me1 (Diagenode, C15410194), H3K4me2 (Cell Signaling, 9725), H3K4me3 (Cell Signaling, 9751), PARP (Abcam, ab74290, USA), ASH2L (Cell Signaling, D93F6); Cleaved Casp3 (Cell Signaling, 5A1E) were prepared 1:1000 in 1X TBS-T and membranes were incubated overnight at 4˚C. Following day, membranes were washed with 1X TBS-T three times. Secondary antibody (Li-Cor, IRDye® 800CW Goat anti-Rabbit IgG Secondary Antibody) was prepared as 1:10000 in 1X TBS-T containing 0.01% SDS and membranes were incubated one hour at RT in the dark. Later, membranes were washed, and images were acquired using Li-Cor Odyssey® FC Imaging System (LI-COR Biosciences; Lincoln, NE) using SuperSignal™ West Femto Maximum Sensitivity Substrate (Thermo Scientific, 34095, USA).

**Cell cycle:** Cells were collected 14 days post-transduction and centrifuged at 1200 rpm for 5 minutes. Pellet was washed with 1X PBS and centrifuged as previously. Supernatant was removed only to leave around 100 µl of 1X PBS to resuspend the pellet. For fixation, resuspended cells were added to cold 70% Ethanol (1ml/$10^6$cells) drop by drop while vortexing. Fixed samples were kept in -20˚C for 3 to 24 hours. Fixed samples were centrifuged as described previously and washed with 1X DPBS. Cell pellets were resuspended in 150 µL of Muse Cell Cycle Reagent (The Muse® Cell Cycle Kit, Luminex,

MCH100106) and incubated for 30 minutes RT in the dark. 10,000 events/sample was measured with Muse Cell Analyzer (Merck, Darmstadt, Germany) and analyzed by Muse Cell Analyzer software.

**Quantitative RT-PCR:** 1000 ng total RNA per sample was reverse transcribed with random hexamers using M-MLV Reverse transcriptase (Invitrogen). cDNAs were used for PCR using SYBR-Green Master Mix (Roche) in a Lightcycler 480 Instrument II (Roche). 30 sec at 95 $^{o}$C, 45 sec at 50 $^{o}$C and 30 sec at 72 $^{o}$C thermal cycler conditions were repeated for 40 cycles. GAPDH was used as endogenous control for relative quantification with 2-DeltaCT method. List of primers can be found in **Supplementary Table 4.**

**TCGA:** Gene expression profiles of “glioblastoma multiforme” (GBM) and “brain lower grade glioma” (LGG) tumors were preprocessed by the unified RNA-Seq pipeline of The Cancer Genome Atlas (TCGA) consortium. For both cancer types, HTSeq-FPKM files of all primary tumors from the most recent data freeze (i.e., Data Release 32–March 29, 2022) were downloaded. Clinical annotation files of cancer patients were used to extract their tumor characteristics (i.e. neoplasm_histologic_grade). Clinical Supplement files of all patients from the most recent data freeze were downloaded. The gene expression profiles of primary tumors were first log2-transformed within each cohort before further analysis. There were 248, 261, and 156 primary tumors in LGG (G2), LGG (G3), and GBM groups, respectively. The log2-transformed gene expression values of these three groups were compared against each other using the Wilcoxon rank sum test. There were 38, 50, 26, and 30 primary tumors in Classical, Mesenchymal, Neural, and Proneural groups, respectively. The log2-transformed gene expression values of Classical, Neural, and Proneural groups were compared against those of Mesenchymal group using the Wilcoxon rank sum test.

**Immunohistochemistry:** To determine the tissue expression of ASH2L, Brain Glioblastoma tissue microarray (TMA) was purchased from US Biomax (GL806f). TMA slides were deparaffinized followed by antigen retrieval using Tris-based solution (CC1, Ventana) for 64 min at 95$^{o}$C. ASH2L (rabbit monoclonal, 5019, Cell Signaling, 1:300) was used for primary antibody staining in SignalStain Antibody Diluent (Cell Signaling) at room temperature for 16 minutes. Later, the TMA slides were counterstained and mounted. Aperio AT2 Scanner (Leica Biosystems) was used for imaging the TMA slides and Aperio ImageScope (Leica Biosystems) program with Positive Pixel Count Algorithm was used for digital scoring. Glioblastoma and normal brain cores were compared based on IHC score for ASH2L expression levels and unpaired t-test was used for statistical analysis.

**RNA sequencing:** U373 cell line was transduced with either *ASH2L* or NT sgRNAs and puromycin selection was carried out for three days. Cell pellets were collected as triplicates at 14 days post transduction for RNA isolation. Total RNAs were isolated by using MN Nucleospin RNA isolation kit according to manufacturer’s instructions. Library preparation and sequencing was performed at University of Oxford (Oxford, UK). Briefly, RNA was DNase I-treated, cleaned and concentrated (Zymo RNA Clean and Concentrator, Zymo Research) then enriched for poly(A) mRNA (NEBNext poly(A) mRNA Magnetic Isolation Module, NEB Biosystems, Ipswich, UK). Sequencing libraries were prepared using the NEBNext Ultra II RNA Library Prep kit (New England Biolabs). RNA quality was assessed using High Sensitivity RNA Screentape and measured with an Agilent 4200 tapestation. Single-indexed and multiplexed samples were run on an Illumina Next Seq 500 sequencer using a NextSeq 500 v2 kit (FC-404–2005; Illumina, Can Diago, CA) for paired-end sequencing (length was 42 bp). The sequencing data for each sample was aligned to the GENCODE GRCh38[11] human reference transcriptome using STAR[12] (Dobin *et al.* 2013), Salmon[13] was used to quantify the transcriptome BAM files based on the transcripts per million (TPM) method. The R package *tximport*[14] was used to aggregate the transcript-level abundance for each sample to the gene-level. The package DESeq2[15] was used to normalize and analyse the count data to identify differentially expressed genes between ASH2L and NTU samples. Genes that have a false discovery rate (FDR) adjusted p-value < 0.05 and either a $log_2$ fold change > 1 (up-regulated) or $log_2$ fold change < -1 (down-regulated) were highlighted as significantly differentially expressed. All plots used to visualise the gene expression analysis results were generated using *ggplot2*. Gene Set Enrichment Analysis[16] was performed using a rank-ordered gene list of all genes within the *fgsea* R package to identify pathways from the complete list of curated gene-sets (Gene Ontology, Reactome and Cancer specific pathways) in the MsigDB. Significant pathways were defined with an FDR < 0.05 as outlined in the guidelines for small sample size comparisons and to account for multiple testing[16]. Data are deposited to the NCBI GEO database with the accession number GSE201657.

**Quantitative mass spectrometry of the ASH2L interactome:** GFP-affinity purification, sample preparation and data analysis were performed as reported[17,18]. Briefly, glioblastoma cells harboring ASH2L-GFP cDNAs were grown in 15-cm dishes until subconfluency (approximately 300 million cells) and induced with 1 mg/mL doxycycline for 24h. Cells were harvested by dislodging with trypsin and pellets were washed with cold PBS (Gibco, #10010-015). Cell pellets were re-suspended in 5 packed-cell volumes (PCVs) of ice-cold Buffer A (10 mM Hepes-KOH pH 7.9, 1.5 mM MgCl2, 10 mM KCl), incubated for 10 min on ice and then centrifuged at 400 g and 4°C for 5 min. Supernatants were aspirated and cells were lysed in 2 PCVs Buffer A containing 1x CPI (Roche, #11836145001), 0.5 mM DTT and 0.15 % NP40. The suspension was homogenized in Dounce homogenizer followed by centrifugation at 3,200 g and 4°C for 15 min. Supernatant and pellet contained cytoplasmic and nuclear fractions, respectively. The nuclear pellet was washed gently with 10 volumes of Buffer A containing 1x CPI (Roche, #11836145001), 0.5 mM DTT and 0.15 % NP40 and centrifuged for 5 min at 3,200 g at 4°C min. Nuclear proteins were extracted by 2 PCV volumes of high salt Buffer B (420 mM NaCl, 20 mM Hepes-KOH pH 7.9, 20% v/v glycerol, 2 mM MgCl2, 0.2 mM EDTA, 0.1 % NP40, 1x CPI, 0.5 mM DTT) during gentle agitation at 4°C for 1.5 h. Both the nuclear and cytoplasmic extracts were centrifuged at 3,200 g and 4°C for 60 min. Supernatants were collected and protein concentration was measured by Bradford assay. 1 mg of nuclear or 1 mg of cytoplasmic extract was used for GFP-affinity purification as described[19]. In short, protein lysates were incubated in binding buffer (20 mM Hepes-KOH pH 7.9, 300 mM NaCl, 20% glycerol, 2 mM MgCl2, 0.2 mM EDTA, 0.1% NP-40, 0.5 mM DTT and 1x Roche protease inhibitor cocktail) on a rotating wheel at 4˚ C for 1 h in triplicates with GFP-Trap agarose beads (#gta-200, Chromotek) or control agarose beads (Chromotek). The beads were washed two times with binding buffer containing 0.5% NP-40, two times with PBS containing 0.5% NP-40, and two times with PBS. On-bead digestion of bound proteins was performed overnight in elution buffer (100 mM Tris-HCl pH 7.5, 2 M urea, 10 mM DTT) with 0.1 µg/ml of trypsin at RT and eluted tryptic peptides were bound to C18 stage tips (ThermoFisher, USA) prior to mass spectrometry analysis. Tryptic peptides were eluted from the C18 stage tips in H2O:acetonitrile (35:65) and dried prior to resuspension in 10 % formic acid. A third of this elution was analyzed by nanoflow-LC-MS/MS with a high performance nanoflow-HPLC Orbitrap based ms/ms system (Thermo Fisher Scientific). The analysis time was 90 min. The flow rate was 300nl/min, buffer A was 0.1% (v/v) formic acid and buffer B was 0.1% formic acid on 80% acetonitrile. A gradient of increasing organic proportion was used with a C18 separation column (2 µm particle size, 100 Å pore size, 25 cm length, 50 µm i.d., Thermo Fisher Scientific). Blank samples consisting of 10% formic acid were run for 45 min between all triplicate samples to avoid carry-over between runs. The raw data files were analyzed with MaxQuant software (version 1.5.3.30) using Uniprot human FASTA database[19,20]. Label-free quantification values (LFQ) and match between run options were selected. Intensity based absolute quantification (iBAQ) algorithm was also activated for subsequent relative protein abundance estimation[21].The obtained protein files were analyzed by Perseus software (MQ package, version 1.6.12), in which contaminants and reverse hits were filtered out[20]. Protein identification based on non-unique peptides as well as proteins identified by only one peptide in the different triplicates were excluded to increase protein prediction accuracy. For identification of the bait interactors, LFQ intensity-based values were transformed on the logarithmic scale (log2) to generate Gaussian distribution of the data. This allowed for imputation of missing values based on the normal distribution of the overall data (in Perseus, width = 0.3; shift = 1.8). The normalized LFQ intensities were compared between grouped GFP triplicates and non-GFP triplicates, using 1% as the permutation-based false discovery rate (FDR) in a two-tailed t-test. The threshold for significance (S0), based on the FDR and the ratio between GFP and non-GFP samples was kept at the constant value of 2. Relative abundance plots were obtained by comparison of the iBAQ values of GFP interactors. The values of the non-GFP iBAQ values were subtracted from the corresponding proteins in the GFP pull-down and were next normalized on the ASH2L-GFP bait protein for scaling and data representation purposes. All mass spectrometry data have been deposited to the ProteomeXchange Consortium via the PRIDE partner repository under the dataset identifier PXD033358.

**Genome localization experiments by greenCUT&RUN:** Genome localization analysis of GFP-tagged ASH2L was performed by greenCUT&RUN with the combination of enhancer-MNase and LaG16-MNase, as described[22]. Mononucleosomal Drosophila DNA was used as spike-in DNA for normalization purposes and sequencing libraries were prepared as described[17]. Briefly, purified DNA fragments were subjected to library preparation with NEB Next Ultra II and NEB Multiplex Oligo Set I/II as per manufacturer (New England Biolabs) protocol without size selection. For each library, DNA concentration was determined using a Qubit instrument (Invitrogen, USA) and size distribution was analyzed with Agilent Bioanalyzer chips (DNA high sensitivity assay). The 100-nucleotide paired-end sequencing reads were generated (Illumina, HiSeq 3000) with 6-20 Million reads per sample. greenCUT&RUN sequencing datasets have been deposited to the Sequence Read Archive (SRA) portal of the NCBI with accession ID PRJNA828380.

**Bioinformatic analyses of NGS data:** The datasets generated in-house were initially passed through quality control using Trim-galore (version 0.6.6). Further reads were aligned on the human (version hg38) and Drosophila reference genome (BDFP5) using bowtie2 (version 2.3.5.1) with option: –dovetail –local –very-sensitive-local –no-unal –no-mixed –no-discordant -I 10 -X 700[17,23]. Peaks were called using HOMER with default parameters except filtering based on clonal signals was disabled using option: -C 0. Both narrow and broad peaks were called with HOMER and merged using bedtools (version 2.27.1). Reads of the control samples were normalized with SpikeIn and used to generate TagDirectories by HOMER using option: -totalReads before peak calling. To calculate normalized total reads, ratio of the SpikeIn per human reads of control and experiment were calculated and multiplied with total number of control reads. The peaks of ASH2L overlapping with differentially expressed genes from RNAseq were identified using bedtools. Heatmaps were generated using deeptools (version 3.3.2) with default parameters. To generate the bamcompare files for the experiments, coverage was normalized based on the ratio of the SpikeIn reads (per human reads) present in experiment and controls. All next-generation sequencing datasets have been deposited to the Sequence Read Archive (SRA) portal of the NCBI with accession ID PRJNA828380.

**In vivo tumor growth**: All *in vivo* experiments were approved by the institutional ethical committee of Koç University. U87MG cells were infected with Firefly luciferace (Fluc)-mCherry (FMC) viruses. 6-8-week-old non-obese diabetic/severe combined immunodeficiency (NOD/SCID) mice were used in experiments. Control and ASH2L knocked-out cells were implanted intracranially using stereotaxic injection [from bregma, AP: -2 mm, ML: 1.5 mm, V (from dura): 2 mm; n=5/group] as described[24]. Tumors were monitored using noninvasive bioluminescence imaging IVIS Lumina III (Perkin Elmer, USA) following intraperitoneal 150 ug/g body weight of D-Luciferin injection. 34 days after treatment, mice were sacrificed. Quantification of tumor progression was performed with GraphPad PRISM software (San Diego, CA, USA).

| EPIKOL ID | Gene Name_Domain_gRNA# | gRNA Sequence |
|---|---|---|
| epikol_00001 | ACAT1_AT_1 | GCTTGGTTCCATTGCAATTC |
| epikol_00002 | ACAT1_AT_2 | AGTGCTACAAGAACACCCAT |
| epikol_00003 | ACAT1_AT_3 | TGCAATGGAACCAAGCTTAG |
| epikol_00004 | ACAT1_AT_4 | ATGGAACCAAGCTTAGTGGC |
| epikol_00005 | ACAT1_AT_5 | CTGCCTAAAAAAGATCCAAT |
| epikol_00006 | ACTL6B_1 | TCAGTCCGCGCTGGGTACGC |
| epikol_00007 | ACTL6B_2 | CAGTCCGCGCTGGGTACGCT |
| epikol_00008 | ACTL6B_3 | GGCTCCTTCTCAGTCCGCGC |
| epikol_00009 | ACTL6B_4 | GCTCCTTCTCAGTCCGCGCT |
| epikol_00010 | ACTL6B_5 | AGTCCGCGCTGGGTACGCTG |
| epikol_00011 | ACTR_PAS_1 | TGTCTGGTAATAAAGCTCTC |
| epikol_00012 | ACTR_PAS_2 | GCTTTGAAGATATAATCCGA |
| epikol_00013 | ACTR_PAS_3 | CGGATTATATCTTCAAAGCC |
| epikol_00014 | ACTR_PAS_4 | TTCTTGATAGTGACGTTTCT |
| epikol_00015 | ACTR_PAS_5 | GTCTAAATGATGGGCAGTCA |
| epikol_00016 | ASH1_BROMO_1 | CTTTGAAGATCTGGGCTAGG |
| epikol_00017 | ASH1_BROMO_2 | TTTCTTTGAAGATCTGGGCT |
| epikol_00018 | ASH1_BROMO_3 | CACAAATTTCTTTGAAGATC |
| epikol_00019 | ASH1_BROMO_4 | ACAAATTTCTTTGAAGATCT |
| epikol_00020 | ASH1_BROMO_5 | TTTGAAGATCTGGGCTAGGC |
| epikol_00021 | ASH1_SET_1 | TCTAGAACGATTTCGAGCTG |
| epikol_00022 | ASH1_SET_2 | AGGGGAGGTCGTCAGTGAAC |
| epikol_00023 | ASH1_SET_3 | GTTCACTGACGACCTCCCCT |
| epikol_00024 | ASH1_SET_4 | CATCATTGAATACCTAGGGG |
| epikol_00025 | ASH1_SET_5 | GTTCATCATTGAATACCTAG |
| epikol_00026 | ASH2L_PHD_1 | TGGTTCACGGCTGACACATT |
| epikol_00027 | ASH2L_PHD_2 | GAAGAGAATGGCCGACAGTT |
| epikol_00028 | ASH2L_PHD_3 | TGAAGAGAATGGCCGACAGT |
| epikol_00029 | ASH2L_PHD_4 | GAATGGCCGACAGTTGGGTG |
| epikol_00030 | ASH2L_PHD_5 | GCTCTACCTCACCCAACTGT |
| epikol_00031 | ASH2L_SPRY_1 | GCCTCTCATGGAGTACGGAA |
| epikol_00032 | ASH2L_SPRY_2 | CCCAGTCTGGCAGCGGTATC |
| epikol_00033 | ASH2L_SPRY_3 | ACCTTTCCGTACTCCATGAG |
| epikol_00034 | ASH2L_SPRY_4 | CTCAGATGACCGGCTGACTG |
| epikol_00035 | ASH2L_SPRY_5 | AGTCTGGCAGCGGTATCTGG |
| epikol_00036 | ASXL1_PHD_1 | CTTTCAGGCTGCACGCACAC |
| epikol_00037 | ASXL1_PHD_2 | TTCTGTCACGATGACTGTAT |
| epikol_00038 | ASXL1_PHD_3 | GTTCACTGACAGCAGCACGG |
| epikol_00039 | ASXL1_PHD_4 | GACAGAACGCACCGCAGCCT |
| epikol_00040 | ASXL1_PHD_5 | AAAGCCATGATCATGTGCCA |
| epikol_00041 | ASXL2_PHD_1 | TCTTCGGAGAACGCCTGAAC |
| epikol_00042 | ASXL2_PHD_2 | ACGCCTGAACAGGAATGGTC |
| epikol_00043 | ASXL2_PHD_3 | AGGGGCCGATGCAATCATCA |
| epikol_00044 | ASXL2_PHD_4 | GCAGGAGACGCACAGTTTGG |

| | | |
|---|---|---|
| epikol_00045 | ASXL2_PHD_5 | AAGCGGCAGTAACATTTCGA |
| epikol_00046 | ASXL3_PHD_1 | AAGGACCTATGCAGTCGTCA |
| epikol_00047 | ASXL3_PHD_2 | TTCTGCCATGACGACTGCAT |
| epikol_00048 | ASXL3_PHD_3 | CTGTCGGCAAAGTTCTGAAC |
| epikol_00049 | ASXL3_PHD_4 | TATGCAGTCGTCATGGCAGA |
| epikol_00050 | ASXL3_PHD_5 | AATCCTGCCACAGGCTTGTC |
| epikol_00051 | ATAD2_BROMO_1 | CAACAGGCTTAGTAAACACT |
| epikol_00052 | ATAD2_BROMO_2 | TCTTAAGAAATGTTACACAT |
| epikol_00053 | ATAD2_BROMO_3 | TACAATCCAGATAGAGATCC |
| epikol_00054 | ATAD2_BROMO_4 | ATTACAGATGAAAGGTCCAT |
| epikol_00055 | ATAD2_BROMO_5 | TCTATCTGGATTGTATTCTA |
| epikol_00056 | ATAD2B_BROMO_1 | TAACATCTTCAGCAAACCGG |
| epikol_00057 | ATAD2B_BROMO_2 | GAGAGTTGCGGTTGTTTCTC |
| epikol_00058 | ATAD2B_BROMO_3 | TTCTCAGGGATGTAACCAAG |
| epikol_00059 | ATAD2B_BROMO_4 | CAGGGATGTAACCAAGAGGC |
| epikol_00060 | ATAD2B_BROMO_5 | GATGTTAAAGCGTTTATCTG |
| epikol_00061 | ATRX_HELIC_1 | TGAAATTCTTCGAATGGCAG |
| epikol_00062 | ATRX_HELIC_2 | TTCGAATGGCAGAGGAAATT |
| epikol_00063 | ATRX_HELIC_3 | CTTCGAATGGCAGAGGAAAT |
| epikol_00064 | ATRX_HELIC_4 | TCTCTTTGAAATTCTTCGAA |
| epikol_00065 | ATRX_HELIC_5 | TCGAATGGCAGAGGAAATTG |
| epikol_00066 | BAZ1A_BROMO_1 | CGAAGGAGTTCTGGCCGACA |
| epikol_00067 | BAZ1A_BROMO_2 | TGGTACGACATGATGACAGC |
| epikol_00068 | BAZ1A_BROMO_3 | AGGAGTTCTGGCCGACAGGG |
| epikol_00069 | BAZ1A_BROMO_4 | ATTCATGAACTCCTCCCTGT |
| epikol_00070 | BAZ1A_BROMO_5 | TGAACAACTTGTTGTAGAAT |
| epikol_00071 | BAZ1A_PHD_1 | GAGCTTTGGTCGAACACAGT |
| epikol_00072 | BAZ1A_PHD_2 | TGCAAGATATGTCGAAAGAA |
| epikol_00073 | BAZ1A_PHD_3 | CTTTGTGATGGCTGTGATAG |
| epikol_00074 | BAZ1A_PHD_4 | TTCTTTGTGATGGCTGTGAT |
| epikol_00075 | BAZ1A_PHD_5 | GAAAACATGGTTCTTTGTGA |
| epikol_00076 | BAZ1B_BROMO_1 | AGCGGTACTTCACGATCTTG |
| epikol_00077 | BAZ1B_BROMO_2 | TCGTGAAGTACCGCTTCAGC |
| epikol_00078 | BAZ1B_BROMO_3 | GGTACTTCACGATCTTGTGG |
| epikol_00079 | BAZ1B_BROMO_4 | GCTGAGGTTTACAACTGCCG |
| epikol_00080 | BAZ1B_BROMO_5 | TGATGTGATCACGCACCCCA |
| epikol_00081 | BAZ2A_BROMO_1 | CTCACCAAACGTGGGTTCAC |
| epikol_00082 | BAZ2A_BROMO_2 | CGGTACCCACTCACCAAACG |
| epikol_00083 | BAZ2A_BROMO_3 | TGAACCCACGTTTGGTGAGT |
| epikol_00084 | BAZ2A_BROMO_4 | GGTACCCACTCACCAAACGT |
| epikol_00085 | BAZ2A_BROMO_5 | GTGAACCCACGTTTGGTGAG |
| epikol_00086 | BAZ2A_MBD_1 | CCTGGTATTATGGCCCCTGT |
| epikol_00087 | BAZ2A_MBD_2 | CCCACAGGGGCCATAATACC |
| epikol_00088 | BAZ2A_MBD_3 | AGGTCTCCCCCTGCCATCGG |
| epikol_00089 | BAZ2A_MBD_4 | GAGAGAGGTGCGCATCAAGA |
| epikol_00090 | BAZ2A_MBD_5 | GAAGGGCAGCCACCGATGGC |
| epikol_00091 | BAZ2B_BROMO_1 | GTAAACTTGAAACTTGTTCC |
| epikol_00092 | BAZ2B_BROMO_2 | TGGAAACTCATGAGGATGCA |

| | | |
|---|---|---|
| epikol_00093 | BAZ2B_BROMO_3 | GGAACAAGTTTCAAGTTTAC |
| epikol_00094 | BAZ2B_BROMO_4 | CTAATTGTGGAAAAATCCAT |
| epikol_00095 | BAZ2B_BROMO_5 | ACTTAGTTTCTCTCTAATTG |
| epikol_00096 | BAZ2B_HMBD_1 | AACTTTGGAGGGCGCCTTCA |
| epikol_00097 | BAZ2B_HMBD_2 | GTAGCATATTATGCTCCATG |
| epikol_00098 | BAZ2B_HMBD_3 | AATATGCTACTTCTCCTTGA |
| epikol_00099 | BAZ2B_HMBD_4 | CTCCATGTGGAAAGAAACTT |
| epikol_00100 | BAZ2B_HMBD_5 | TGCCTAAGTTTCTTTCCACA |
| epikol_00101 | BMI_RING_1 | TAGACATTCTATTATGGTTG |
| epikol_00102 | BMI_RING_2 | GGAATGTAGACATTCTATTA |
| epikol_00103 | BMI_RING_3 | AACGTGTATTGTTCGTTACC |
| epikol_00104 | BMI_RING_4 | GACAATACTTGCTGGTCTCC |
| epikol_00105 | BMI_RING_5 | ACAAATAGGACAATACTTGC |
| epikol_00106 | BPTF_BROMO_1 | GGATTATGAGGGGTTGAAGA |
| epikol_00107 | BPTF_BROMO_2 | AGGATTATGAGGGGTTGAAG |
| epikol_00108 | BPTF_BROMO_3 | GGTGCATCATTAGGGTCTAC |
| epikol_00109 | BPTF_BROMO_4 | CATTAGGGTCTACTGGTTCA |
| epikol_00110 | BPTF_BROMO_5 | GAAAGGCCAGGCCATCTTAT |
| epikol_00111 | BPTF_GECKO_3 | GTTCCGCAGTACCTCGTAAA |
| epikol_00112 | BPTF_GECKO_4 | ACGCTATCTTTCAGATCAGC |
| epikol_00113 | BPTF_GECKO_5 | ACCATTAGCTGCCGCCAGAA |
| epikol_00114 | BPTF_PHD_1 | TGGTACCATGGGCGCTGCGT |
| epikol_00115 | BPTF_PHD_2 | AGATGCCAACGCAGCGCCCA |
| epikol_00116 | BRD1_BROMO_1 | GCTGCGCAAATATCCTGGCG |
| epikol_00117 | BRD1_BROMO_2 | TGCGCAGCCCGTGAGTCTGA |
| epikol_00118 | BRD1_BROMO_3 | GGGCTGCGCAAATATCCTGG |
| epikol_00119 | BRD1_BROMO_4 | TGCAAGACAAGGACCCCGCC |
| epikol_00120 | BRD1_BROMO_5 | GAGCGCAGCAGCACCGTCAG |
| epikol_00121 | BRD1_PHD_1 | AAAAGACAGATGACGACCGC |
| epikol_00122 | BRD1_PHD_2 | GTGTTCATCGAGCCCATCGA |
| epikol_00123 | BRD1_PHD_3 | GGGCTCGATGAACACCGTGT |
| epikol_00124 | BRD1_PHD_4 | AAAGACAGATGACGACCGCT |
| epikol_00125 | BRD1_PHD_5 | TGTTCATCGAGCCCATCGAT |
| epikol_00126 | BRD2_BROMO_1 | ATCAGTTCGCATGGCCATTC |
| epikol_00127 | BRD2_BROMO_2 | TGTGGAAACATCAGTTCGCA |
| epikol_00128 | BRD2_BROMO_3 | CCTTGTGTAGGTATTGCAGC |
| epikol_00129 | BRD2_BROMO_4 | GCCTGTGGATGCTGTCAAAC |
| epikol_00130 | BRD2_BROMO_5 | CCTGTGGATGCTGTCAAACT |
| epikol_00131 | BRD3_BROMO_1 | ATTTGATTGCGTCCACGGGC |
| epikol_00132 | BRD3_BROMO_2 | TTCAATTTGATTGCGTCCAC |
| epikol_00133 | BRD3_BROMO_3 | GTTCAATTTGATTGCGTCCA |
| epikol_00134 | BRD3_BROMO_4 | TCTGGAAACACCAGTTCGCC |
| epikol_00135 | BRD3_BROMO_5 | TGCGTCCACGGGCTGGTAGA |
| epikol_00136 | BRD4_BROMO_1 | TTCAGCTTGACGGCATCCAC |
| epikol_00137 | BRD4_BROMO_2 | GAGTGGTGCTCAAGACACTA |
| epikol_00138 | BRD4_BROMO_3 | GCTGGAAAGGCCATGCAAAC |
| epikol_00139 | BRD4_BROMO_4 | CTCTGAGCAGGTATTGCAGT |
| epikol_00140 | BRD4_BROMO_5 | GAGCAGGTATTGCAGTTGGT |

| | | |
|---|---|---|
| epikol_00141 | BRD7_GECKO_1 | AACTGATGAGACAATTGCAG |
| epikol_00142 | BRD7_GECKO_2 | GTTATTCCAACCTTCTTGTT |
| epikol_00143 | BRD7_GECKO_3 | AGCTCTTTAGCCAAACAAGA |
| epikol_00144 | BRD7_GECKO_4 | AGTTACCTTTAGTTCTTCTA |
| epikol_00145 | BRD7_GECKO_5 | GTTGATTCAAAGCTTCTTGA |
| epikol_00146 | BRD8_BROMO_1 | AGCTCTCCATACAAGCATGA |
| epikol_00147 | BRD8_BROMO_2 | GTTACAGATGACATAGCACC |
| epikol_00148 | BRD8_BROMO_3 | GGTGCTATGTCATCTGTAAC |
| epikol_00149 | BRD8_BROMO_4 | TGTCATCTGTAACAGGCTGC |
| epikol_00150 | BRD8_BROMO_5 | TGCACAATGCTGTGGTAGCC |
| epikol_00151 | BRD9_BROMO_1 | GCTGGCGGAGGAAGTGTTCC |
| epikol_00152 | BRD9_BROMO_2 | TGTTCCAGGAGTTGCTGAAT |
| epikol_00153 | BRD9_BROMO_3 | GGAGCAATTGCATCCGTGAC |
| epikol_00154 | BRD9_BROMO_4 | GTCACGGATGCAATTGCTCC |
| epikol_00155 | BRD9_BROMO_5 | TTCATGGTGCCAAAATCCAT |
| epikol_00156 | BRDT_BROMO_1 | ATGGCCCTTTCAACGTCCTG |
| epikol_00157 | BRDT_BROMO_2 | GCATCCACAGGACGTTGAAA |
| epikol_00158 | BRDT_BROMO_3 | AGCATCCACAGGACGTTGAA |
| epikol_00159 | BRDT_BROMO_4 | TGTAGTTTCACAGCATCCAC |
| epikol_00160 | BRDT_BROMO_5 | AAGTTGTCCTAAAGGATTTA |
| epikol_00161 | BRPF1_BROMO_1 | CTGCCTCAAGTATAACGCCA |
| epikol_00162 | BRPF1_BROMO_2 | TCTGTCTGAGGTAACCGAAT |
| epikol_00163 | BRPF1_BROMO_3 | TTGATGTGGTCTAGGTAGTC |
| epikol_00164 | BRPF1_BROMO_4 | AGGCAACATCTTCAGCGAGC |
| epikol_00165 | BRPF1_BROMO_5 | CATCCTCCTTCGCAAAACCT |
| epikol_00166 | BRPF1_PHD_1 | AGCAGACAGATGACGGGCGC |
| epikol_00167 | BRPF1_PHD_2 | GCAGACAGATGACGGGCGCT |
| epikol_00168 | BRPF1_PHD_3 | ACCGCCCTTGTTGGGGCACA |
| epikol_00169 | BRPF1_PHD_4 | TTTGCAAACAACGGGGCTCA |
| epikol_00170 | BRPF1_PHD_5 | ATTTGCAAACAACGGGGCTC |
| epikol_00171 | BRPF3_BROMO_1 | TGTTCTGTTGAGGACAACAC |
| epikol_00172 | BRPF3_BROMO_2 | GACAACACTGGACCTGCTGC |
| epikol_00173 | BRPF3_BROMO_3 | GTCCTCAACAGAACATTGAA |
| epikol_00174 | BRPF3_BROMO_4 | CAGGATCCTTCTCCTGCAGC |
| epikol_00175 | BRPF3_BROMO_5 | ACTGGACCTGCTGCAGGAGA |
| epikol_00176 | BRPF3_PHD_1 | TTGAGGCTGGTTTCGCGCAT |
| epikol_00177 | BRPF3_PHD_2 | ATTGAGGCTGGTTTCGCGCA |
| epikol_00178 | BRPF3_PHD_3 | TTGAAGGCGCCACCCTTATT |
| epikol_00179 | BRPF3_PHD_4 | ACAATATCCCGCCTGCCCGC |
| epikol_00180 | BRPF3_PHD_5 | CGCCACCCTTATTGGGGCAA |
| epikol_00181 | BRWD1_BROMO_1 | GGATATTCAACCAAATCAAC |
| epikol_00182 | BRWD1_BROMO_2 | ATTTAGACAACCTGTTGATT |
| epikol_00183 | BRWD1_BROMO_3 | AAATCAACAGGTTGTCTAAA |
| epikol_00184 | BRWD1_BROMO_4 | CAACTGGAAGAAACAGTGTA |
| epikol_00185 | BRWD1_BROMO_5 | GTAAGGGAAACTCTAGATGC |
| epikol_00186 | BRWD3_BROMO_1 | AGAATGTGAACGGGTTATTC |
| epikol_00187 | BRWD3_BROMO_2 | CCGTTCACATTCTTCGTCTC |
| epikol_00188 | BRWD3_BROMO_3 | GAATGTGAACGGGTTATTCA |

| | | |
|---|---|---|
| epikol_00189 | BRWD3_BROMO_4 | CAGAGACGAAGAATGTGAAC |
| epikol_00190 | BRWD3_BROMO_5 | CAAACCCCAGGAAGGAGAGT |
| epikol_00191 | CARM1_MTA_1 | AGCACGGAAAATCTACGCGG |
| epikol_00192 | CARM1_MTA_2 | ACGGAAAATCTACGCGGTGG |
| epikol_00193 | CARM1_MTA_3 | CTTGGGCGGCAAAAAACGAC |
| epikol_00194 | CARM1_MTA_4 | TGGAGCACGGAAAATCTACG |
| epikol_00195 | CARM1_MTA_5 | TGACCACGATGCGGTCCGTC |
| epikol_00196 | CBX1_GECKO_1 | GGCCGACTCACCATTTCATC |
| epikol_00197 | CBX1_GECKO_2 | AAAAGTTCTCGACCGTCGAG |
| epikol_00198 | CBX1_GECKO_3 | CTTTGCCCTTTACCACTCGA |
| epikol_00199 | CBX1_GECKO_4 | CTGCAGGAAAAACTCTGATG |
| epikol_00200 | CBX1_GECKO_5 | TTACAGTCAGAAAAGCCACG |
| epikol_00201 | CBX2_CHROM_1 | ACCTGGTCAAGTGGCGCGGC |
| epikol_00202 | CBX2_CHROM_2 | GAGTACCTGGTCAAGTGGCG |
| epikol_00203 | CBX2_CHROM_3 | ACCAGCCGCGCCACTTGACC |
| epikol_00204 | CBX2_CHROM_4 | GGAAGGCCAGGAGCAGCCTC |
| epikol_00205 | CBX2_CHROM_5 | TGGAAGGCCAGGAGCAGCCT |
| epikol_00206 | CBX3_GECKO_1 | AACACAGTGCTGACAATACT |
| epikol_00207 | CBX3_GECKO_2 | AGACTTGGTGCTGGCGAAAG |
| epikol_00208 | CBX3_GECKO_3 | AAACAGGCTGACAAACCAAG |
| epikol_00209 | CBX3_GECKO_4 | AGAAACGCTTCAATCAATTC |
| epikol_00210 | CBX3_GECKO_5 | AAACAGGAAAGATTCAGATG |
| epikol_00211 | CBX4_CHROM_1 | GAGTATCTGGTGAAATGGAG |
| epikol_00212 | CBX4_CHROM_2 | ATCTGGTGAAATGGAGAGGC |
| epikol_00213 | CBX4_CHROM_3 | TGCTGATCGCCTTCCAGAAC |
| epikol_00214 | CBX4_CHROM_4 | TGGAAGGCGATCAGCAGCCT |
| epikol_00215 | CBX4_CHROM_5 | GGAAGGCGATCAGCAGCCTG |
| epikol_00216 | CBX5_CHROM_1 | CTAGACAGGCGCGTGGTTAA |
| epikol_00217 | CBX5_CHROM_2 | GCTAGACAGGCGCGTGGTTA |
| epikol_00218 | CBX5_CHROM_3 | GAAGGTGCTAGACAGGCGCG |
| epikol_00219 | CBX5_CHROM_4 | GCGCGTGGTTAAGGGACAAG |
| epikol_00220 | CBX5_CHROM_5 | AAGTGGAATATCTACTGAAG |
| epikol_00221 | CBX6_CHROM_1 | TCGGATCCGCCGTTTGATGA |
| epikol_00222 | CBX6_CHROM_2 | CCGCCGTTTGATGATGGATT |
| epikol_00223 | CBX6_CHROM_3 | CCGAATCCATCATCAAACGG |
| epikol_00224 | CBX6_CHROM_4 | CGGCCGAATCCATCATCAAA |
| epikol_00225 | CBX6_CHROM_5 | GCATCGAGTACCTGGTGAAA |
| epikol_00226 | CBX7_CHROM_1 | GCATCCGGAAGAAGCGCGTG |
| epikol_00227 | CBX7_CHROM_2 | AAGTCGAGTATCTGGTGAAG |
| epikol_00228 | CBX7_CHROM_3 | ATCTGGTGAAGTGGAAAGGA |
| epikol_00229 | CBX7_CHROM_4 | TCGTAGGCCATGACGAGGCG |
| epikol_00230 | CBX7_CHROM_5 | TCTCCTCGTAGGCCATGACG |
| epikol_00231 | CBX8_CHROM_1 | CCCTCCTGAAGCGGCGCATA |
| epikol_00232 | CBX8_CHROM_2 | TCCGTATGCGCCGCTTCAGG |
| epikol_00233 | CBX8_CHROM_3 | CCGTATGCGCCGCTTCAGGA |
| epikol_00234 | CBX8_CHROM_4 | CGGCCGAAGCCCTCCTGAAG |
| epikol_00235 | CBX8_CHROM_5 | GCGCCGCTTCAGGAGGGCTT |
| epikol_00236 | CDY1_GECKO_1 | TTAACATGGATTCTATCCCT |

| | | |
|---|---|---|
| epikol_00237 | CDY1_GECKO_2 | GTCTTTCCTCATGGCTTCCC |
| epikol_00238 | CDY1_GECKO_3 | CTGATCATTCTGTGTCTTTC |
| epikol_00239 | CDY1_GECKO_4 | ATATCCTAGAATACCTCTGT |
| epikol_00240 | CDY1_GECKO_5 | CTCTGTCCAAAGGTCGTATA |
| epikol_00241 | CDY1L_CHROM_1 | GTGGAAAGGCTATGACAGCG |
| epikol_00242 | CDY1L_CHROM_2 | ACAGCACCTCGTGAACTGTG |
| epikol_00243 | CDY1L_CHROM_3 | CGAGGACGACACTTGGGAGC |
| epikol_00244 | CDYL_GECKO_4 | AAAGCCGGTCGGAGCTTTAT |
| epikol_00245 | CDYL_GECKO_5 | TACGACGTCTGACAGATGAC |
| epikol_00246 | CHAF1A_1 | TCTTCCACCAGCTCGCCCGA |
| epikol_00247 | CHAF1A_2 | CTTCCACCAGCTCGCCCGAG |
| epikol_00248 | CHAF1A_3 | CTCTTCCACCAGCTCGCCCG |
| epikol_00249 | CHAF1A_4 | TGAGTACAGAGTCTTCTTCG |
| epikol_00250 | CHAF1A_5 | AGGCGGCCCCTCGGGCGAGC |
| epikol_00251 | CHAF1B_1 | ACACATCGAGGGTCTTCGCC |
| epikol_00252 | CHAF1B_2 | CCTGAGGGGGAGTCGTGCCG |
| epikol_00253 | CHAF1B_3 | AGGGGTTCCCTCTACCGGTC |
| epikol_00254 | CHAF1B_4 | GGGGTTCCCTCTACCGGTCT |
| epikol_00255 | CHAF1B_5 | GCCTGAGGGGGAGTCGTGCC |
| epikol_00256 | CHD1_CHROM_1 | GGTCCCATATCCACAACACT |
| epikol_00257 | CHD1_CHROM_2 | CTCCCAAGTGTTGTGGATAT |
| epikol_00258 | CHD1_CHROM_3 | CTCAAGCAGCAGAATGTTAG |
| epikol_00259 | CHD1_CHROM_4 | GTCCCATATCCACAACACTT |
| epikol_00260 | CHD1_CHROM_5 | TCTAACATTCTGCTGCTTGA |
| epikol_00261 | CHD1_HELIC_1 | AGGCTAGAGCCCATCGAATT |
| epikol_00262 | CHD1_HELIC_2 | TTGTTATATTTGATTCCGAT |
| epikol_00263 | CHD1_HELIC_3 | ACAGAATGATCTTCAGGCAC |
| epikol_00264 | CHD1_HELIC_4 | CCCTAGACCTCCAGCTCTTG |
| epikol_00265 | CHD1_HELIC_5 | GATCAATAAAAGGAGAACTG |
| epikol_00266 | CHD1L_HELIC_1 | ATGCTAGTAGCTTATCCAGC |
| epikol_00267 | CHD1L_HELIC_2 | CTACTAGCATTCCTGTATTC |
| epikol_00268 | CHD1L_HELIC_3 | TACTAGCATTCCTGTATTCT |
| epikol_00269 | CHD1L_HELIC_4 | CTTGGAGAATATCCAACATC |
| epikol_00270 | CHD1L_HELIC_5 | TATCCAACATCTGGGTCATT |
| epikol_00271 | CHD2_CHROM_1 | GAAGCTAATGGCGACCCTAG |
| epikol_00272 | CHD2_CHROM_2 | CCTTCCACTTGATGAGGTAC |
| epikol_00273 | CHD2_CHROM_3 | GTATATGCGATTGAAGCTAA |
| epikol_00274 | CHD2_CHROM_4 | ACCAACCCTTCCACTTGATG |
| epikol_00275 | CHD2_CHROM_5 | CAGTGTCAAAGTCACCACTA |
| epikol_00276 | CHD2_HELIC_1 | GGTGAGAATGTTGGATATCC |
| epikol_00277 | CHD2_HELIC_2 | GTTGACAAGACTTCGAGAAA |
| epikol_00278 | CHD2_HELIC_3 | TAATAGTTAGGTATTCAGCC |
| epikol_00279 | CHD2_HELIC_4 | TTGACAAGACTTCGAGAAAG |
| epikol_00280 | CHD2_HELIC_5 | TGACAAGACTTCGAGAAAGG |
| epikol_00281 | CHD3_CHROM_1 | TACCATATGACCAGTCCACG |
| epikol_00282 | CHD3_CHROM_2 | ACCATATGACCAGTCCACGT |
| epikol_00283 | CHD3_CHROM_3 | TCCCACGTGGACTGGTCATA |
| epikol_00284 | CHD3_CHROM_4 | ATATGACCAGTCCACGTGGG |

| | | |
|---|---|---|
| epikol_00285 | CHD3_CHROM_5 | CATCTTCCTCCCACGTGGAC |
| epikol_00286 | CHD3_HELIC_1 | TCAGCTTTCGCAGCATCTTC |
| epikol_00287 | CHD3_HELIC_2 | GAAGATGCTGCGAAAGCTGA |
| epikol_00288 | CHD3_HELIC_3 | CTGCGAAAGCTGAAGGAGCA |
| epikol_00289 | CHD3_HELIC_4 | TACAAGTATGAGCGCATCGA |
| epikol_00290 | CHD3_HELIC_5 | GCGCATCGATGGTGGTATCA |
| epikol_00291 | CHD4_PHD_1 | GGATGATCTCACCGCCTTGC |
| epikol_00292 | CHD4_PHD_2 | CACGGGGACAGGTATCACAC |
| epikol_00293 | CHD4_PHD_3 | CCTTCTCCATGTCGGGATCC |
| epikol_00294 | CHD4_PHD_4 | CGACATGGAGAAGGCTCCCG |
| epikol_00295 | CHD4_PHD_5 | CAGACCATGTGGTAAGCACG |
| epikol_00296 | CHD5_CHROM_1 | GTCGATGTCATCGATCTCCC |
| epikol_00297 | CHD5_CHROM_2 | TGCTTGAGGTTGTCGTAGTA |
| epikol_00298 | CHD5_CHROM_3 | TCCCAGGTGCACTGGTCGTA |
| epikol_00299 | CHD5_CHROM_4 | CTACTACGACAACCTCAAGC |
| epikol_00300 | CHD5_CHROM_5 | AGGTGCACTGGTCGTAGGGC |
| epikol_00301 | CHD5_PHD_1 | GGCCTACCATCTCGTATGCC |
| epikol_00302 | CHD5_PHD_2 | ATACGAGATGGTAGGCCCTC |
| epikol_00303 | CHD5_PHD_3 | CCCTCGGGCAGGTGTCGCAC |
| epikol_00304 | CHD5_PHD_4 | CCTGTGCGACACCTGCCCGA |
| epikol_00305 | CHD5_PHD_5 | GTCCAGGCATACGAGATGGT |
| epikol_00306 | CHD6_CHROM_1 | CTTGGAGGTGGCCCACACCA |
| epikol_00307 | CHD6_CHROM_2 | CTGTTTCTGCATCCTTGGTG |
| epikol_00308 | CHD6_CHROM_3 | TGTTTCTGCATCCTTGGTGT |
| epikol_00309 | CHD6_CHROM_4 | CACACCAAGGATGCAGAAAC |
| epikol_00310 | CHD6_CHROM_5 | ACACCAAGGATGCAGAAACA |
| epikol_00311 | CHD6_HELIC_1 | GGTAATCTTCTAGGATGTCG |
| epikol_00312 | CHD6_HELIC_2 | CTACTCCCTAAGCTGATTGC |
| epikol_00313 | CHD6_HELIC_3 | TGGCCACCTGCAATCAGCTT |
| epikol_00314 | CHD6_HELIC_4 | CTCCCTAAGCTGATTGCAGG |
| epikol_00315 | CHD6_HELIC_5 | GGCCACCTGCAATCAGCTTA |
| epikol_00316 | CHD7_CHROM_1 | GCACGTAGCACAGATGACCG |
| epikol_00317 | CHD7_CHROM_2 | TACGTGCAAAGTCCATTATC |
| epikol_00318 | CHD7_CHROM_3 | TGCACGTAGCACAGATGACC |
| epikol_00319 | CHD7_CHROM_4 | TTGCACGTAGCACAGATGAC |
| epikol_00320 | CHD7_CHROM_5 | CATAGATCAAGCAAAGATCG |
| epikol_00321 | CHD7_HELIC_1 | GGTGCGCTGCTTGGACATAC |
| epikol_00322 | CHD7_HELIC_2 | TTCCCAGATGGTGCGCTGCT |
| epikol_00323 | CHD7_HELIC_3 | AGGATCGACGGCCGAGTAAG |
| epikol_00324 | CHD7_HELIC_4 | CAAGCTGCTGCCAAAACTGA |
| epikol_00325 | CHD7_HELIC_5 | CTGCTGCCAAAACTGAAGGC |
| epikol_00326 | CHD8_CHROM_1 | GCTTCAGTATATTGTCCAGA |
| epikol_00327 | CHD8_CHROM_2 | CTTCAGTATATTGTCCAGAA |
| epikol_00328 | CHD8_CHROM_3 | TACTATCTCCCAACTAGAGA |
| epikol_00329 | CHD8_CHROM_4 | AGCCCATTCACAATGCAGAT |
| epikol_00330 | CHD8_CHROM_5 | CCTCTTATCCTTCTCTAGTT |
| epikol_00331 | CHD8_HELIC_1 | GGTTCGTTCAGCCGGCAAAC |
| epikol_00332 | CHD8_HELIC_2 | TGGCCACCAGCTTTAAGCTT |

| | | |
|---|---|---|
| epikol_00333 | CHD8_HELIC_3 | GTCAATAAGAACCAGTTTGC |
| epikol_00334 | CHD8_HELIC_4 | AATAATCCTCTAGGATGTCT |
| epikol_00335 | CHD8_HELIC_5 | TTGCTTCCAAAGCTTAAAGC |
| epikol_00336 | CHD9_GECKO_1 | TATCTAAAGCAAGAAGTTAT |
| epikol_00337 | CHD9_GECKO_2 | CTCTATATCATCTTTAAAGA |
| epikol_00338 | CHD9_CHROM_3 | GGCCCACTCACAGTGAAGAT |
| epikol_00339 | CHD9_CHROM_5 | TTTCAAAAGCTGCTCTTCTG |
| epikol_00340 | CHD9_GECKO_4 | CATCTTTAAAGATGATATAG |
| epikol_00341 | CHD9_HELIC_1 | GGTTCGTTGCCTTGACATTC |
| epikol_00342 | CHD9_HELIC_2 | CTTCCCAAAATGAAAGCCGG |
| epikol_00343 | CHD9_HELIC_3 | GATGAGCACTTTATGACCTC |
| epikol_00344 | CHD9_HELIC_4 | ATGACCTCCGGCTTTCATTT |
| epikol_00345 | CHD9_HELIC_5 | GATAGTCCTCCAGAATGTCA |
| epikol_00346 | CLOCK_PAS_1 | CATCTAGACATAGTTTAGAA |
| epikol_00347 | CLOCK_PAS_2 | GGCAAATACCCTATTATGGG |
| epikol_00348 | CLOCK_PAS_3 | AGGCTATGATTACTATCATG |
| epikol_00349 | CLOCK_PAS_4 | AATGGCAAATACCCTATTAT |
| epikol_00350 | CLOCK_PAS_5 | AAATGGCAAATACCCTATTA |
| epikol_00351 | CREBBP_BROMO_1 | CCTAGAAGCACTGTATCGAC |
| epikol_00352 | CREBBP_BROMO_2 | AGGGCCTGGCGTAACTCCTC |
| epikol_00353 | CREBBP_BROMO_3 | GCCGGAAAGGTAATGACTCT |
| epikol_00354 | CREBBP_BROMO_4 | TGCCGGAAAGGTAATGACTC |
| epikol_00355 | CREBBP_BROMO_5 | CTGTCGATACAGTGCTTCTA |
| epikol_00356 | CREBBP_KAT_1 | CTCCACCGTCTTGTCTGAGC |
| epikol_00357 | CREBBP_KAT_2 | CAAAAACCTCCCCGGCTTCA |
| epikol_00358 | CREBBP_KAT_3 | CCCCGGCTTCAGGGTGATTC |
| epikol_00359 | CREBBP_KAT_4 | GAATCACCCTGAAGCCGGGG |
| epikol_00360 | CREBBP_KAT_5 | ACAAAAACCTCCCCGGCTTC |
| epikol_00361 | CXXC1_PHD_1 | AGCAGTTGATGTCCGGTTTG |
| epikol_00362 | CXXC1_PHD_2 | CCGACAGTACCACTCCCGGA |
| epikol_00363 | CXXC1_PHD_3 | CATCCGGGAGTGGTACTGTC |
| epikol_00364 | CXXC1_PHD_4 | GTACCACTCCCGGATGGCCT |
| epikol_00365 | CXXC1_PHD_5 | CCATCCGGGAGTGGTACTGT |
| epikol_00366 | DMAP1_DMTA_1 | GCAGCGGCGCACGGAACGCA |
| epikol_00367 | DMAP1_DMTA_2 | ACAGGAGCTGCGCAAGATTG |
| epikol_00368 | DMAP1_DMTA_3 | CACTGCAGAGCAGCGGCGCA |
| epikol_00369 | DMAP1_DMTA_4 | TCTTGCGCAGCTCCTGTAGC |
| epikol_00370 | DMAP1_DMTA_5 | CGTGCGCCGCTGCTCTGCAG |
| epikol_00371 | DNMT1_DMTA_1 | GCAGCCAGAAAACACATCCA |
| epikol_00372 | DNMT1_DMTA_2 | TTCTGGCTGCGGGGGGTTGT |
| epikol_00373 | DNMT1_DMTA_3 | GGCTGCGGGGGGTTGTCGGA |
| epikol_00374 | DNMT1_DMTA_4 | AGTAGGTGCGCGAATTGAAG |
| epikol_00375 | DNMT1_DMTA_5 | CGCAGCCAGAAAACACATCC |
| epikol_00376 | DNMT3A_DMTA_1 | AAGAGAGACAGCACCCGGAT |
| epikol_00377 | DNMT3A_DMTA_2 | AAAGAGAGACAGCACCCGGA |
| epikol_00378 | DNMT3A_DMTA_3 | CGGGTGCTGTCTCTCTTTGA |
| epikol_00379 | DNMT3A_DMTA_4 | GGGGAAGATCATGTACGTCG |
| epikol_00380 | DNMT3A_DMTA_5 | GGACCGCTACATTGCCTCGG |

| | | |
|---|---|---|
| epikol_00381 | DNMT3B_DMTA_1 | AACAATGACAGGACTCGAAT |
| epikol_00382 | DNMT3B_DMTA_2 | CGATGCCATCAAACAATGAC |
| epikol_00383 | DNMT3B_DMTA_3 | CGAGTCCTGTCATTGTTTGA |
| epikol_00384 | DNMT3B_DMTA_4 | AAACAATGACAGGACTCGAA |
| epikol_00385 | DNMT3B_DMTA_5 | TCAAATACGTGAACGACGTG |
| epikol_00386 | DNMT3L_1 | CATCTGCTGCGGAAGTCTCC |
| epikol_00387 | DNMT3L_2 | AGATCCCTCCCTCAAACAGA |
| epikol_00388 | DNMT3L_3 | ACACAGCACCCTCTGTTTGA |
| epikol_00389 | DNMT3L_4 | AGCACCCTCTGTTTGAGGGA |
| epikol_00390 | DNMT3L_5 | CAGATCCCTCCCTCAAACAG |
| epikol_00391 | DOT1L_MTA_1 | AGACATCCCGGCCAAGTATG |
| epikol_00392 | DOT1L_MTA_2 | CAGCAACCTGGAGCACGACC |
| epikol_00393 | DOT1L_MTA_3 | CGTCGAGAAAGCAGACATCC |
| epikol_00394 | DOT1L_MTA_4 | TGTGGGCCAGGTCGTGCTCC |
| epikol_00395 | DOT1L_MTA_5 | GCCATAGTGATGTTTGCAGT |
| epikol_00396 | DTX3L_RING_1 | CATGGCTTTGTTGATACAAG |
| epikol_00397 | DTX3L_RING_2 | GACATGGCTTTGTTGATACA |
| epikol_00398 | DTX3L_RING_3 | GTCTGGCATGTGGGACAGAT |
| epikol_00399 | DTX3L_RING_4 | ACATGGCTTTGTTGATACAA |
| epikol_00400 | DTX3L_RING_5 | CAGAATTCATGCTTGCACTT |
| epikol_00401 | EED_WD40_1 | TTTAACTGGCACAGTAAAGA |
| epikol_00402 | EED_WD40_2 | CCGGTTGTTGCAATCTTACG |
| epikol_00403 | EED_WD40_3 | CCACGTAAGATTGCAACAAC |
| epikol_00404 | EED_WD40_4 | ATGGCTCGTATTGCTATCAT |
| epikol_00405 | EED_WD40_5 | CAATACGAGCCATCCTCTGC |
| epikol_00406 | EHMT1_SET_1 | CTCTACCGGACGCGGGACAT |
| epikol_00407 | EHMT1_SET_2 | GCTCTACCGGACGCGGGACA |
| epikol_00408 | EHMT1_SET_3 | GCTGCAGCTCTACCGGACGC |
| epikol_00409 | EHMT1_SET_4 | GGCTGCAGCTCTACCGGACG |
| epikol_00410 | EHMT1_SET_5 | CCCAGCCCATGTCCCGCGTC |
| epikol_00411 | EHMT2_SET_1 | CGCAGATGAAGGTCCCCTGT |
| epikol_00412 | EHMT2_SET_2 | CTCTACCGAACAGCCAAGAT |
| epikol_00413 | EHMT2_SET_3 | GCTCTACCGAACAGCCAAGA |
| epikol_00414 | EHMT2_SET_4 | CCGAACAGCCAAGATGGGCT |
| epikol_00415 | EHMT2_SET_5 | TCGCAGATGAAGGTCCCCTG |
| epikol_00416 | ELP3_1 | ATCAAGACCTTGCGATACTG |
| epikol_00417 | ELP3_2 | GCAATGATATCCACCAGGCG |
| epikol_00418 | ELP3_3 | CGGCAGCAATGATATCCACC |
| epikol_00419 | ELP3_4 | ATGGGTTTCGCCTTTAACTT |
| epikol_00420 | ELP3_5 | TGATATCCACCAGGCGGGGC |
| epikol_00421 | EP300_BROMO_1 | TTTGGAGGCACTTTACCGTC |
| epikol_00422 | EP300_BROMO_2 | GGGTCCACAGGTTGACGAAA |
| epikol_00423 | EP300_BROMO_3 | AGGGTCCACAGGTTGACGAA |
| epikol_00424 | EP300_BROMO_4 | CCTTCCCTTTCGTCAACCTG |
| epikol_00425 | EP300_BROMO_5 | CCTAAAAGCTGAGGGTCCAC |
| epikol_00426 | EP300_RING_1 | CTGTAATAAGTGGCATCACG |
| epikol_00427 | EP300_RING_2 | CCGTAGCAACACAGTGTCTG |
| epikol_00428 | EP300_RING_3 | CCACAGACACTGTGTTGCTA |

| | | |
|---|---|---|
| epikol_00429 | EP300_RING_4 | GTTCTGGTAACTGTAATAAG |
| epikol_00430 | EP300_RING_5 | CCAAAGAAACGCTCTCCCCT |
| epikol_00431 | EP400_HELIC_1 | CATCGATTCTTACATAGGTG |
| epikol_00432 | EP400_HELIC_2 | ATTGAAATCTGAAGGACGTC |
| epikol_00433 | EP400_HELIC_3 | AGGAGTGGTGCGATAGGATC |
| epikol_00434 | EP400_HELIC_4 | AATTGAAATCTGAAGGACGT |
| epikol_00435 | EP400_HELIC_5 | TTCTTACATAGGTGAGGTAA |
| epikol_00436 | ERCC6_HELIC_1 | TTGGGTACTGGAAACGTTCT |
| epikol_00437 | ERCC6_HELIC_2 | GTTGAAAATATGGCACAAGC |
| epikol_00438 | ERCC6_HELIC_3 | TTGAAAATATGGCACAAGCA |
| epikol_00439 | ERCC6_HELIC_4 | TCATTGTATCTCGTAATCAG |
| epikol_00440 | ERCC6_HELIC_5 | TTGAGTCTTTGTTGAAAATA |
| epikol_00441 | EZH1_SET_1 | CGGATGGGGCACCTTCATAA |
| epikol_00442 | EZH1_SET_2 | CCCCTCTGATGTGGCCGGAT |
| epikol_00443 | EZH1_SET_3 | CCCCCTCTGATGTGGCCGGA |
| epikol_00444 | EZH1_SET_4 | CTTTATGAAGGTGCCCCATC |
| epikol_00445 | EZH1_SET_5 | GCCCCATCCGGCCACATCAG |
| epikol_00446 | EZH2_SET_1 | CTGGCACCATCTGACGTGGC |
| epikol_00447 | EZH2_SET_2 | CACCATCTGACGTGGCAGGC |
| epikol_00448 | EZH2_SET_3 | ACCATCTGACGTGGCAGGCT |
| epikol_00449 | EZH2_SET_4 | CCCCAGCCTGCCACGTCAGA |
| epikol_00450 | EZH2_SET_5 | CATCTGACGTGGCAGGCTGG |
| epikol_00451 | GCN5_HELIC_1 | AACAGAGTATGGAGGTTATC |
| epikol_00452 | GCN5_HELIC_2 | TGGGGAGGTAACAAAGATAT |
| epikol_00453 | GCN5_HELIC_3 | ACAGAGTATGGAGGTTATCA |
| epikol_00454 | GCN5_HELIC_4 | CAAATTCAAACAGAGTATGG |
| epikol_00455 | GCN5_HELIC_5 | TATGGAGGTTATCAGGGCTT |
| epikol_00456 | GCN5_NAT_1 | CCAGACAGACCACCAAAGTC |
| epikol_00457 | GCN5_NAT_2 | GCTGTTCACCCAGATTATCA |
| epikol_00458 | GCN5_NAT_3 | CCAGACTTTGGTGGTCTGTC |
| epikol_00459 | GCN5_NAT_4 | CTGTTCACCCAGATTATCAA |
| epikol_00460 | GCN5_NAT_5 | GACTTTGGTGGTCTGTCTGG |
| epikol_00461 | GTF3C4_1 | CCGAGGTGCGGTGGATAACC |
| epikol_00462 | GTF3C4_2 | CATCTGCGACGTGCACAACC |
| epikol_00463 | GTF3C4_3 | ATCTGCGACGTGCACAACCC |
| epikol_00464 | GTF3C4_4 | TGCGGTGGATAACCAGGTCC |
| epikol_00465 | GTF3C4_5 | CGACGTGCACAACCCGGGCC |
| epikol_00466 | H2AFY2_1 | GTTCAAGTACCGGATCAGCG |
| epikol_00467 | H2AFY2_2 | TTCAAGTACCGGATCAGCGT |
| epikol_00468 | H2AFY2_3 | CAAGCTGTCCCGTTCAGCTA |
| epikol_00469 | H2AFY2_4 | GGCGGCAGTCATTGAGTACC |
| epikol_00470 | H2AFY2_5 | CAGGGGCGCCCACGCTGATC |
| epikol_00471 | H2AFZ_1 | CTTATCAAGGCTACCATAGC |
| epikol_00472 | H2AFZ_2 | TTATCAAGGCTACCATAGCT |
| epikol_00473 | H2AFZ_GECKO_1 | ACCCATTTACCTCTGCGGTG |
| epikol_00474 | H2AFZ_GECKO_2 | CGCGCCGCAGGCTGGCGGTA |
| epikol_00475 | H2AFZ_GECKO_3 | TTCATCGACACCTAAAATCT |
| epikol_00476 | HAT1_1 | AAGAGTTGATGGGTATACTC |

| | | |
|---|---|---|
| epikol_00477 | HAT1_2 | GAGTATACCCATCAACTCTT |
| epikol_00478 | HAT1_3 | GTATACCCATCAACTCTTTG |
| epikol_00479 | HAT1_4 | GTATGAAGTAAGGTTCCGAA |
| epikol_00480 | HAT1_5 | CAACACGGAACATTGTTGAC |
| epikol_00481 | HBO1_1 | ACCAGGTATCAAGCTCATAG |
| epikol_00482 | HBO1_2 | AGGATATGGAGAATGATACC |
| epikol_00483 | HBO1_3 | CGATATATCTCATCACCAGG |
| epikol_00484 | HBO1_4 | CTGCCAAAACCTGTGCCTGT |
| epikol_00485 | HBO1_5 | CTTGAAAATTATGCAGGTAG |
| epikol_00486 | HCFC1_1 | GGGACTCGCCTCCTGGTGTT |
| epikol_00487 | HCFC1_2 | CTCGCCTCCTGGTGTTTGGT |
| epikol_00488 | HCFC1_3 | ACTCGCCTCCTGGTGTTTGG |
| epikol_00489 | HCFC1_4 | TTGGTGGGATGGTGGAGTAT |
| epikol_00490 | HCFC1_5 | CCTCCTGGTGTTTGGTGGGA |
| epikol_00491 | HDAC1_HDAC_1 | AGTAGTAACAGACTTTCCTC |
| epikol_00492 | HDAC1_HDAC_2 | TGAGTCATGCGGATTCGGTG |
| epikol_00493 | HDAC1_HDAC_3 | CTATGGTCTCTACCGAAAAA |
| epikol_00494 | HDAC1_HDAC_4 | TTCGGTGAGGCTTCATTGGG |
| epikol_00495 | HDAC1_HDAC_5 | GGATTCGGTGAGGCTTCATT |
| epikol_00496 | HDAC10_HDAC_1 | AGGACGCTCGATCTCGCACT |
| epikol_00497 | HDAC10_HDAC_2 | TCTGCGGTTGTCAGCCCGCG |
| epikol_00498 | HDAC10_HDAC_3 | GGACGCTCGATCTCGCACTC |
| epikol_00499 | HDAC10_HDAC_4 | GTTGTCAGCCCGCGAGGCCT |
| epikol_00500 | HDAC10_HDAC_5 | CGCGCTGCCGCAGGCGATCC |
| epikol_00501 | HDAC11_HDAC_1 | AGGCCCCTTCGGACCCAGAC |
| epikol_00502 | HDAC11_HDAC_2 | TTCCTCCTGTCTGGGTCCGA |
| epikol_00503 | HDAC11_HDAC_3 | ATCCCTTTGATGCCGGAAAA |
| epikol_00504 | HDAC11_HDAC_4 | CCCTTTGATGCCGGAAAATG |
| epikol_00505 | HDAC11_HDAC_5 | TCCCTTTGATGCCGGAAAAT |
| epikol_00506 | HDAC2_HDAC_1 | TGGGTCATGCGGATTCTATG |
| epikol_00507 | HDAC2_HDAC_2 | GCGGATTCTATGAGGCTTCA |
| epikol_00508 | HDAC2_HDAC_3 | ACAGCAAGTTATGGGTCATG |
| epikol_00509 | HDAC2_HDAC_4 | CATAACTTGCTGTTAAATTA |
| epikol_00510 | HDAC2_HDAC_5 | CATAATTTAACAGCAAGTTA |
| epikol_00511 | HDAC3_HDAC_1 | TATTTCTACGACCCCGACGT |
| epikol_00512 | HDAC3_HDAC_2 | CTATTTCTACGACCCCGACG |
| epikol_00513 | HDAC3_HDAC_3 | CACGTCGGGGTCGTAGAAAT |
| epikol_00514 | HDAC3_HDAC_4 | AGTGGAAGTTGCCCACGTCG |
| epikol_00515 | HDAC3_HDAC_5 | TAGTGGAAGTTGCCCACGTC |
| epikol_00516 | HDAC4_HDAC_1 | GCATCAGCGTGTCATACACG |
| epikol_00517 | HDAC4_HDAC_2 | TGGATCCTCCCGGCGTGCTC |
| epikol_00518 | HDAC4_HDAC_3 | TCCTCCCGGCGTGCTCGGGG |
| epikol_00519 | HDAC4_HDAC_4 | CTGCAGGAGACGGGCCTCCG |
| epikol_00520 | HDAC4_HDAC_5 | TGCTACTCCCGCAGGTGCAC |
| epikol_00521 | HDAC5_HDAC_1 | GTGTGTTCCCGCACATGCAC |
| epikol_00522 | HDAC5_HDAC_2 | TGGATCCGGCCAGCATGCTC |
| epikol_00523 | HDAC5_HDAC_3 | GGATCCGGCCAGCATGCTCA |
| epikol_00524 | HDAC5_HDAC_4 | TAAAGCACCAGTGCATGTGC |

| | | |
|---|---|---|
| epikol_00525 | HDAC5_HDAC_5 | CACGTGCACCCTGAGCATGC |
| epikol_00526 | Ino80_HELIC_3 | GCTTTGTCTGCCCTAAGCGG |
| epikol_00527 | Ino80_HELIC_4 | TGGACAGGGCCCACCGCTTA |
| epikol_00528 | Ino80_HELIC_5 | CTTTGTCTGCCCTAAGCGGT |
| epikol_00529 | Jarid1a_GECKO_5 | ACGACTGACCCAAGTGTCAT |
| epikol_00530 | Jarid1a_JMJC_1 | CCATGCTCCATTAGCACGTT |
| epikol_00531 | Jarid1a_JMJC_2 | CATGCTCCATTAGCACGTTG |
| epikol_00532 | Jarid1a_JMJC_3 | TAGCACGTTGGGGTTCATGA |
| epikol_00533 | Jarid1a_JMJC_4 | GGCTGGGATTCAAATAACTC |
| epikol_00534 | Jarid1a_JMJC_5 | TTTCCTCGTGCCTATCACTC |
| epikol_00535 | Jarid1a_PHD_1 | CCACTACCTGATGTGCCCAA |
| epikol_00536 | Jarid1a_PHD_2 | CCTTTGGGCACATCAGGTAG |
| epikol_00537 | Jarid1a_PHD_3 | TTGGGCACATCAGGTAGTGG |
| epikol_00538 | Jarid1a_PHD_4 | CTGATGTGCCCAAAGGAGAC |
| epikol_00539 | Jarid1b_JMJC_1 | ATACCCTGGGACTCCATACC |
| epikol_00540 | Jarid1b_JMJC_2 | CCCCAGCACACTGATTAGTT |
| epikol_00541 | Jarid1b_JMJC_3 | CCTCAGCAAAATTAAAACCC |
| epikol_00542 | Jarid1b_JMJC_4 | TACCGAACTAATCAGTGTGC |
| epikol_00543 | Jarid1b_JMJC_5 | TTTCCAAGAGCCTACCACAG |
| epikol_00544 | Jarid1b_PHD_1 | GACCGGCTACTGTTGTGTGA |
| epikol_00545 | Jarid1b_PHD_2 | AGCCATCACACAACAGTAGC |
| epikol_00546 | Jarid1b_PHD_3 | CCTCTCCATGATGTTCCCAA |
| epikol_00547 | Jarid1b_PHD_4 | CCCTTGGGAACATCATGGAG |
| epikol_00548 | Jarid1b_PHD_5 | AGTCTCCCTTGGGAACATCA |
| epikol_00549 | Jarid1c_JMJC_1 | ACTCCATTAACTACCTCCAC |
| epikol_00550 | Jarid1c_JMJC_2 | GTGAGCCGAAGACCTGGTAT |
| epikol_00551 | Jarid1c_JMJC_3 | AAGTGAGGGCACCCCATACC |
| epikol_00552 | Jarid1c_JMJC_4 | CCATGGGACATGAGGGTGTT |
| epikol_00553 | Jarid1c_JMJC_5 | AGCCGCTGTGGTAAGCACGG |
| epikol_00554 | Jarid1c_PHD_1 | GGCACCGCCAGACACCCTTG |
| epikol_00555 | Jarid1c_PHD_2 | GATGTGTTCTCGAGGGGATG |
| epikol_00556 | Jarid1c_PHD_3 | TCTGCCGGATGTGTTCTCGA |
| epikol_00557 | Jarid1c_PHD_4 | CATCCCCTCGAGAACACATC |
| epikol_00558 | Jarid1c_PHD_5 | CTGAGATCCCCAAGGGTGTC |
| epikol_00559 | Jarid1d_JMJC_1 | ACTCTATTAACTATCTGCAT |
| epikol_00560 | Jarid1d_JMJC_2 | CTGGTATGGTGTACCCTCCC |
| epikol_00561 | Jarid1d_JMJC_3 | CCCAACACTTTGATGTCCCA |
| epikol_00562 | Jarid1d_JMJC_4 | TCCGCACAAACCAGTGTGCA |
| epikol_00563 | Jarid1d_JMJC_5 | GGCTGGCTATCAAACAGCTC |
| epikol_00564 | Jarid1d_PHD_1 | CATCCCCACGGGAGCATACT |
| epikol_00565 | Jarid1d_PHD_2 | TTGCCAAGTATGCTCCCGTG |
| epikol_00566 | Jarid1d_PHD_3 | ATTTGCCAAGTATGCTCCCG |
| epikol_00567 | Jarid1d_PHD_4 | TTTGCCAAGTATGCTCCCGT |
| epikol_00568 | Jarid1d_PHD_5 | CTGAAATCCCCAGAGGCATC |
| epikol_00569 | Jarid2_JMJC_1 | TACATTGACTACTTACACAC |
| epikol_00570 | Jarid2_JMJC_2 | GTGTGTAAGTAGTCAATGTA |
| epikol_00571 | Jarid2_JMJC_3 | AAGAGGGGATCAAGGTGCAC |
| epikol_00572 | Jarid2_JMJC_4 | CCGGGAAGCAGACGACAAAC |

| | | |
|---|---|---|
| epikol_00573 | Jarid2_JMJC_5 | CTCTTTGCACAGCACCTCCG |
| epikol_00574 | Jarid2_JMJCN_1 | CATCGAGTCGGTCCGCGCTC |
| epikol_00575 | Jarid2_JMJCN_2 | TCCGCTCATCTACATCGAGT |
| epikol_00576 | Jarid2_JMJCN_3 | CGGATCGTGGAACTCCTTGG |
| epikol_00577 | Jarid2_JMJCN_4 | ATCGTGGAACTCCTTGGCGG |
| epikol_00578 | Jarid2_JMJCN_5 | GGGTGATCCCCCCTCCGGAC |
| epikol_00579 | Jhdm1d_JMJC_1 | ACTTCGGTGGAACTTCAGTC |
| epikol_00580 | Jhdm1d_JMJC_2 | ATTTGGCACGTTATGAATCT |
| epikol_00581 | Jhdm1d_JMJC_3 | CTCCAAAGAACACCTCACTC |
| epikol_00582 | Jhdm1d_JMJC_4 | CCATGCTGTGCTCACTTCTC |
| epikol_00583 | Jhdm1d_JMJC_5 | CCTGAGAAGTGAGCACAGCA |
| epikol_00584 | Jmjd1a_JMJC_1 | GCCATCTCGCCTTGTGTACT |
| epikol_00585 | Jmjd1a_JMJC_2 | CGCCTTGTGTACTCGGGCAG |
| epikol_00586 | Jmjd1a_JMJC_3 | CCCGAGTACACAAGGCGAGA |
| epikol_00587 | Jmjd1a_JMJC_4 | GTTCCATATTTCCGATCTTC |
| epikol_00588 | Jmjd1a_JMJC_5 | GCTAATGTCATGGTCTATGT |
| epikol_00589 | Jmjd1b_JMJC_1 | ATTGAGCCTGCCATCTCGTT |
| epikol_00590 | Jmjd1b_JMJC_2 | GGCTACCTAGCTACTTTGTA |
| epikol_00591 | Jmjd1b_JMJC_3 | CTACTTTGTAAGGCCTGATC |
| epikol_00592 | Jmjd1b_JMJC_4 | AATATACCAAACGAGATGGC |
| epikol_00593 | Jmjd1b_JMJC_5 | CCATCTCGTTTGGTATATTC |
| epikol_00594 | Jmjd1b_JMJCSF_1 | ACGAAGCAGAGGATTCATGA |
| epikol_00595 | Jmjd1b_JMJCSF_2 | CCTCATAGAGTCGCTTACGG |
| epikol_00596 | Jmjd1b_JMJCSF_3 | CTCATAGAGTCGCTTACGGA |
| epikol_00597 | Jmjd1b_JMJCSF_4 | ACTCCTCATAGAGTCGCTTA |
| epikol_00598 | Jmjd1b_JMJCSF_5 | CCTCCGTAAGCGACTCTATG |
| epikol_00599 | Jmjd1c_JMJCSF_1 | TGAACAAAAAGCTCCGTCAA |
| epikol_00600 | Jmjd1c_JMJCSF_2 | TCACGTATTGGATCATGTTC |
| epikol_00601 | Jmjd1c_JMJCSF_3 | CATGTTCTGGTAGAACTTCA |
| epikol_00602 | Jmjd1c_JMJCSF_4 | ATCCAATACGTGACCAAAGT |
| epikol_00603 | Jmjd1c_JMJCSF_5 | AAGGAACTGAATAAGAGTAC |
| epikol_00604 | Jmjd2a_JMJC_1 | TACTCTGTTCCACCTGAGCA |
| epikol_00605 | Jmjd2a_JMJC_2 | CAGCATTAACGGGGAAATCA |
| epikol_00606 | Jmjd2a_JMJC_3 | ATATTTCTTCAGCATTAACG |
| epikol_00607 | Jmjd2a_JMJC_4 | AGCCGGCATGGTAACCATAA |
| epikol_00608 | Jmjd2a_JMJC_5 | ACTCCGCACAGTTAAAACCA |
| epikol_00609 | Jmjd2a_JMJCN_1 | TCCCAAGGAGCTCATCGGGC |
| epikol_00610 | Jmjd2a_JMJCN_2 | TCATCGGGCAGGGCTAGCCA |
| epikol_00611 | Jmjd2a_JMJCN_3 | TTGAATCCCAAGGAGCTCAT |
| epikol_00612 | Jmjd2a_JMJCN_4 | GCCCTGCCCGATGAGCTCCT |
| epikol_00613 | Jmjd2a_JMJCN_5 | TTTCGGAACTCTTCCATAGT |
| epikol_00614 | Jmjd2b_JMJC_1 | ACCAGAGCACGGCAAGCGCC |
| epikol_00615 | Jmjd2b_JMJC_2 | TACGCCATCCCACCAGAGCA |
| epikol_00616 | Jmjd2b_JMJC_3 | AGGGGATCCCGTACTTCTTC |
| epikol_00617 | Jmjd2b_JMJC_4 | CGCAGCCCTGCGAGCTCCCG |
| epikol_00618 | Jmjd2b_JMJC_5 | ACCACGCCGGCTTCAATCAC |
| epikol_00619 | Jmjd2b_JMJCN_1 | CGTGGCCTACATAGAGTCGC |
| epikol_00620 | Jmjd2b_JMJCN_2 | GTGGCCTACATAGAGTCGCA |

| | | |
|---|---|---|
| epikol_00621 | Jmjd2b_JMJCN_3 | GGCTCCCTGCGACTCTATGT |
| epikol_00622 | Jmjd2b_JMJCN_4 | CTTGGCCAGGCCCGCCCGGT |
| epikol_00623 | Jmjd2b_JMJCN_5 | GGGAGCCCACCGGGCGGGCC |
| epikol_00624 | Jmjd2c_JMJC_1 | GGAGATAATTAATGCTATAG |
| epikol_00625 | Jmjd2c_JMJC_2 | AGCATTAATTATCTCCACTT |
| epikol_00626 | Jmjd2c_JMJC_3 | TATGCTATACCTCCGGAGCA |
| epikol_00627 | Jmjd2c_JMJC_4 | AACCAGCATGGTAGCCATAT |
| epikol_00628 | Jmjd2c_JMJC_5 | TCATGAATTCTCCAGCCTCC |
| epikol_00629 | Jmjd2c_JMJCN_1 | CCTTTGCAAGACCCGCACGA |
| epikol_00630 | Jmjd2c_JMJCN_2 | CCATCGTGCGGGTCTTGCAA |
| epikol_00631 | Jmjd2c_JMJCN_3 | TCTAAAGGAGCCCATCGTGC |
| epikol_00632 | Jmjd2c_JMJCN_4 | CTTTAGACTCCATGTATGCA |
| epikol_00633 | Jmjd2c_JMJCN_5 | CAAGGTATTTGTTGAACTCC |
| epikol_00634 | JMJD2D_GECKO_1 | AACCTGAACGCTATGACCTG |
| epikol_00635 | JMJD2D_GECKO_2 | GCTCCAAATCTTCGAAATTC |
| epikol_00636 | JMJD2D_GECKO_3 | CAGATTATCCACCCGTCAAA |
| epikol_00637 | Jmjd2d_JMJC_1 | TGGCCAGGCGTTCCAGGCGC |
| epikol_00638 | Jmjd2d_JMJC_2 | CCATGGTTTCAACTGCGCAG |
| epikol_00639 | Jmjd2d_JMJC_3 | CCACAACCCCGGGAACTGCC |
| epikol_00640 | Jmjd2d_JMJC_4 | CCTCTGCGCAGTTGAAACCA |
| epikol_00641 | Jmjd2d_JMJCN_1 | AGCCAGCTCTGTGTGCACCT |
| epikol_00642 | Jmjd2d_JMJCN_2 | TCATTAAACTCTTCTTTGGT |
| epikol_00643 | Jmjd2d_JMJCN_3 | AGGTGCACACAGAGCTGGCT |
| epikol_00644 | JmjD3_JMJC_1 | GCGAACCACTCGCAGTCGCC |
| epikol_00645 | JmjD3_JMJC_2 | TCGCGGTGCACGAGCACTAC |
| epikol_00646 | JmjD3_JMJC_3 | CGCGGTGCACGAGCACTACT |
| epikol_00647 | JmjD3_JMJC_4 | TGCTCCGTCAACATCAACAT |
| epikol_00648 | JmjD3_JMJC_5 | ACCTCGTGTGGATTAATGCG |
| epikol_00649 | Jmjd4_JMJC_1 | TGGACCCCACAGTTTGCAAC |
| epikol_00650 | Jmjd4_JMJC_2 | TCAGAGACTACATCACCTAC |
| epikol_00651 | Jmjd4_JMJC_3 | AGGCGGGCTACTCCTCTCCC |
| epikol_00652 | Jmjd4_JMJC_4 | GCTGTCTCTACCTCAAAGAC |
| epikol_00653 | Jmjd4_JMJC_5 | AGTTGCAAACTGTGGGGTCC |
| epikol_00654 | Jmjd5_CUPIN_1 | AGTGGTCAGCCACGCCTTTC |
| epikol_00655 | Jmjd5_CUPIN_2 | GCCACGCCTTTCAGGATCAC |
| epikol_00656 | Jmjd5_CUPIN_3 | ACTGGCCGTGCATGCAGAAG |
| epikol_00657 | Jmjd5_CUPIN_4 | GAACTCGTTGACCGTCATGA |
| epikol_00658 | Jmjd5_CUPIN_5 | CGTTGACCGTCATGAGGGTC |
| epikol_00659 | Jmjd6_JMJC_1 | ATCAAAGTGACCCGAGACGA |
| epikol_00660 | Jmjd6_JMJC_2 | GGGCCACCACGCTCCGGAAC |
| epikol_00661 | Jmjd6_JMJC_3 | TGGGATTCACATCGACCCTC |
| epikol_00662 | Jmjd6_JMJC_4 | TAGTTCAGGGCCACAAGCGC |
| epikol_00663 | Jmjd6_JMJC_5 | AAAGTGACCCGAGACGAAGG |
| epikol_00664 | Jmjd7_CUPIN_1 | GCGGATAATGCACGGCCTGT |
| epikol_00665 | Jmjd7_CUPIN_2 | CGGATAATGCACGGCCTGTT |
| epikol_00666 | Jmjd7_CUPIN_3 | AGAGCGTTGCGGATAATGCA |
| epikol_00667 | Jmjd7_CUPIN_4 | CGCTCCACTTCTACCGGGAC |
| epikol_00668 | Jmjd7_CUPIN_5 | GGATAATGCACGGCCTGTTG |

| | | |
|---|---|---|
| epikol_00669 | Jmjd8_JMJC_1 | CAGGATGACGGGCCTGACGA |
| epikol_00670 | Jmjd8_JMJC_2 | CGTCAGGCCCGTCATCCTGC |
| epikol_00671 | Jmjd8_JMJC_3 | GCAACCTGTCGCGGGAGCAC |
| epikol_00672 | Jmjd8_JMJC_4 | CGAAGCCAGCAACCTGTCGC |
| epikol_00673 | Jmjd8_JMJC_5 | GGGCCCTGTGCTCCCGCGAC |
| epikol_00674 | Kat2a_BROMO_1 | GATGACCTCGTAGTAGTCAG |
| epikol_00675 | Kat2a_BROMO_2 | GCGGCAGTACTCGCTGTCCG |
| epikol_00676 | Kat2a_BROMO_3 | CTGTCGCGAGTACAACCCCC |
| epikol_00677 | Kat2a_BROMO_4 | GAAGCCGCTACTACGTGACC |
| epikol_00678 | Kat2a_BROMO_5 | CGGCAGTACTCGCTGTCCGG |
| epikol_00679 | Kat2a_NAT_1 | GACAATCTCCGTGAAGCCCT |
| epikol_00680 | Kat2a_NAT_2 | GGCCTTGATCAAGGATGGGC |
| epikol_00681 | Kat2a_NAT_3 | GCTTGATGTGATACTCCTTC |
| epikol_00682 | Kat2a_NAT_4 | TGATGTGATACTCCTTCAGG |
| epikol_00683 | Kat2a_NAT_5 | CCTTCAGGTGGTTCATCAGG |
| epikol_00684 | Kat2b_BROMO_1 | TGAAGGGCCAAGCGCTTTGA |
| epikol_00685 | Kat2b_BROMO_2 | CAGGATATTATGAAGTTATA |
| epikol_00686 | Kat2b_BROMO_3 | ATTCTTGAGGCGTTCACTCA |
| epikol_00687 | Kat2b_BROMO_4 | ACACGTAGTACCTATTCTTG |
| epikol_00688 | Kat2b_BROMO_5 | GGGGTTGTACTCTTTGCAAT |
| epikol_00689 | Kat2b_NAT_1 | ATTAAAGATGGCCGTGTTAT |
| epikol_00690 | Kat2b_NAT_2 | AAAGATGGCCGTGTTATTGG |
| epikol_00691 | Kat2b_NAT_3 | TCTCTGTGAATCCTTGAGAT |
| epikol_00692 | Kat2b_NAT_4 | TTCCGTATGTTCCCATCTCA |
| epikol_00693 | Kat2b_NAT_5 | AACAGATACCACCAATAACA |
| epikol_00694 | Kat5_CHROM_1 | ATGGGTGACGCATGAGCGGC |
| epikol_00695 | Kat5_CHROM_2 | ATGAATGGGTGACGCATGAG |
| epikol_00696 | Kat5_CHROM_3 | GTCCTTCACGCTCAGGATCT |
| epikol_00697 | Kat5_CHROM_4 | GGCCGAGATCCTGAGCGTGA |
| epikol_00698 | Kat5_CHROM_5 | GCGTGAAGGACATCAGTGGC |
| epikol_00699 | Kat5_NAT_1 | ATCAACGGAAGACTACAATG |
| epikol_00700 | Kat5_NAT_2 | GCTTGCCGTAGCCCCGGCGC |
| epikol_00701 | Kat5_NAT_3 | CTGCCTCCCTACCAGCGCCG |
| epikol_00702 | Kat5_NAT_4 | TCAGCAGCTTGCCGTAGCCC |
| epikol_00703 | Kat5_NAT_5 | GCCGTAGCCCCGGCGCTGGT |
| epikol_00704 | L3mbtl1_MBT_1 | AGTACATGGACGGGTGTTGA |
| epikol_00705 | L3mbtl1_MBT_2 | AAGTACATGGACGGGTGTTG |
| epikol_00706 | L3mbtl1_MBT_3 | GCAGTCACTCACAACAAGAA |
| epikol_00707 | L3mbtl1_MBT_4 | CATGTACTTCATCCTCACCG |
| epikol_00708 | L3mbtl1_MBT_5 | TCTTGTTGTGAGTGACTGCC |
| epikol_00709 | LSD2_AMOX_1 | CAACGGTGGATTAAACTGAA |
| epikol_00710 | LSD2_AMOX_2 | TCAATTATCACCGAGTACCC |
| epikol_00711 | LSD2_AMOX_3 | TTCAATTATCACCGAGTACC |
| epikol_00712 | LSD2_AMOX_4 | TCTGCTAACTCCCGGGTACT |
| epikol_00713 | LSD2_AMOX_5 | GAGTGTGGTCACCAGCAAAC |
| epikol_00714 | Mbd1_MEC_1 | CGCGAAGTCTTTCGCAAGTC |
| epikol_00715 | Mbd1_MEC_2 | ATAGGTGTCTGAGCGTCCAC |
| epikol_00716 | Mbd1_MEC_3 | CTTCGCGGCGCTTCCAGCCA |

| | | |
|---|---|---|
| epikol_00717 | Mbd1_MEC_4 | GTTGAGCTGACTCGATACCT |
| epikol_00718 | Mbd1_MEC_5 | GCCTTGTTTGAAGTCGAAGA |
| epikol_00719 | Mbd2_MEC_1 | CCTTCTTCCATCCGGGGGGG |
| epikol_00720 | Mbd2_MEC_2 | GAGGAAGTGATCCGAAAATC |
| epikol_00721 | Mbd2_MEC_3 | CCTCCCCCCCGGATGGAAGA |
| epikol_00722 | Mbd2_MEC_4 | AGGAAGTGATCCGAAAATCT |
| epikol_00723 | Mbd2_MEC_5 | CCCCCCCGGATGGAAGAAGG |
| epikol_00724 | Mbd3_MEC_1 | GCCGACAGCCCCGACCTTCT |
| epikol_00725 | Mbd3_MEC_2 | CGGGGCTGTCGGCCGGCCAC |
| epikol_00726 | Mbd3_MEC_3 | GGCCGACAGCCCCGACCTTC |
| epikol_00727 | Mbd3_MEC_4 | GGAGTGCCCGGCGCTCCCGC |
| epikol_00728 | Mbd3_MEC_5 | AGAAGGTCGGGGCTGTCGGC |
| epikol_00729 | Mbd4_MEC_1 | GATCTGAACTTCAGTCCTTG |
| epikol_00730 | Mbd4_MEC_2 | GTTGTGAAGCAAAGGTTATT |
| epikol_00731 | Mbd4_MEC_3 | GCTAATTATCTTCACAAAAA |
| epikol_00732 | Mbd4_MEC_4 | TTTACTGTACTTTCTAAAAG |
| epikol_00733 | Mbd4_MEC_5 | ACAGTAAAATCAAAATCTTC |
| epikol_00734 | Mecp2_MEC_1 | GGACACGGAAGCTTAAGCAA |
| epikol_00735 | Mecp2_MEC_2 | TCCGTGTCCAGCCTTCAGGC |
| epikol_00736 | Mecp2_MEC_3 | GACTTCACGGTAACTGGGAG |
| epikol_00737 | Mecp2_MEC_4 | ATTGCGTACTTCGAAAAGGT |
| epikol_00738 | Mecp2_MEC_5 | AAAAGCCTTTCGCTCTAAAG |
| epikol_00739 | MgeA5_NAG_1 | GCGAGAGATGTATTCAGTGG |
| epikol_00740 | MgeA5_NAG_2 | GGTGTGGATAGCAACGTAGT |
| epikol_00741 | MgeA5_NAG_3 | ACCTCTTCGATGGACTCTAC |
| epikol_00742 | MgeA5_NAG_4 | CTATGCGATCTCACCTGGAT |
| epikol_00743 | MgeA5_NAG_5 | TATGTGTGCAGCAGACAAAG |
| epikol_00744 | Mina_CUPIN_1 | AATGTCTGCCGGTGTGTCAA |
| epikol_00745 | Mina_CUPIN_2 | CTCTTCCATAGTACATCCCC |
| epikol_00746 | Mina_CUPIN_3 | CTGCACTGGCCACATACTAT |
| epikol_00747 | Mina_CUPIN_4 | AATGTGTACATAACTCCCGC |
| epikol_00748 | Mina_CUPIN_5 | CATCATCATAATGGGGCGGC |
| epikol_00749 | Mll1_BROMO_1 | CCGGTAGCGTAGCAAATGGC |
| epikol_00750 | Mll1_BROMO_2 | TCTGTCTCGGGATTTAAGTC |
| epikol_00751 | Mll1_BROMO_3 | GTCTCGGGATTTAAGTCTGG |
| epikol_00752 | Mll1_BROMO_4 | AGGTCCTTCGGGGGAGCTGC |
| epikol_00753 | Mll1_BROMO_5 | CAGCAGCCTTTAGATCTAGA |
| epikol_00754 | Mll1_PHD_1 | GGAAGGGCTCACAACAGACT |
| epikol_00755 | Mll1_PHD_2 | TGGAGGACCAGCTGGAAAAT |
| epikol_00756 | Mll1_PHD_3 | AGAGGAGAACGAGCGCCCTC |
| epikol_00757 | Mll1_PHD_4 | TCTCCTCTAAACAAAACTTG |
| epikol_00758 | Mll1_PHD_5 | CTCTAAACAAAACTTGTGGA |
| epikol_00759 | MLL2_PHD_1 | GCACTGAGAGCACTGCGAAC |
| epikol_00760 | MLL2_PHD_2 | GAGAGCACTGCGAACAGGCA |
| epikol_00761 | MLL2_PHD_3 | TGGTATGTGGCAGCTTTGGC |
| epikol_00762 | MLL2_PHD_4 | ACTGCGAACAGGCAAGGAGG |
| epikol_00763 | MLL2_PHD_5 | ACCAAGGTGATGCTGCTCAA |
| epikol_00764 | MLL2_SET_1 | ATTGATGCTACGTTGACCGG |

| | | |
|---|---|---|
| epikol_00765 | MLL2_SET_2 | CACCATCATTCGGAACGAGG |
| epikol_00766 | MLL2_SET_3 | CTGGCTCGCTCCCGTATCCA |
| epikol_00767 | MLL2_SET_4 | CCTTGGCTGCATAGAGCCCC |
| epikol_00768 | MLL2_SET_5 | GTGATTGATGCTACGTTGAC |
| epikol_00769 | MLL3_GECKO_1 | GACACAGATCGCTGAAGAGT |
| epikol_00770 | MLL3_GECKO_2 | AAACCAACCTTCTAACAAGA |
| epikol_00771 | MLL3_GECKO_3 | TCTTCAGCGATCTGTGTCTG |
| epikol_00772 | MLL3_GECKO_4 | CTCCATGGCAAGATAAATCC |
| epikol_00773 | MLL3_GECKO_5 | CTTGATCATCTGAAGCTGTC |
| epikol_00774 | MLL3_SET_1 | CAATGTCTCGAGCAGCATAC |
| epikol_00775 | MLL3_SET_2 | GTGATTGACGCGACGCTCAC |
| epikol_00776 | MLL3_SET_3 | TGAGCGTCGCGTCAATCACA |
| epikol_00777 | MLL3_SET_4 | TTGACGCGACGCTCACAGGA |
| epikol_00778 | MLL3_SET_5 | ATTGACGCGACGCTCACAGG |
| epikol_00779 | MLL4_PHD_1 | TGCCCCAGCATCACGACACC |
| epikol_00780 | MLL4_PHD_2 | AGGTGTCGTGATGCTGGGGC |
| epikol_00781 | MLL4_PHD_3 | AGCACCAGGTGTCGTGATGC |
| epikol_00782 | MLL4_PHD_4 | GCACCAGGTGTCGTGATGCT |
| epikol_00783 | MLL4_PHD_5 | CACCAGGTGTCGTGATGCTG |
| epikol_00784 | MLL4_SET_1 | AGCGGGAGAAGTTCTACGAT |
| epikol_00785 | MLL4_SET_2 | CGATGTTGCGCTTACAGAAC |
| epikol_00786 | MLL4_SET_3 | AGATCAGCCATCCACGGGCG |
| epikol_00787 | MLL4_SET_4 | CAACATCGACGCGGGGGAGA |
| epikol_00788 | MLL4_SET_5 | AAGCGGGAGAAGTTCTACGA |
| epikol_00789 | MLL5_PHD_1 | CGTTCACATAGATATGTATC |
| epikol_00790 | MLL5_PHD_2 | TTGACTGCATGGGGATTGAT |
| epikol_00791 | MLL5_PHD_3 | GGCAACATATTGACTGCATG |
| epikol_00792 | MLL5_PHD_4 | TGGCAACATATTGACTGCAT |
| epikol_00793 | MLL5_PHD_5 | TTGGCAACATATTGACTGCA |
| epikol_00794 | MLL5_SET_1 | TATTCAATGATAAGTGCATC |
| epikol_00795 | MLL5_SET_2 | TCAATGATAAGTGCATCAGG |
| epikol_00796 | MLL5_SET_3 | GGAATGAGGCTCGATTCATC |
| epikol_00797 | MLL5_SET_4 | ATGAGGCTCGATTCATCAGG |
| epikol_00798 | MLL5_SET_5 | GGAATGAGGCTCGATTCATC |
| epikol_00799 | MMP8_GECKO_1 | ATTCCATTGGGTCCATCAAA |
| epikol_00800 | MMP8_GECKO_2 | TGCAACACTCCAGAGTTCAA |
| epikol_00801 | MMP8_GECKO_3 | CTATACCCCACAGCTGTCAG |
| epikol_00802 | MMP8_GECKO_4 | AACGCACTAACTTGACCTAC |
| epikol_00803 | MMP8_GECKO_5 | GCAACCAGTATCAGTCTACA |
| epikol_00804 | MORF4L1_GECKO_1 | GCCTCTTCTTTATGAAGCAA |
| epikol_00805 | MORF4L1_GECKO_2 | ACATATTACCAAATAATCTC |
| epikol_00806 | MORF4L1_GECKO_3 | ACCTTTGCTTCATAAAGAAG |
| epikol_00807 | MORF4L1_GECKO_4 | CCACATCTCCTGAGATTATT |
| epikol_00808 | MORF4L1_GECKO_5 | CATGCATTTACCTAGAGGTC |
| epikol_00809 | MORF4L1_GECKO_6 | TGTACTGAGGGAGCTCCTTC |
| epikol_00810 | MORF4L1_TUDOR_1 | CTCCTTCAGGAACAGGAAAA |
| epikol_00811 | MORF4L1_TUDOR_2 | GAGGGAGCTCCTTCAGGAAC |
| epikol_00812 | MORF4L1_TUDOR_3 | AGGAGGGGGTGGTGTACTGA |

| | | |
|---|---|---|
| epikol_00813 | MORF4L1_TUDOR_4 | CAGGAGGGGGTGGTGTACTG |
| epikol_00814 | MSL3_GECKO_1 | GTTCAATTAGCTGGGATAGA |
| epikol_00815 | MSL3_GECKO_2 | TCGTAGATTACAGCGTAAAT |
| epikol_00816 | MSL3_GECKO_3 | GTGCCTGACAATTACCCCCC |
| epikol_00817 | MSL3_GECKO_4 | TGAGCAAATGAGCGCGAGCG |
| epikol_00818 | MSL3_GECKO_5 | ATCGTACAGCACTCGCGCCT |
| epikol_00819 | MTA1_BAH_1 | ATACCTGATCCGGAGAATCG |
| epikol_00820 | MTA1_BAH_2 | CGATTCTCCGGATCAGGTAT |
| epikol_00821 | MTA1_BAH_3 | TCGATTCTCCGGATCAGGTA |
| epikol_00822 | MTA1_BAH_4 | CCAAAGTGGTGTGCTTCTAC |
| epikol_00823 | MTA1_BAH_5 | CTATAAGGCCGGACCGGGGG |
| epikol_00824 | MTA1_SANT_1 | GAGGAAGCCCTGGAAAAATA |
| epikol_00825 | MTA1_SANT_2 | CAGGGCTTCCTCGAAAAGGT |
| epikol_00826 | MTA1_SANT_3 | TCATTGAGTACTACTACATG |
| epikol_00827 | MTA1_SANT_4 | GTAGTAGTACTCAATGATGC |
| epikol_00828 | MTA1_SANT_5 | ATGCTGGTCAGCGACTTCCA |
| epikol_00829 | MTA2_BAH_1 | TTACCTGGTTAGACGGATTG |
| epikol_00830 | MTA2_BAH_2 | TCTAACCAGGTAAGGATTGC |
| epikol_00831 | MTA2_BAH_3 | GCAATCCTTACCTGGTTAGA |
| epikol_00832 | MTA2_BAH_4 | GTTCCCGGTGCTTCAGTTGA |
| epikol_00833 | MTA2_BAH_5 | AGATATCTTGAGCCAGTACC |
| epikol_00834 | MTA2_SANT_1 | GGACTTCAATGATATTCGCC |
| epikol_00835 | MTA2_SANT_2 | TAGGGCCTCCTCAAATAGCA |
| epikol_00836 | MTA2_SANT_3 | CTCAGAGGCCATGCTATTTG |
| epikol_00837 | MTA2_SANT_4 | GGCCCTAGAGAAGTATGGGA |
| epikol_00838 | MTA2_SANT_5 | AGAGGCCATGCTATTTGAGG |
| epikol_00839 | MTA3_BAH_1 | TCTTATTAGGTATGGGTTGC |
| epikol_00840 | MTA3_BAH_2 | GCAACCCATACCTAATAAGA |
| epikol_00841 | MTA3_BAH_3 | TCTATCCTTCTTATTAGGTA |
| epikol_00842 | MTA3_BAH_4 | AGTATTGTCATATCTTGATA |
| epikol_00843 | MTA3_BAH_5 | TGATTCTGTCTCATTCAGAA |
| epikol_00844 | MTA3_GECKO_1 | GCAACCCATACCTAATAAGA |
| epikol_00845 | MTA3_GECKO_2 | AAAACACTATTAGCTGACAA |
| epikol_00846 | MTA3_GECKO_3 | GTTCTTCTATCCTTCTTATT |
| epikol_00847 | MTA3_GECKO_4 | TCAATCTGTCGATCCGTAAG |
| epikol_00848 | MTA3_GECKO_5 | ACCGTGCTACAACTAAAAAC |
| epikol_00849 | MYST1_CHROM_1 | GGAATTCTATGTACACTACG |
| epikol_00850 | MYST1_CHROM_2 | CCTGGTCGTTCACTCGAGAC |
| epikol_00851 | MYST1_CHROM_3 | GAATTCTATGTACACTACGT |
| epikol_00852 | MYST1_CHROM_4 | GTGGGTAGACAAGAACCGGC |
| epikol_00853 | MYST1_CHROM_5 | TACAGCATCCTTCACTGTCT |
| epikol_00854 | MYST1_HAT_1 | CAATTTCGTAGTTCCCGATG |
| epikol_00855 | MYST1_HAT_2 | CACCATTCCCCGAAGACTAT |
| epikol_00856 | MYST1_HAT_3 | TCACCATTCCCCGAAGACTA |
| epikol_00857 | MYST1_HAT_4 | GGGGAATGGTGAGAAATACC |
| epikol_00858 | MYST1_HAT_5 | TTCCCATAGTCTTCGGGGAA |
| epikol_00859 | MYST2_GECKO_1 | GGGTGACTCGAGCAGATCGT |
| epikol_00860 | MYST2_GECKO_2 | GACAACTCACCATGTGCCGG |

| | | |
|---|---|---|
| epikol_00861 | MYST2_GECKO_3 | TTCAATCTCTGTGTTTGAAG |
| epikol_00862 | MYST2_GECKO_4 | CTCGAGATCTGTAGAAAAAT |
| epikol_00863 | MYST2_GECKO_5 | TCTTGCAGGCCAAATGTGTG |
| epikol_00864 | MYST3_HAT_1 | TGGCTCCACATCGTAATAGA |
| epikol_00865 | MYST3_HAT_2 | ATGGCTCCACATCGTAATAG |
| epikol_00866 | MYST3_HAT_3 | TCTTCCTCAATACCAGCGTA |
| epikol_00867 | MYST3_HAT_4 | CTAACACAGAATGATGTCAA |
| epikol_00868 | MYST3_HAT_5 | TAGCCCTTACGCTGGTATTG |
| epikol_00869 | MYST3_PHD_1 | ACTGTTGCCACAGTCGGCAC |
| epikol_00870 | MYST3_PHD_2 | CAGAAACTACAGATGGGGAT |
| epikol_00871 | MYST3_PHD_3 | TTCACTCGAACCGTTAGTTC |
| epikol_00872 | MYST3_PHD_4 | TCACTCGAACCGTTAGTTCA |
| epikol_00873 | MYST3_PHD_5 | CACTCGAACCGTTAGTTCAG |
| epikol_00874 | MYST4_HAT_1 | CAGCAAATGAAATTTACCGA |
| epikol_00875 | MYST4_HAT_2 | CTTCGGTAAATTTCATTTGC |
| epikol_00876 | MYST4_HAT_3 | TGATGAAAAGGGCTGTCATC |
| epikol_00877 | MYST4_HAT_4 | ACCGTCCAAATCCTTGCCTT |
| epikol_00878 | MYST4_HAT_5 | ACCAAAGGCAAGGATTTGGA |
| epikol_00879 | MYST4_PHD_1 | CCCAAACAGAAGCTACATAT |
| epikol_00880 | MYST4_PHD_2 | ACAGAAGCTACATATTGGAA |
| epikol_00881 | MYST4_PHD_3 | GCACAAGAGAGGAGTTCTTC |
| epikol_00882 | MYST4_PHD_4 | TGCCACAATCTGCACAAGAG |
| epikol_00883 | MYST4_PHD_5 | TCCAATATGTAGCTTCTGTT |
| epikol_00884 | NAP1L1_GECKO_1 | CATTCACATTCTTCTTCCGT |
| epikol_00885 | NAP1L1_GECKO_2 | TACCTTTCAATGTATCCTGT |
| epikol_00886 | NAP1L1_GECKO_3 | TAATGCAATTTATGAACCTA |
| epikol_00887 | NAP1L1_GECKO_4 | CTATCAGCCTCTATTTGATA |
| epikol_00888 | NAP1L1_GECKO_5 | CACCAACAGGATACATTGAA |
| epikol_00889 | NAP1L2_NAP_1 | CGCAGTTCCCCTATAGGGAT |
| epikol_00890 | NAP1L2_NAP_2 | CTCAATCGCAGTTCCCCTAT |
| epikol_00891 | NAP1L2_NAP_3 | TATGATCCCCATCCCTATAG |
| epikol_00892 | NAP1L2_NAP_4 | ACTGCGATTGAGTATTCCAC |
| epikol_00893 | NAP1L2_NAP_5 | TTATGATCCCCATCCCTATA |
| epikol_00894 | NAP1L3_NAP_1 | CAGTAGCAGTTGTGCGACTC |
| epikol_00895 | NAP1L3_NAP_2 | CAGAGTCGCACAACTGCTAC |
| epikol_00896 | NAP1L3_NAP_3 | TGAAACCGCCTATCATACAG |
| epikol_00897 | NAP1L3_NAP_4 | AGATCCTAAAGAAGTCCCCC |
| epikol_00898 | NAP1L3_NAP_5 | GATTCCTATGATTGGGAAGC |
| epikol_00899 | NAT10_HELIC_1 | ACAGAGTATGGAGGTTATCA |
| epikol_00900 | NAT10_HELIC_2 | GGGCGTATCGGATTGACTCC |
| epikol_00901 | NAT10_HELIC_3 | ATGCCATGAAAACAAGGTAG |
| epikol_00902 | NAT10_HELIC_4 | CGTGGTCTTATTCTCAGCAG |
| epikol_00903 | NAT10_HELIC_5 | TTAGCTTGAGGGACAGTGAC |
| epikol_00904 | NAT10_NAT_1 | CCAGACAGACCACCAAAGTC |
| epikol_00905 | NAT10_NAT_2 | GCTGTTCACCCAGATTATCA |
| epikol_00906 | NAT10_NAT_3 | CCAGACTTTGGTGGTCTGTC |
| epikol_00907 | NAT10_NAT_4 | CTGTTCACCCAGATTATCAA |
| epikol_00908 | NAT10_NAT_5 | GACTTTGGTGGTCTGTCTGG |

| | | |
|---|---|---|
| epikol_00909 | NCOA1_PAS_1 | CGTCTACAGCATACTGCACG |
| epikol_00910 | NCOA1_PAS_2 | CAGCTACTTAGGTTACAATC |
| epikol_00911 | NCOA1_PAS_3 | GAGAATGTAACCAGCTACTT |
| epikol_00912 | NCOA1_PAS_4 | GCATCCTGCAGTTAAAGGTA |
| epikol_00913 | NCOA1_PAS_5 | TTTCGTCGTGTTGCCTCTTG |
| epikol_00914 | NCOA1_SRC_1 | TCTACACCGGCTCTTACAGG |
| epikol_00915 | NCOA1_SRC_2 | CAACTGCCGAACAGCAGTTA |
| epikol_00916 | NCOA1_SRC_3 | GCCTCTGCTAACTCTTCAGG |
| epikol_00917 | NCOA1_SRC_4 | ACAAAGTGGTGATATCTGAG |
| epikol_00918 | NCOA1_SRC_5 | TCTGCCTCTGCTAACTCTTC |
| epikol_00919 | NCOA2_PAS_1 | AAGGTATGGCTGTTCCGCCT |
| epikol_00920 | NCOA2_PAS_2 | CTTGGTCTGGCGAACCTCCG |
| epikol_00921 | NCOA2_PAS_3 | GTATGGCTGTTCCGCCTCGG |
| epikol_00922 | NCOA2_PAS_4 | TACCTTCAATTGTCGGATGC |
| epikol_00923 | NCOA2_PAS_5 | GCATCCGACAATTGAAGGTA |
| epikol_00924 | NCOA2_SRC_1 | GTTTGTATCCGACAAAGAGC |
| epikol_00925 | NCOA2_SRC_2 | TACACATGGAACCTCGCTCA |
| epikol_00926 | NCOA2_SRC_3 | GGGCTCCATCTGATCAGATT |
| epikol_00927 | NCOA2_SRC_4 | GTTAACTTGGCCAAGTCCAC |
| epikol_00928 | NCOA2_SRC_5 | GACCACCAAATCTGATCAGA |
| epikol_00929 | NCOA3_PAS_1 | CGTCTCGATTCACCACAAAT |
| epikol_00930 | NCOA3_PAS_2 | CTATTTGTGGTGAATCGAGA |
| epikol_00931 | NCOA3_PAS_3 | GCAATATAAGCAAGAGGACC |
| epikol_00932 | NCOA3_PAS_4 | GGTCCTCTTGCTTATATTGC |
| epikol_00933 | NCOA3_PAS_5 | TGTAAACACTTGTGTTAACC |
| epikol_00934 | NCOA3_SRC_1 | ACCTGTTCTTCTGATGACCG |
| epikol_00935 | NCOA3_SRC_2 | CCTCTACATCCAATATGCAT |
| epikol_00936 | NCOA3_SRC_3 | TAACAGTGACCCATGCATAT |
| epikol_00937 | NCOA3_SRC_4 | ACCCCGGTCATCAGAAGAAC |
| epikol_00938 | NCOA3_SRC_5 | CCCATGCATATTGGATGTAG |
| epikol_00939 | NO66_CUPIN_1 | TCGTCCGTTGATGTAGCGAG |
| epikol_00940 | NO66_CUPIN_2 | GACGCCGCTCGCTACATCAA |
| epikol_00941 | NO66_CUPIN_3 | CCCCCACTACGACGACATCG |
| epikol_00942 | NO66_CUPIN_4 | GCCTCGATGTCGTCGTAGTG |
| epikol_00943 | NO66_CUPIN_5 | TCCATGAAAGCGGTTCTTCG |
| epikol_00944 | NSD1_PHD_1 | CAGTCAATCCAAGGCACTCC |
| epikol_00945 | NSD1_PHD_2 | TTGGCATCTCAGTCAATCCA |
| epikol_00946 | NSD1_PHD_3 | GCTTTCCACCTGGAGTGCCT |
| epikol_00947 | NSD1_PHD_4 | GGGTGAGCTGCTGTTATGTG |
| epikol_00948 | NSD1_PHD_5 | TATGTGAGGCTCAGTGCTGT |
| epikol_00949 | NSD1_SET_1 | CCGCACATTACAGCGGGGTT |
| epikol_00950 | NSD1_SET_2 | CCCAACCCCGCTGTAATGTG |
| epikol_00951 | NSD1_SET_3 | ACACAGAAGTGGTCTGTGAA |
| epikol_00952 | NSD1_SET_4 | GTGAATGGAGATACCCGTGT |
| epikol_00953 | NSD1_SET_5 | TAGTGCAAAAAGGCCTACAC |
| epikol_00954 | NSD2_PHD_1 | AGGTGGAAAGCTCCGCAGCA |
| epikol_00955 | NSD2_PHD_2 | CTGCAGGTGAACCTCCCTTC |
| epikol_00956 | NSD2_PHD_3 | GGCTTTCCCGGAGGCCAGAA |

| | | |
|---|---|---|
| epikol_00957 | NSD2_PHD_4 | CCTGCCTTGGGCTTTCCCGG |
| epikol_00958 | NSD2_PHD_5 | GGGCTTTCCCGGAGGCCAGA |
| epikol_00959 | NSD2_SET_1 | TGATGTCCCTCTTGGCGACC |
| epikol_00960 | NSD2_SET_2 | GAATTTGTTAACGAGTACGT |
| epikol_00961 | NSD2_SET_3 | ATTTGTTAACGAGTACGTTG |
| epikol_00962 | NSD2_SET_4 | AATTTGTTAACGAGTACGTT |
| epikol_00963 | NSD2_SET_5 | TGGGGCCAGCGTCTATTATA |
| epikol_00964 | NSD3_PHD_1 | ATGCAGATGAACTTGCTATC |
| epikol_00965 | NSD3_PHD_2 | ATGCCAATCCCAGGCACTCC |
| epikol_00966 | NSD3_PHD_3 | CAGGAAGTGATGCCAATCCC |
| epikol_00967 | NSD3_PHD_4 | GTGCTGCAAACACTTTCACC |
| epikol_00968 | NSD3_PHD_5 | GACTCTCTGATTCCTTGTGA |
| epikol_00969 | NSD3_SET_1 | CAAAACGGAGCGGAGAGGCT |
| epikol_00970 | NSD3_SET_2 | ATTAGTTACACTGTTCTCGT |
| epikol_00971 | NSD3_SET_3 | ATAATTGATGCCGGCCCAAA |
| epikol_00972 | NSD3_SET_4 | TAATTATAACCTAGATTGTC |
| epikol_00973 | NSD3_SET_5 | TTGGGCCGGCATCAATTATA |
| epikol_00974 | PADI4_PAD_1 | CACGCGTACACCTCCTGCGG |
| epikol_00975 | PADI4_PAD_2 | TGTGGTGTTCCAAGACAGCG |
| epikol_00976 | PADI4_PAD_3 | CTTGGAACACCACAGCCTCG |
| epikol_00977 | PADI4_PAD_4 | GGGAGGCCCTGCTGTTCGAA |
| epikol_00978 | PADI4_PAD_5 | CAGGGGCAAGGAATACCCGC |
| epikol_00979 | PARP1_PARP_1 | CGCATACTCCATCCTCAGTG |
| epikol_00980 | PARP1_PARP_2 | CCCAAAGTCGTGGGGGATCA |
| epikol_00981 | PARP1_PARP_3 | ATCTTTAAGATAGAGCGTGA |
| epikol_00982 | PARP1_PARP_4 | CTAATGTTAGCTGAAGGATC |
| epikol_00983 | PARP1_PARP_5 | TCCAGACTAATGTTAGCTGA |
| epikol_00984 | PAXIP1_BRCT_1 | GGGCTTTGGTTACGTTCTAT |
| epikol_00985 | PAXIP1_BRCT_2 | ATTTCTTATTGAGGGTTAGC |
| epikol_00986 | PAXIP1_BRCT_3 | GGCTTTGGTTACGTTCTATG |
| epikol_00987 | PAXIP1_BRCT_4 | AGAACGTAACCAAAGCCCAC |
| epikol_00988 | PAXIP1_BRCT_5 | TTTGATTGTTCCAGAGCCAA |
| epikol_00989 | PBRM1_BAH_1 | GGCCTGGTGCGTCCTCGTGT |
| epikol_00990 | PBRM1_BAH_2 | TCTTCTGGGTGAATGAAGAT |
| epikol_00991 | PBRM1_BAH_3 | TGGTATTTCAGTTGGCCTGC |
| epikol_00992 | PBRM1_BAH_4 | GTGTGCTGTGTTGTCATTCA |
| epikol_00993 | PBRM1_BAH_5 | GAAAAAGTATGGGTTCGAGA |
| epikol_00994 | PBRM1_BROMO_1 | TAATACCATCCGAGACTATA |
| epikol_00995 | PBRM1_BROMO_2 | CCGAGACTATAAGGATGAAC |
| epikol_00996 | PBRM1_BROMO_3 | TCTCTGTGAGCTCTTCATTA |
| epikol_00997 | PBRM1_BROMO_4 | ATTATAGAGTTCATGGCACA |
| epikol_00998 | PBRM1_BROMO_5 | CAAACTCATTTCTTGTTCGA |
| epikol_00999 | PCAF_BROMO_1 | TGAAGGGCCAAGCGCTTTGA |
| epikol_01000 | PCAF_BROMO_2 | CAGGATATTATGAAGTTATA |
| epikol_01001 | PCAF_BROMO_3 | ATTCTTGAGGCGTTCACTCA |
| epikol_01002 | PCAF_BROMO_4 | ACACGTAGTACCTATTCTTG |
| epikol_01003 | PCAF_BROMO_5 | GGGGTTGTACTCTTTGCAAT |
| epikol_01004 | PCAF_NAT_1 | ATTAAAGATGGCCGTGTTAT |

| | | |
|---|---|---|
| epikol_01005 | PCAF_NAT_2 | TCTCTGTGAATCCTTGAGAT |
| epikol_01006 | PCAF_NAT_3 | TTCCGTATGTTCCCATCTCA |
| epikol_01007 | PCAF_NAT_4 | AACAGATACCACCAATAACA |
| epikol_01008 | PCAF_NAT_5 | ATCTCTGTGAATCCTTGAGA |
| epikol_01009 | PCGF2_GECKO_1 | CTGCCCCATGTGTGACGTGC |
| epikol_01010 | PCGF2_GECKO_2 | TCAAGACATTGTCTACAAAT |
| epikol_01011 | PCGF2_GECKO_3 | CGATGAAGTACCCCCCGCAG |
| epikol_01012 | PCGF2_GECKO_4 | CATCGACGCCACCACTATCG |
| epikol_01013 | PCGF2_GECKO_5 | AACCTGCATCGTGCGCTACC |
| epikol_01014 | PCGF2_RING_1 | CATCGACGCCACCACTATCG |
| epikol_01015 | PCGF2_RING_2 | CGATGAAGTACCCCCCGCAG |
| epikol_01016 | PCGF2_RING_3 | CAGGCACTCCACGATAGTGG |
| epikol_01017 | PCGF2_RING_4 | AACCTGCATCGTGCGCTACC |
| epikol_01018 | PCGF2_RING_5 | GGCAGTATTTGTTGGTCTCC |
| epikol_01019 | PHF1_PHD_1 | CAGCTGACCAGCCGGTTCCC |
| epikol_01020 | PHF1_PHD_2 | ACTTCTCACAGCTGACCAGC |
| epikol_01021 | PHF1_PHD_3 | CGCTCTGAGACTGTGGTCCC |
| epikol_01022 | PHF1_PHD_4 | TGTCTGTCGCTCTGAGACTG |
| epikol_01023 | PHF1_PHD_5 | GCTCTGAGACTGTGGTCCCT |
| epikol_01024 | PHF1_TUDOR_1 | GACTGATGGGCTGCTATACT |
| epikol_01025 | PHF1_TUDOR_2 | TGCTGGCCAGATGGACTGAT |
| epikol_01026 | PHF1_TUDOR_3 | GTGCTGGCCAGATGGACTGA |
| epikol_01027 | PHF1_TUDOR_4 | GTCAAGATGTGCTGGCCAGA |
| epikol_01028 | PHF1_TUDOR_5 | ATTCGCAGTTTCTGGTTCTA |
| epikol_01029 | PHF2_GECKO_1 | GACGTCACTGACATAGAACG |
| epikol_01030 | PHF2_GECKO_2 | CCAAACTGTGAGAAAACCCA |
| epikol_01031 | PHF2_JMJC_1 | TGTCATGGGTAGAAAACTAC |
| epikol_01032 | PHF2_JMJC_2 | AGTTTCTTTACAATGTCAGG |
| epikol_01033 | PHF2_JMJC_3 | GTACTGCCTAATCTGCGTGA |
| epikol_01034 | PHF2_JMJC_4 | AACTGTCCTTCACGCAGATT |
| epikol_01035 | PHF2_JMJC_5 | CACGCAGATTAGGCAGTACT |
| epikol_01036 | PHF2_PHD_1 | TGAAGCGGGTAACGTCGTAG |
| epikol_01037 | PHF2_PHD_2 | GATCGAGTGCGACGCCTGCA |
| epikol_01038 | PHF2_PHD_3 | AGTGCGACGCCTGCAAGGAC |
| epikol_01039 | PHF8_JMJC_1 | AGCTATCTCGCACACTCATG |
| epikol_01040 | PHF8_JMJC_2 | GACAGCTTTCGAACAATCTT |
| epikol_01041 | PHF8_JMJC_3 | GATTGTTCGAAAGCTGTCAT |
| epikol_01042 | PHF8_JMJC_4 | CAATCTTCGGTGTCTCCACA |
| epikol_01043 | PHF8_JMJC_5 | TGTCATGGGTCGAAAACTTG |
| epikol_01044 | PHF8_PHD_1 | GGGTCACATCGTAAGGCAGC |
| epikol_01045 | PHF8_PHD_2 | GATCGAGTGTGACATGTGCC |
| epikol_01046 | PHF8_PHD_3 | CATCGTAAGGCAGCCGGCAG |
| epikol_01047 | PHF8_PHD_4 | ATGTGCCAGGACTGGTTTCA |
| epikol_01048 | PHF8_PHD_5 | TGTTGGTGTTGAAGAGGAGA |
| epikol_01049 | PPHLN1_1 | TGCTATTGCTTTAGAGCGAG |
| epikol_01050 | PPHLN1_2 | CTGTGAAACTTTCGGGATGG |
| epikol_01051 | PPHLN1_3 | AGACTGTGAAACTTTCGGGA |
| epikol_01052 | PPHLN1_4 | CGAAAGTTTCACAGTCTTGT |

| | | |
|---|---|---|
| epikol_01053 | PPHLN1_5 | GACAAGACTGTGAAACTTTC |
| epikol_01054 | PRDM1_SET_1 | CAGTTCCTAAGAACGCCAAC |
| epikol_01055 | PRDM1_SET_2 | CATTAAAGCCGTCAATGAAG |
| epikol_01056 | PRDM1_SET_3 | GGTCCAAAACGTGTGCCCTT |
| epikol_01057 | PRDM1_SET_4 | GGTGTAGATTTCACCTATTA |
| epikol_01058 | PRDM1_SET_5 | TGGTGTAGATTTCACCTATT |
| epikol_01059 | PRDM10_SET_1 | GCGCTTGGGGATGCGCCGCT |
| epikol_01060 | PRDM10_SET_2 | GGGGCCAAACTGGGTGCGCT |
| epikol_01061 | PRDM10_SET_3 | GGCCATACTGGTAAGCCACC |
| epikol_01062 | PRDM10_SET_4 | AATACACATGGTGGCCATAC |
| epikol_01063 | PRDM10_SET_5 | GGGATTTACATGAAGACCTA |
| epikol_01064 | PRDM11_SET_1 | CACATCTTCGGCCCCTATGA |
| epikol_01065 | PRDM11_SET_2 | TGTAAACGAGGTCATCCCCA |
| epikol_01066 | PRDM11_SET_3 | GGTGGAGATCTGCCCCTCAT |
| epikol_01067 | PRDM11_SET_4 | GGGCCGAAGATGTGGCCCTT |
| epikol_01068 | PRDM11_SET_5 | GCCAGCTGATTTGTCCTGGG |
| epikol_01069 | PRDM12_SET_1 | CGTGCTCCGGGGCGATCACG |
| epikol_01070 | PRDM12_SET_2 | CACCGGCCGCGTGATCGCCC |
| epikol_01071 | PRDM12_SET_3 | CTCCGGGGCGATCACGCGGC |
| epikol_01072 | PRDM12_SET_4 | TCCCGCCTTGATCCACGTCT |
| epikol_01073 | PRDM12_SET_5 | GATCAAGGCGGGAACCGAGA |
| epikol_01074 | PRDM13_SET_1 | GGCCGGCTTGCGCCTCGGAC |
| epikol_01075 | PRDM13_SET_2 | TTGCGCCTCGGACCGGTGCC |
| epikol_01076 | PRDM13_SET_3 | CACCGGTCCGAGGCGCAAGC |
| epikol_01077 | PRDM13_SET_4 | CAAGTACCTGTCAGACCGCA |
| epikol_01078 | PRDM13_SET_5 | GCTGGACGTCTCGCAATGCT |
| epikol_01079 | PRDM14_SET_1 | CATCACAGAATTGTCTCCGT |
| epikol_01080 | PRDM14_SET_2 | TCCGTAGGTCTTCACTTCAC |
| epikol_01081 | PRDM14_SET_3 | GAAGCGGGCACAGTTGACAT |
| epikol_01082 | PRDM14_SET_4 | GGGCCCAAACCTGACTCCTT |
| epikol_01083 | PRDM14_SET_5 | TCGCCAAAGGAGTCAGGTTT |
| epikol_01084 | PRDM15_SET_1 | GTCCAGGAGGGTCGCCAAAT |
| epikol_01085 | PRDM15_SET_2 | AGTCCAGGAGGGTCGCCAAA |
| epikol_01086 | PRDM15_SET_3 | TTCCCATTTGGCGACCCTCC |
| epikol_01087 | PRDM15_SET_4 | CCGCTTGACGAGCTGAGTGA |
| epikol_01088 | PRDM15_SET_5 | CGACTGGAAGATGGAGCCGA |
| epikol_01089 | PRDM16_SET_1 | AAAGGAGACAGACTTCGGAT |
| epikol_01090 | PRDM16_SET_2 | GTGGAAGTGTCGCCCCAGGA |
| epikol_01091 | PRDM16_SET_3 | GCGTGGATGCAAATCAGGCG |
| epikol_01092 | PRDM16_SET_4 | GTTCTGCGTGGATGCAAATC |
| epikol_01093 | PRDM16_SET_5 | GTCATTAAGGACATTGAGCC |
| epikol_01094 | PRDM2_SET_1 | TATGTGAATTGGGCTTGCTC |
| epikol_01095 | PRDM2_SET_2 | GGGTCTTGTCAACAGCAGAA |
| epikol_01096 | PRDM2_SET_3 | AAAAAATTTGGGCCATTTGT |
| epikol_01097 | PRDM2_SET_4 | ACTGGCTGCGATATGTGAAT |
| epikol_01098 | PRDM2_SET_5 | CTGGCTGCGATATGTGAATT |
| epikol_01099 | PRDM3_SET_1 | GAAAGACCCCAGTTATGGAT |
| epikol_01100 | PRDM3_SET_2 | GTCAACCAGATGTTGGAAGC |

| | | |
|---|---|---|
| epikol_01101 | PRDM3_SET_3 | GATGCCAGTCAACCAGATGT |
| epikol_01102 | PRDM3_SET_4 | CTCAAGTACATTAGATTCGC |
| epikol_01103 | PRDM3_SET_5 | GTAGTTGCAGACATTGCGCC |
| epikol_01104 | PRDM4_SET_1 | GGTCCAAAGCAAGTCCGCAC |
| epikol_01105 | PRDM4_SET_2 | ATTCCTGTGCGGACTTGCTT |
| epikol_01106 | PRDM4_SET_3 | ACTTGCTTTGGACCTCTAAT |
| epikol_01107 | PRDM4_SET_4 | ACAAGGCAGTTAACCATATC |
| epikol_01108 | PRDM4_SET_5 | TGACTCTGCTGGCCAATTAG |
| epikol_01109 | PRDM5_SET_1 | TGGATGAAAATATGGATTAC |
| epikol_01110 | PRDM5_SET_2 | GAAGCCAGTTGGAGTGCCGT |
| epikol_01111 | PRDM5_SET_3 | CTGGCTTCGCTTCGTTCATG |
| epikol_01112 | PRDM5_SET_4 | ATGAACGAAGCGAAGCCAGT |
| epikol_01113 | PRDM5_SET_5 | ACAGACACGGAGCTTCTGAT |
| epikol_01114 | PRDM6_SET_1 | GTCCAATCCAGGTGCCTTGC |
| epikol_01115 | PRDM6_SET_2 | ACCTGGATTGGACCTTTCCA |
| epikol_01116 | PRDM6_SET_3 | GGGAACCTAGTAAGTCGAGC |
| epikol_01117 | PRDM6_SET_4 | CTAGTAAGTCGAGCTGGATG |
| epikol_01118 | PRDM6_SET_5 | CTACAGCACTTTATTGATGG |
| epikol_01119 | PRDM8_SET_1 | AGCATTCTCAGGGATGTCGC |
| epikol_01120 | PRDM8_SET_2 | GAACAGTACCGTATATCTTT |
| epikol_01121 | PRDM8_SET_3 | CATATAGGGAAGTATGGCTC |
| epikol_01122 | PRDM8_SET_4 | TGTTCTACCGCTCTCTCCGC |
| epikol_01123 | PRDM8_SET_5 | AACTAGTAACTCCTCGTCTT |
| epikol_01124 | PRDM9_GECKO_1 | CAGAAGATTCTGATGAAGAA |
| epikol_01125 | PRDM9_GECKO_2 | GTTCTCTCTGTGTCTTCTTC |
| epikol_01126 | PRDM9_GECKO_3 | TCCCTCTTACCTTGCTGCCT |
| epikol_01127 | PRDM9_SET_1 | CAGCCAGGAGTATCCATTGT |
| epikol_01128 | PRDM9_SET_2 | TGGATGGAAAAGATAAATCC |
| epikol_01129 | PRMT1_SAM_1 | CCGGGGCCCGCAAGGTCATC |
| epikol_01130 | PRMT1_SAM_2 | CCCGATGACCTTGCGGGCCC |
| epikol_01131 | PRMT1_SAM_3 | TGCTCTATGCCCGGGACAAG |
| epikol_01132 | PRMT1_SAM_4 | CTTGTCCCGGGCATAGAGCA |
| epikol_01133 | PRMT1_SAM_5 | CAACACCGTGCTCTATGCCC |
| epikol_01134 | PRMT2_SAM_1 | TCTTCTGTGCACACTATGCG |
| epikol_01135 | PRMT2_SAM_2 | TGTCAGCAAAGCCGTTCTGC |
| epikol_01136 | PRMT2_SAM_3 | GCCCGAGAAGGTGGACGTGC |
| epikol_01137 | PRMT2_SAM_4 | CGAGTCCATCCTGTATGCCC |
| epikol_01138 | PRMT2_SAM_5 | TGAAGGAGGACGGGGTCATT |
| epikol_01139 | PRMT5_SAM_1 | ATACAGCTTTATCCGCCGGT |
| epikol_01140 | PRMT5_SAM_2 | CAGCATACAGCTTTATCCGC |
| epikol_01141 | PRMT5_SAM_3 | CCGCAGGGAAGCGTTCACCA |
| epikol_01142 | PRMT5_SAM_4 | CTGATGAGACTACGGTCACT |
| epikol_01143 | PRMT5_SAM_5 | CGTAGTCTCATCAGACATGA |
| epikol_01144 | PRMT6_SAM_1 | TCAGCCACTTGGTTCGCGCG |
| epikol_01145 | PRMT6_SAM_2 | CTCTACCGCGTACACGCGCC |
| epikol_01146 | PRMT6_SAM_3 | CCGGCGCGTGTACGCGGTAG |
| epikol_01147 | PRMT6_SAM_4 | CCTCTACCGCGTACACGCGC |
| epikol_01148 | PRMT6_SAM_5 | GGTGGTGCGGTTCAACGGGC |

| | | |
|---|---|---|
| epikol_01149 | PRMT7_SAM_1 | CACGGGACTCTTGTCAATGA |
| epikol_01150 | PRMT7_SAM_2 | CTCGATGGCATAGCAGAAGT |
| epikol_01151 | PRMT7_SAM_3 | TGGCTTTAGTGATAAGATTA |
| epikol_01152 | PRMT7_SAM_4 | TTCACAGCAGCATCAGCCAT |
| epikol_01153 | PRMT7_SAM_5 | TCAATGATGGCGGTCACAGC |
| epikol_01154 | PRMT8_SAM_1 | CCTTTCCATGTTCGCTGCCA |
| epikol_01155 | PRMT8_SAM_2 | CCCTGCCTTGGCAGCGAACA |
| epikol_01156 | PRMT8_SAM_3 | CCATGTTCGCTGCCAAGGCA |
| epikol_01157 | PRMT8_SAM_4 | TCAACACGGTGATCTTTGCC |
| epikol_01158 | PRMT8_SAM_5 | GATCATTAAGGCCAACCACT |
| epikol_01159 | RBBP4_GECKO_1 | CTACTCCATGAGTCTCTGTT |
| epikol_01160 | RBBP4_GECKO_2 | AGTAAGTGCCCACTGAGATT |
| epikol_01161 | RBBP4_GECKO_3 | AACCCAGACTTGCGTCTCCG |
| epikol_01162 | RBBP4_GECKO_4 | GACCTTGTCAGCTGATCCTG |
| epikol_01163 | RBBP4_GECKO_5 | CATTTAAGACCATCTGCCTG |
| epikol_01164 | RBBP5_WD40_1 | ATGATGGCCGAATTGTCATC |
| epikol_01165 | RBBP5_WD40_2 | TTTCAGGCGACTGTGACCAG |
| epikol_01166 | RBBP5_WD40_3 | AAATCGGAGTCATCGTCCAC |
| epikol_01167 | RBBP5_WD40_4 | GTCAATTGAGTTTGCCCGGA |
| epikol_01168 | RBBP5_WD40_5 | GATGGGGAATACATCGTGGC |
| epikol_01169 | RBX1_RING_1 | CTTCTGAAGTAGCGGACGCC |
| epikol_01170 | RBX1_RING_2 | AGAAGAGTGTACTGTCGCAT |
| epikol_01171 | RBX1_RING_3 | TCGCTGGCTCAAAACACGAC |
| epikol_01172 | RBX1_RING_4 | AGTACACTCTTCTGAAGTAG |
| epikol_01173 | RBX1_RING_5 | ACTTCCACTGCATCTCTCGC |
| epikol_01174 | RNF168_RING_1 | TCGACGGTCGACTGGAAGCA |
| epikol_01175 | RNF168_RING_2 | CTTCCAGTCGACCGTCGAAA |
| epikol_01176 | RNF168_RING_3 | CAGCGTGTGGTTACACGGGA |
| epikol_01177 | RNF168_RING_4 | AGCACGGTTTACACAGCGTG |
| epikol_01178 | RNF168_RING_5 | GATCTGCATGGAAATCCTCG |
| epikol_01179 | RNF2_GECKO_2 | AATTCACTGTGTAGACTTCG |
| epikol_01180 | RNF2_GECKO_3 | ATTATTGTGCTTGTTGATCC |
| epikol_01181 | RNF2_GECKO_4 | TTTCAACATACCCACTTCTA |
| epikol_01182 | RNF2_GECKO_5 | TGAGTTACAACGAACACCTC |
| epikol_01183 | RNF2_RING_1 | TCATCACAGCCCTTAGAAGT |
| epikol_01184 | RNF20_RING_1 | ACGCATGTTACAGCACGGAC |
| epikol_01185 | RNF20_RING_2 | TACACTTGGGACATTTGCGC |
| epikol_01186 | RNF20_RING_3 | CTTGGGACATTTGCGCTGGC |
| epikol_01187 | RNF20_RING_4 | GTGCTGTAACATGCGTAAAA |
| epikol_01188 | RNF20_RING_5 | ACTTGGGACATTTGCGCTGG |
| epikol_01189 | RNF8_RING_1 | AGAATGCCCCATTTGTCGGA |
| epikol_01190 | RNF8_RING_2 | GCTCCTACTGTATCAATGAA |
| epikol_01191 | RNF8_RING_3 | GAAACTGTGGGCACAGTTCA |
| epikol_01192 | RNF8_RING_4 | ACAGTAGGAGCAGAAACTGT |
| epikol_01193 | RNF8_RING_5 | AGATAGAATGCCCCATTTGT |
| epikol_01194 | SATB1_1 | GCAATGCCATTTCGATCAGC |
| epikol_01195 | SATB1_2 | CGCCATTGAATATGATTGCA |
| epikol_01196 | SATB1_3 | GGAGCATGCAGAATTTGTGC |

| | | |
|---|---|---|
| epikol_01197 | SATB1_4 | TGTTCCACCACACAGAAAAC |
| epikol_01198 | SATB1_5 | AGAATTTGTGCTGGTGAGAA |
| epikol_01199 | SETD1A_SET_1 | CCGGAGCCGGATCCACGAGT |
| epikol_01200 | SETD1A_SET_2 | GGATCCGGCTCCGGCCAAAT |
| epikol_01201 | SETD1A_SET_3 | TGTCGTGGTCCACCCGGAAC |
| epikol_01202 | SETD1A_SET_4 | GCGGGAGAAGCGCTACGTGC |
| epikol_01203 | SETD1A_SET_5 | ATCTCCTCGTCCACGCCAAT |
| epikol_01204 | SETD2_SET_1 | GTCATACTCACAGAAAAGAA |
| epikol_01205 | SETD2_SET_2 | AGAGTTTAAAGCTCGAGTGA |
| epikol_01206 | SETD2_SET_3 | ACCTTTGTCCTAGAATATTG |
| epikol_01207 | SETD2_SET_4 | GGACTGTGAACGGACAACTG |
| epikol_01208 | SETD2_SET_5 | AACGTTAACTCTGAGCCTGA |
| epikol_01209 | SETD3_SET1_1 | GGCCAGTGCGATGTTTCCCA |
| epikol_01210 | SETD3_SET1_2 | AGCCATGGGAAACATCGCAC |
| epikol_01211 | SETD3_SET1_3 | GACCGAATCCTTCAAGCCAT |
| epikol_01212 | SETD3_SET1_4 | CGATGTTTCCCATGGCTTGA |
| epikol_01213 | SETD3_SET1_5 | GAGCCAGCCCTAACTCCTTC |
| epikol_01214 | SETD3_SET2_1 | GAAGGATTCTTTCACTTACG |
| epikol_01215 | SETD3_SET2_2 | TGCCAACAAACTACCCTTGA |
| epikol_01216 | SETD3_SET2_3 | GGATGGTTCCCGCGTGACCC |
| epikol_01217 | SETD3_SET2_4 | CAGTCTCTTCTGTTATGACG |
| epikol_01218 | SETD3_SET2_5 | ACGCGGGAACCATCCTCTGT |
| epikol_01219 | SETD4_SET_1 | CTGTGCACTCGCTCCGTACC |
| epikol_01220 | SETD4_SET_2 | GGGCCTCAGGTACACGGCTC |
| epikol_01221 | SETD4_SET_3 | GAAAGGCATTCCCGCTGCCT |
| epikol_01222 | SETD4_SET_4 | AAATTAGAACGACTTCACGT |
| epikol_01223 | SETD4_SET_5 | GAAGAGGTATTCATCTGTTA |
| epikol_01224 | SETD6_SET_1 | GTTCTAGGGAGCGAACCCTG |
| epikol_01225 | SETD6_SET_2 | TGAAGGGCAGCACGATGGAC |
| epikol_01226 | SETD6_SET_3 | TTCTAGATTGGCGTTGTGAT |
| epikol_01227 | SETD6_SET_4 | GGCACCATCACGGGGGAGTT |
| epikol_01228 | SETD6_SET_5 | ATCTCATGGCCTTTAGGAAT |
| epikol_01229 | SETD7_SET_1 | GATACGTGGTTATAGGGCTC |
| epikol_01230 | SETD7_SET_2 | AGGCACAGTACTTGGATACG |
| epikol_01231 | SETD7_SET_3 | TCCCAAGGAGGCACAGTACT |
| epikol_01232 | SETD7_SET_4 | CATTTGATGGGCCCAAAACG |
| epikol_01233 | SETD7_SET_5 | GAAGAGCTCACCGTTGCCTA |
| epikol_01234 | SETD8_GECKO_1 | TGGGCAAATACCTCTAGCCA |
| epikol_01235 | SETD8_GECKO_2 | AGGTACCAGGGGACCTTTCC |
| epikol_01236 | SETD8_GECKO_3 | TAACTGAGTTCTCTTCCTGA |
| epikol_01237 | SETD8_GECKO_4 | ACGGAGCGCCATGAAGTCCG |
| epikol_01238 | SETD8_GECKO_5 | GCCTGCTTACCCCGTCGGTG |
| epikol_01239 | SETDB1_SET1_1 | GGGCTGGGGTATCCGCTGCT |
| epikol_01240 | SETDB1_SET1_2 | TGCTTGGATGACATTGCCAA |
| epikol_01241 | SETDB1_SET1_3 | TCTGGACCATATCGAGAGCG |
| epikol_01242 | SETDB1_SET1_4 | AGTTCTCCACGCTCTCGATA |
| epikol_01243 | SETDB1_SET1_5 | CTTTGCAGACAAGGAGGGTC |
| epikol_01244 | SETDB1_SET2_1 | ATCATTGATGCCAAGCTTGA |

| | | |
|---|---|---|
| epikol_01245 | SETDB1_SET2_2 | GGGTATCCACGAAGACATTC |
| epikol_01246 | SETDB1_SET2_3 | GTTTGTCCAGAATGTCTTCG |
| epikol_01247 | SETDB1_SET2_4 | GGGCTGGGACAGAACTTACT |
| epikol_01248 | SETDB1_SET2_5 | GCCAAGCTTGAAGGCAACCT |
| epikol_01249 | SETDB2_SET1_1 | TGTCTAGATGACATTGACAG |
| epikol_01250 | SETDB2_SET1_2 | GGTGTTCAAAACTGAGCAGA |
| epikol_01251 | SETDB2_SET1_3 | CTTATGGTATTGATGAAAAC |
| epikol_01252 | SETDB2_SET1_4 | AAAACTGAGCAGAAGGGATG |
| epikol_01253 | SETDB2_SET1_5 | GCTAACACTGAAAAATCTTA |
| epikol_01254 | SETDB2_SET2_1 | TTATTGGATGCCACAAAAGA |
| epikol_01255 | SETDB2_SET2_2 | TTGGTGAAGAATGCCACCAA |
| epikol_01256 | SETDB2_SET2_3 | ACATTCTGTACCAAGAGATT |
| epikol_01257 | SETDB2_SET2_4 | AAGCAAGAACAGAGCTAACA |
| epikol_01258 | SETDB2_SET2_5 | ACAGAGCTAACATGGGATTA |
| epikol_01259 | SETMAR_SET_1 | ACCATTGAGTCAATTCGGAC |
| epikol_01260 | SETMAR_SET_2 | AGGCTGGGGACTTCGTACCT |
| epikol_01261 | SETMAR_SET_3 | TGACTCAATGGTACCTAAGT |
| epikol_01262 | SETMAR_SET_4 | GCGTCTTGAACACTTGGAAG |
| epikol_01263 | SETMAR_SET_5 | CCAATTACATTATAGCCATC |
| epikol_01264 | SFMBT1_1 | GGTCCACAGCAGTTCCCTAT |
| epikol_01265 | SFMBT1_2 | CTCACAGGTAGTGATAACGG |
| epikol_01266 | SFMBT1_3 | CATAGCCATCATAGCGGAGA |
| epikol_01267 | SFMBT1_4 | AGGCTGATCTCTACCCCATT |
| epikol_01268 | SFMBT1_5 | GTGGACACACGTTTGCAAAA |
| epikol_01269 | SFMBT2_1 | TCTGCGCTACTGCGGTTACG |
| epikol_01270 | SFMBT2_2 | CTGGTGTGACGTAGTCATCG |
| epikol_01271 | SFMBT2_3 | CCCGCACGTGGTAATGATCG |
| epikol_01272 | SFMBT2_4 | CATCGCGGATTTGCACCCCG |
| epikol_01273 | SFMBT2_5 | GAACAACCCGGACACGTACT |
| epikol_01274 | SHPRH_H15_1 | GATACGATGTTCAACGGAAC |
| epikol_01275 | SHPRH_H15_2 | GGTCTCGTGAAACAGATCAA |
| epikol_01276 | SHPRH_H15_3 | AGAACTGACATACTTATAGA |
| epikol_01277 | SHPRH_H15_4 | ATACTTATAGATGGAAAGGA |
| epikol_01278 | SHPRH_H15_5 | GTGAAACAGATCAAAGGCCA |
| epikol_01279 | SHPRH_HELIC_1 | CGTTGAGAAAACGAGTGCTT |
| epikol_01280 | SHPRH_HELIC_2 | AGCTCTTACTGACAACAACA |
| epikol_01281 | SHPRH_HELIC_3 | ATAGGGAGGGTGCACCGAAT |
| epikol_01282 | SHPRH_HELIC_4 | TCATGGGCAGGGTTCAATAT |
| epikol_01283 | SHPRH_HELIC_5 | ATCCATTAGAACCTGTGTGC |
| epikol_01284 | SIN3A_1 | CTTTGGTAAGGCTCGATAGC |
| epikol_01285 | SIN3A_2 | AGGAGTCCGTCCTGTACACT |
| epikol_01286 | SIN3A_3 | TCTGGCAACAATCCGGGTTC |
| epikol_01287 | SIN3A_4 | AGACCAATCTGGCAACAATC |
| epikol_01288 | SIN3A_5 | CTGCTGGTAACTCTTTGGTA |
| epikol_01289 | SIN3B_1 | GCTGGATCCTATGCGCTTGC |
| epikol_01290 | SIN3B_2 | TTTGGGGAGTGCCCGGTAGC |
| epikol_01291 | SIN3B_3 | AATCGTCTGTCCACGAACCC |
| epikol_01292 | SIN3B_4 | TGGATTCCAGGATATAGTGC |

| | | |
|---|---|---|
| epikol_01293 | SIN3B_5 | CTTCTTGGAGCTGACGAACG |
| epikol_01294 | SIRT1_SIRT_1 | TCAAACATCGCTTGAGGATC |
| epikol_01295 | SIRT1_SIRT_2 | CCATCCCTTGACCTGAAGTC |
| epikol_01296 | SIRT1_SIRT_3 | TACCCAGAACATAGACACGC |
| epikol_01297 | SIRT1_SIRT_4 | AACAGGTTGCGGGAATCCAA |
| epikol_01298 | SIRT1_SIRT_5 | ATAGCAAGCGGTTCATCAGC |
| epikol_01299 | SIRT2_SIRT_1 | TAGGTTGTCATAGAGGCCGG |
| epikol_01300 | SIRT2_SIRT_2 | TTCCTTGGCGAGGGCGAAGA |
| epikol_01301 | SIRT2_SIRT_3 | CTACTTCATGCGCCTGCTGA |
| epikol_01302 | SIRT2_SIRT_4 | CCAGCCCGGCTATTCGCTCC |
| epikol_01303 | SIRT2_SIRT_5 | GGCACGAATACCCGCTAAGC |
| epikol_01304 | SIRT3_SIRT_1 | GGTACGGGAGATCGTACTGC |
| epikol_01305 | SIRT3_SIRT_2 | CTCTACACGCAGAACATCGA |
| epikol_01306 | SIRT3_SIRT_3 | CAAAAATGGCCTCGGGGTAC |
| epikol_01307 | SIRT3_SIRT_4 | ATGAGCTTCAACCAGCTTTG |
| epikol_01308 | SIRT3_SIRT_5 | GGGTCTTTGGCAGACTGTGC |
| epikol_01309 | SIRT4_SIRT_1 | GTCTGGTATCCCCGATTCGG |
| epikol_01310 | SIRT4_SIRT_2 | AGTTTCTCGCCCAGTACCGC |
| epikol_01311 | SIRT4_SIRT_3 | TTTATGCCCGCACTGACCGC |
| epikol_01312 | SIRT4_SIRT_4 | TACTGGGCGAGAAACTTCGT |
| epikol_01313 | SIRT4_SIRT_5 | GTCCCAACCTGCGTTCAATG |
| epikol_01314 | SIRT5_SIRT_1 | AGTGGTGTTCCGACCTTCAG |
| epikol_01315 | SIRT5_SIRT_2 | GGGTGATGACCACGACTCGC |
| epikol_01316 | SIRT5_SIRT_3 | GTTCTACCACTACCGGCGGG |
| epikol_01317 | SIRT5_SIRT_4 | TATGGCGCGGTGCCCGGCGT |
| epikol_01318 | SIRT5_SIRT_5 | GGGAGTTCTACCACTACCGG |
| epikol_01319 | SIRT6_SIRT_1 | GTGGTGTTCCACACGGGTGC |
| epikol_01320 | SIRT6_SIRT_2 | CGTGGGCCGCGCGCTCTCAA |
| epikol_01321 | SIRT6_SIRT_3 | CTCAAAGGTGGTGTCGAACT |
| epikol_01322 | SIRT6_SIRT_4 | CATGGAGCCCGTCCACGTTC |
| epikol_01323 | SIRT6_SIRT_5 | CTTCCACAAACATGTTCCCG |
| epikol_01324 | SIRT7_SIRT_1 | GCGTCTATCCCAGACTACCG |
| epikol_01325 | SIRT7_SIRT_2 | GGGTTGGCTCGGCCTCGCTC |
| epikol_01326 | SIRT7_SIRT_3 | CAGACGGGTGATGCTCATGT |
| epikol_01327 | SIRT7_SIRT_4 | GTACATGTTCCCGTGGAGCT |
| epikol_01328 | SIRT7_SIRT_5 | AGCTCGGAGATGGCCGTGCG |
| epikol_01329 | SMARCA1_DEXD_1 | GAATATTTCGGTAGTGTTTC |
| epikol_01330 | SMARCA1_DEXD_2 | AAAGTTTCACTGGCGATACC |
| epikol_01331 | SMARCA1_DEXD_3 | ATTCGTGATGAAATGATGCC |
| epikol_01332 | SMARCA1_DEXD_4 | ACGCAAACATCCCACTCTCC |
| epikol_01333 | SMARCA1_DEXD_5 | ACTAACCGCTTGCTCCTAAC |
| epikol_01334 | SMARCA1_HELIC_1 | GGTAGTTCTGGATAAACTAT |
| epikol_01335 | SMARCA1_HELIC_2 | TGGTTATGAGTATTGTCGAC |
| epikol_01336 | SMARCA1_HELIC_3 | CAGCCAGATGACTCGCTTGC |
| epikol_01337 | SMARCA1_HELIC_4 | GAAGATTATTGCATGTGGCG |
| epikol_01338 | SMARCA1_HELIC_5 | TGGAGGTCTCGGAATTAACC |
| epikol_01339 | SMARCA2_DEXD_1 | CGGAATCTTAGCCGATGAAA |
| epikol_01340 | SMARCA2_DEXD_2 | GATGAGTGCAATGGTCTGTA |

| | | |
|---|---|---|
| epikol_01341 | SMARCA2_DEXD_3 | TGACAAATGGGCTCCTTCTG |
| epikol_01342 | SMARCA2_DEXD_4 | ACAAGGGAGCGACGCATGGC |
| epikol_01343 | SMARCA2_DEXD_5 | TGAATTTGCCACTCCGTAGC |
| epikol_01344 | SMARCA2_HELIC_1 | GCTGAACTGTATCGGGCCTC |
| epikol_01345 | SMARCA2_HELIC_2 | AGCAGCACGATCTTCAGACT |
| epikol_01346 | SMARCA2_HELIC_3 | TGGCCTGGGCTTAAATCTTC |
| epikol_01347 | SMARCA2_HELIC_4 | CGATGCGGTGAGCTCGGTCT |
| epikol_01348 | SMARCA2_HELIC_5 | TTCTTGCTGAGCACAAGAGC |
| epikol_01349 | SMARCA4_DEXD_1 | GAGGTACGTGATGAGCGCGA |
| epikol_01350 | SMARCA4_DEXD_2 | GGCATCCTGGCCGACGAGAT |
| epikol_01351 | SMARCA4_DEXD_3 | GTCAAACTCGTACGCCCAGT |
| epikol_01352 | SMARCA4_DEXD_4 | TGACAAGTGGGCCCCCTCCG |
| epikol_01353 | SMARCA4_DEXD_5 | ACAAAGGCCCGTCTTGCTGC |
| epikol_01354 | SMARCA4_HELIC_1 | GAAGATTACTTTGCGTATCG |
| epikol_01355 | SMARCA4_HELIC_2 | ATCGCGGCTTTAAATACCTC |
| epikol_01356 | SMARCA4_HELIC_3 | GTGGTTGGTTGCTCGGAGTT |
| epikol_01357 | SMARCA4_HELIC_4 | CTCAGAGCCGGGCTCGTTGA |
| epikol_01358 | SMARCA4_HELIC_5 | ACCACGAAGGCGGAGGACCG |
| epikol_01359 | SMARCA5_DEXD_1 | ACCAAAACCATATGAGGCCC |
| epikol_01360 | SMARCA5_DEXD_2 | TAGTAATAGATGAAGCTCAC |
| epikol_01361 | SMARCA5_DEXD_3 | TTAGGAACCAAAACCATATG |
| epikol_01362 | SMARCA5_DEXD_4 | ACACATACATCCCATTCTCC |
| epikol_01363 | SMARCA5_DEXD_5 | GATGAGTGAATTCAAGAGAT |
| epikol_01364 | SMARCA5_HELIC_1 | ATGCATCTAGTAACCAACAG |
| epikol_01365 | SMARCA5_HELIC_2 | AAATTATGAGTACTGCAGGT |
| epikol_01366 | SMARCA5_HELIC_3 | CAGTCAAATGACAAGGGTAT |
| epikol_01367 | SMARCA5_HELIC_4 | TTCATGTTAAGCACGCGTGC |
| epikol_01368 | SMARCA5_HELIC_5 | TCTGCCCAATTCTATGTGCT |
| epikol_01369 | SMARCAD1_DEXD_1 | CTGGCATACCTCTATCAGGA |
| epikol_01370 | SMARCAD1_DEXD_2 | CATTATTACCCTCCTGATAG |
| epikol_01371 | SMARCAD1_DEXD_3 | TTTATGGTGCCCTACTTTGA |
| epikol_01372 | SMARCAD1_DEXD_4 | ATGACCGTAGTCTGTTTCGA |
| epikol_01373 | SMARCAD1_DEXD_5 | CATAAGGTGCTGGTAGCGAA |
| epikol_01374 | SMARCAD1_HELIC_1 | GAGTCTTTCCATCTAATCTG |
| epikol_01375 | SMARCAD1_HELIC_2 | TAGCCAATTTACCATGATGC |
| epikol_01376 | SMARCAD1_HELIC_3 | CCATAGAGTAGGCCAGACTA |
| epikol_01377 | SMARCAD1_HELIC_4 | TGATGAGTTTAATACCGATA |
| epikol_01378 | SMARCAD1_HELIC_5 | TCAACAAAAGCTGGTGGATT |
| epikol_01379 | SMARCAL1_DEXD_1 | CGGCCACTCCTTCCGGTAAA |
| epikol_01380 | SMARCAL1_DEXD_2 | CTCCCAGGTGAAGCGCACGG |
| epikol_01381 | SMARCAL1_DEXD_3 | CCCAGATTGCATCAACGTCG |
| epikol_01382 | SMARCAL1_DEXD_4 | TGCATCAACGTCGTGGTGAC |
| epikol_01383 | SMARCAL1_DEXD_5 | GGGAAGGACCGCCTGACAGC |
| epikol_01384 | SMARCAL1_HELIC_1 | GCTCTTGCGTAATTGCGTCC |
| epikol_01385 | SMARCAL1_HELIC_2 | TGCACACCATAAGGTGGTCC |
| epikol_01386 | SMARCAL1_HELIC_3 | GTGCAGCACATCCGCATCGA |
| epikol_01387 | SMARCAL1_HELIC_4 | TCCATCACCGCTGCCAATAT |
| epikol_01388 | SMARCAL1_HELIC_5 | AGCAGTTCCAACTGTCGGAG |

| | | |
|---|---|---|
| epikol_01389 | SMARCC2_CHROM_1 | GGGTACGACCAGTCATGAAG |
| epikol_01390 | SMARCC2_CHROM_2 | ATAAGCAGGTTCTTCTGCAC |
| epikol_01391 | SMARCC2_CHROM_3 | GGTACGACCAGTCATGAAGA |
| epikol_01392 | SMARCC2_CHROM_4 | TGCTTATCCCTCTTCATGAC |
| epikol_01393 | SMARCC2_CHROM_5 | GATCCCAGCGAGTGAAATTG |
| epikol_01394 | SMARCD1_RSC6_1 | GATGGTTGTCTGGCCCATAC |
| epikol_01395 | SMARCD1_RSC6_2 | GTATGGGCCAGACAACCATC |
| epikol_01396 | SMARCD1_RSC6_3 | GGCCGGGAGACGTGAATGTA |
| epikol_01397 | SMARCD1_RSC6_4 | GCACAGGACCGCCACTACCC |
| epikol_01398 | SMARCD1_RSC6_5 | GTGTACTGTCCTACTGATGC |
| epikol_01399 | SMARCD3_RSC6_1 | TGCCTCCTGGGAGCTACGGG |
| epikol_01400 | SMARCD3_RSC6_2 | CAAGAGTTTGGTCATCGAGC |
| epikol_01401 | SMARCD3_RSC6_3 | GCATCGGACACCCACGACCC |
| epikol_01402 | SMARCD3_RSC6_4 | GAGCCGCTCAGCCATTGTCC |
| epikol_01403 | SMARCD3_RSC6_5 | GGCGGGGATCCAGTTTGAAC |
| epikol_01404 | SMARCE1_GECKO_1 | AAACTTACTGTTGCAGGAGC |
| epikol_01405 | SMARCE1_GECKO_2 | TTACCATCTGGATCTTCAGC |
| epikol_01406 | SMARCE1_GECKO_3 | GTGGGAGGTGGGGCATAAGA |
| epikol_01407 | SMARCE1_GECKO_4 | TCGACAGAGACAATCTCGCA |
| epikol_01408 | SMARCE1_GECKO_5 | AGACGACGAGAACATTCCGA |
| epikol_01409 | SMYD1_SET_1 | GCCAACAGTCATGGTTCACC |
| epikol_01410 | SMYD1_SET_2 | AGAATTGAGCTCCGGGCCCT |
| epikol_01411 | SMYD1_SET_3 | GCCCTAGGCAAGATCTCAGA |
| epikol_01412 | SMYD1_SET_4 | TACAGTTGGGCCAACAGTCA |
| epikol_01413 | SMYD1_SET_5 | CGTAGGCATCTTCCCCAACC |
| epikol_01414 | SMYD2_SET_1 | AGATGAAGAACTTTCTCATT |
| epikol_01415 | SMYD2_SET_2 | TGTGACCTACAAAGGGACCC |
| epikol_01416 | SMYD2_SET_3 | TTCTGCCAGGGTCCCTTTGT |
| epikol_01417 | SMYD2_SET_4 | GCTGTACAGGAAATCAAGCC |
| epikol_01418 | SMYD2_SET_5 | GGCAGAAGTCAGAGCTGTAC |
| epikol_01419 | SMYD3_SET_1 | GCTTCAAAAAGGTCAAAGGC |
| epikol_01420 | SMYD3_SET_2 | GCGAGCAGTCCGAGACATCG |
| epikol_01421 | SMYD3_SET_3 | GACTGCTCGCAGTAAGAGGT |
| epikol_01422 | SMYD3_SET_4 | TGAACACAATCGAACAGTTG |
| epikol_01423 | SMYD3_SET_5 | ATTGAACACAATCGAACAGT |
| epikol_01424 | SMYD4_SET_1 | ACAGCTTCAGTGTAACGCTC |
| epikol_01425 | SMYD4_SET_2 | TTCTAATCCGCTGTGACGCC |
| epikol_01426 | SMYD4_SET_3 | GGAGCATCGTTACCGACAGC |
| epikol_01427 | SMYD4_SET_4 | AATCCGCTGTGACGCCCGGA |
| epikol_01428 | SMYD4_SET_5 | AGTGCTAATGAAGGACACGC |
| epikol_01429 | SMYD5_SET1_1 | CAGGGGCCGTTCTACGAAGA |
| epikol_01430 | SMYD5_SET1_2 | GGGAGACCATCTTCGTAGAA |
| epikol_01431 | SMYD5_SET1_3 | TGCCACACAGCTCATCCGGA |
| epikol_01432 | SMYD5_SET1_4 | CATCTTCGTAGAACGGCCCC |
| epikol_01433 | SMYD5_SET1_5 | CGAAGATGGTCTCCCCCTTC |
| epikol_01434 | SMYD5_SET2_1 | TTTCTTAACTGTGAAGGATC |
| epikol_01435 | SMYD5_SET2_2 | GCTCTGGAGGATATTAAGCC |
| epikol_01436 | SMYD5_SET2_3 | CTGCATTGGGCACACAACTG |

| | | |
|---|---|---|
| epikol_01437 | SMYD5_SET2_4 | GGAAAGGAGGTCTCTGCATT |
| epikol_01438 | SMYD5_SET2_5 | TGGAAAGGAGGTCTCTGCAT |
| epikol_01439 | SP100_BROMO_1 | AATCACAGTAGACCTTCAAG |
| epikol_01440 | SP100_BROMO_2 | CAGATGTACACCCGAGTAGA |
| epikol_01441 | SP100_BROMO_3 | CTGCACAAACCCTTCTACTC |
| epikol_01442 | SP100_BROMO_4 | TTAACCACATGGGCTTCTGT |
| epikol_01443 | SP100_BROMO_5 | TTGACTTTGTTTAACCACAT |
| epikol_01444 | SRCAP_DEXD_1 | CATCTCTCTGCTTGCCCACT |
| epikol_01445 | SRCAP_DEXD_2 | CAGTTCAACATCACGCTGGT |
| epikol_01446 | SRCAP_DEXD_3 | CCACCAGCGTGATGTTGAAC |
| epikol_01447 | SRCAP_DEXD_4 | GCGCCAGTTCTTGCGACGGA |
| epikol_01448 | SRCAP_DEXD_5 | GATAGCGCCAGTTCTTGCGA |
| epikol_01449 | SRCAP_HELIC_1 | CATCCAGCATTCGGGTCATC |
| epikol_01450 | SRCAP_HELIC_2 | TAGTAGATCCATCCAGGCGC |
| epikol_01451 | SRCAP_HELIC_3 | CACCCAGATGACCCGAATGC |
| epikol_01452 | SRCAP_HELIC_4 | CAGGACCGCTGTCACCGAAT |
| epikol_01453 | SRCAP_HELIC_5 | TCATCCTTTCAACTCGGAGT |
| epikol_01454 | SSRP1_PHD1_1 | CGAATGTCATAACGACCACG |
| epikol_01455 | SSRP1_PHD1_2 | GACGCAGTACTGTGGTGTAG |
| epikol_01456 | SSRP1_PHD1_3 | CTCGTGGTCGTTATGACATT |
| epikol_01457 | SSRP1_PHD1_4 | TCAGGAAGTGGTAGCGAGTT |
| epikol_01458 | SSRP1_PHD1_5 | CGAGTTTGGCCTTGCTTGAT |
| epikol_01459 | SSRP1_PHD2_1 | CTCAGGACTGCTCTACCCGC |
| epikol_01460 | SSRP1_PHD2_2 | TCATCGAAGCGGATGTGCAC |
| epikol_01461 | SSRP1_PHD2_3 | GTCAAAGGAACGAGTAGTAG |
| epikol_01462 | SSRP1_PHD2_4 | GACTGCTCTACCCGCTGGAG |
| epikol_01463 | SSRP1_PHD2_5 | TCCTTTGTCAACTTTGCTCG |
| epikol_01464 | SUPT16H_1 | GAAACTGGACCCGCTCAAAG |
| epikol_01465 | SUPT16H_2 | TACAGGAATGGCGTTGATCA |
| epikol_01466 | SUPT16H_3 | ATGGGGTCAAGAGAGGCTAC |
| epikol_01467 | SUPT16H_4 | CTCTTGACCCCATCAAGGAA |
| epikol_01468 | SUPT16H_5 | CGCTGGTAAATGCTACGGAA |
| epikol_01469 | SUPT16H_FACT_1 | TCCTTGACAAAAGTCGCTTC |
| epikol_01470 | SUPT16H_FACT_2 | GCATCAAATATTAAGGCACC |
| epikol_01471 | SUPT16H_FACT_3 | GGCTGGTACTGTCTGTTCTC |
| epikol_01472 | SUPT16H_FACT_4 | ATACCGAGCATCAAATATTA |
| epikol_01473 | SUPT16H_FACT_5 | GCTGGTACTGTCTGTTCTCC |
| epikol_01474 | SUV39H1_CHROM_1 | GGATCTTCTTGTAATCGCAC |
| epikol_01475 | SUV39H1_CHROM_2 | ACACACTTGAGATTCTGCCG |
| epikol_01476 | SUV39H1_CHROM_3 | TATTACCTGGTGAAATGGCG |
| epikol_01477 | SUV39H1_CHROM_4 | GATATCCACGCCATTTCACC |
| epikol_01478 | SUV39H1_CHROM_5 | GAGATTCTGCCGTGGCTCCC |
| epikol_01479 | SUV39H1_SET_1 | AGCTTCGTCATGGAGTACGT |
| epikol_01480 | SUV39H1_SET_2 | TTCCGCACGGATGATGGGCG |
| epikol_01481 | SUV39H1_SET_3 | CGTGGAGGACGTGTACACCG |
| epikol_01482 | SUV39H1_SET_4 | GGGCCAGATCTACGACCGTC |
| epikol_01483 | SUV39H1_SET_5 | CGCCCTGACGGTCGTAGATC |
| epikol_01484 | SUV39H2_CHROM_1 | GGAATACTTGTGTGACTACA |

| | | |
|---|---|---|
| epikol_01485 | SUV39H2_CHROM_2 | GGCCAGATTCTACAAATACT |
| epikol_01486 | SUV39H2_CHROM_3 | ATCTTGTAAAATGGAAAGGA |
| epikol_01487 | SUV39H2_CHROM_4 | TGGAATATTATCTTGTAAAA |
| epikol_01488 | SUV39H2_CHROM_5 | TATTATCTTGTAAAATGGAA |
| epikol_01489 | SUV39H2_SET_1 | TTTCGAACTAGCAATGGACG |
| epikol_01490 | SUV39H2_SET_2 | AACTAGCAATGGACGTGGCT |
| epikol_01491 | SUV39H2_SET_3 | TGCATCTTTCGAACTAGCAA |
| epikol_01492 | SUV39H2_SET_4 | ACAGTGGATGCGGCTCGATA |
| epikol_01493 | SUV39H2_SET_5 | AATCACGTATCTCTTTGATC |
| epikol_01494 | SUV420H1_SET_1 | TCAGTGTCATGTACTCCACA |
| epikol_01495 | SUV420H1_SET_2 | GAGAACATGCTACTTAGACA |
| epikol_01496 | SUV420H1_SET_3 | AAACTGTGCTCAACTCTGGC |
| epikol_01497 | SUV420H1_SET_4 | GCTCTAAGAGACATTGAACC |
| epikol_01498 | SUV420H1_SET_5 | TCGAGATACAGCATGTGTGA |
| epikol_01499 | SUV420H2_SET_1 | TCATGTACTCAACCCGCAAG |
| epikol_01500 | SUV420H2_SET_2 | CAGCTGAGCACTCCGCTTGC |
| epikol_01501 | SUV420H2_SET_3 | GCAAGCGGAGTGCTCAGCTG |
| epikol_01502 | SUV420H2_SET_4 | GTGACATGCTTCTACGGCGA |
| epikol_01503 | SUV420H2_SET_5 | GTCCCGGAGCACCTTCACGC |
| epikol_01504 | TAF1_BROMO_1 | CAAAGTGATTGTCAATCCAA |
| epikol_01505 | TAF1_BROMO_2 | GGATGATGTAAACCTTATTC |
| epikol_01506 | TAF1_BROMO_3 | AACAGACGTTCACAATCTCC |
| epikol_01507 | TAF1_GECKO_1 | AAGAATCACCTGAGTGGCAA |
| epikol_01508 | TAF1_GECKO_2 | GATAAAATCTAGAATGATAG |
| epikol_01509 | TET1_OXY_1 | GGAGCATACTGCTTATAAAT |
| epikol_01510 | TET1_OXY_2 | AACTCTGTAAGTTATCTTCA |
| epikol_01511 | TET1_OXY_3 | GTTGCCCGAGAATGTCGGCT |
| epikol_01512 | TET1_OXY_4 | GCGGTTATCTTCTCGAGTTA |
| epikol_01513 | TET1_OXY_5 | GCTTATAAAGAGGTAGCACA |
| epikol_01514 | TET2_OXY_1 | CCAAGGAAGTTTAAGCTGCT |
| epikol_01515 | TET2_OXY_2 | TTGGTGCCATAAGAGTGGAC |
| epikol_01516 | TET2_OXY_3 | GCAAAACCTGTCCACTCTTA |
| epikol_01517 | TET2_OXY_4 | AGAGCACCAGAGTGCCGTCT |
| epikol_01518 | TET2_OXY_5 | TTCAGACCCAGACGGCACTC |
| epikol_01519 | TET3_OXY_1 | CTCGCAAGTTCCGCCTCGCA |
| epikol_01520 | TET3_OXY_2 | TCAACGGCTGCAAGTATGCT |
| epikol_01521 | TET3_OXY_3 | GCCGTTGAAGTACATGCTCC |
| epikol_01522 | TET3_OXY_4 | CGCTCCCCTGTACAAGCGAC |
| epikol_01523 | TET3_OXY_5 | GTACAGGGGAGCGACTTCGG |
| epikol_01524 | TIP60_CHROM_1 | GGCCGAGATCCTGAGCGTGA |
| epikol_01525 | TIP60_CHROM_2 | GCGTGAAGGACATCAGTGGC |
| epikol_01526 | TIP60_CHROM_3 | ATGGGTGACGCATGAGCGGC |
| epikol_01527 | TIP60_CHROM_4 | ATGAATGGGTGACGCATGAG |
| epikol_01528 | TIP60_CHROM_5 | CACTGATGTCCTTCACGCTC |
| epikol_01529 | TIP60_NAT_1 | GCTTGCCGTAGCCCCGGCGC |
| epikol_01530 | TIP60_NAT_2 | CTGCCTCCCTACCAGCGCCG |
| epikol_01531 | TIP60_NAT_3 | TCAGCAGCTTGCCGTAGCCC |
| epikol_01532 | TIP60_NAT_4 | ATCAACGGAAGACTACAATG |

| | | |
|---|---|---|
| epikol_01533 | TIP60_NAT_5 | CCGGCGCTGGTAGGGAGGCA |
| epikol_01534 | TRIM24_BROMO_1 | TTACTGCCATGAAATGAGCC |
| epikol_01535 | TRIM24_BROMO_2 | GAACAGGGTCTTGAAAAGCC |
| epikol_01536 | TRIM24_BROMO_3 | AAATCAGCTACAAAATCTTC |
| epikol_01537 | TRIM24_BROMO_4 | GCATTGGCTACTTCTGAATC |
| epikol_01538 | TRIM24_BROMO_5 | GATTCAGAAGTAGCCAATGC |
| epikol_01539 | TRIM33_BROMO_1 | TACTTAATTCATGGCAATAG |
| epikol_01540 | TRIM33_BROMO_2 | CCACAAAGTCATCCGGGATT |
| epikol_01541 | TRIM33_BROMO_3 | CATCCGGGATTTGGTAGTGT |
| epikol_01542 | TRIM33_BROMO_4 | CTTGAAGATCAAACGGACAT |
| epikol_01543 | TRIM33_BROMO_5 | TGGACCTGTCAATCATCCGG |
| epikol_01544 | TRIM66_BROMO_1 | GGGCTGACAGGTTCATGGAA |
| epikol_01545 | TRIM66_BROMO_2 | GCTTCCTCCGGATGATTGAC |
| epikol_01546 | TRIM66_BROMO_3 | CGGATGATTGACAGGTCCAT |
| epikol_01547 | TRIM66_BROMO_4 | CCATGGGCCTCTTGATAATC |
| epikol_01548 | TRIM66_BROMO_5 | GCCTCTGCAACCTCGGAGTC |
| epikol_01549 | TTF2_DEXD_1 | GGATGAGCGCAATCATTGTC |
| epikol_01550 | TTF2_DEXD_2 | AGAAAAGCACAGCTTTGACG |
| epikol_01551 | TTF2_DEXD_3 | CTGGCACGTGAATCCCGGTT |
| epikol_01552 | TTF2_DEXD_4 | ATCTCTACCATGGGCCAAAC |
| epikol_01553 | TTF2_DEXD_5 | GATCACTACCTATAGCCTCG |
| epikol_01554 | TTF2_HELIC_1 | ATCTCTGTTGGCAGAATTGG |
| epikol_01555 | TTF2_HELIC_2 | CTGACTTATGCCACCATCGA |
| epikol_01556 | TTF2_HELIC_3 | GGGATTGACAGAGCCATCGA |
| epikol_01557 | TTF2_HELIC_4 | GAGGCATTTAACCACTCCAG |
| epikol_01558 | TTF2_HELIC_5 | GGTGTTGGTCTAAACCTGAC |
| epikol_01559 | UBE2A_UBC_1 | ACCATTATGTTGTTCTCGGA |
| epikol_01560 | UBE2A_UBC_2 | CATCATAGGTTGGACTCCAA |
| epikol_01561 | UBE2A_UBC_3 | TCCGTCCGAGAACAACATAA |
| epikol_01562 | UBE2A_UBC_4 | TCCTCCAGCCGGAGTCAGCG |
| epikol_01563 | UBE2A_UBC_5 | ATGGAAGACACATCATAGGT |
| epikol_01564 | UBE2B_UBC_1 | GGTGCGCCACTGACACCCAC |
| epikol_01565 | UBE2B_UBC_2 | TTACAAGAGGACCCACCTGT |
| epikol_01566 | UBE2B_UBC_3 | GATAAAAACCTAACAGTTGG |
| epikol_01567 | UBE2B_UBC_4 | TTGGACTCCATCGATTCTGA |
| epikol_01568 | UBE2B_UBC_5 | TTGGCTGGACTGTTAGGATT |
| epikol_01569 | UBE2E1_UBC_1 | GCCTCCAGGATCCGTGTATG |
| epikol_01570 | UBE2E1_UBC_2 | CCACCCTCATACACGGATCC |
| epikol_01571 | UBE2E1_UBC_3 | ACTTTAGAAATGGTTAGTGC |
| epikol_01572 | UBE2E1_UBC_4 | CAGTCAAGGTGTTATTTGCT |
| epikol_01573 | UBE2E1_UBC_5 | CTTCCCACCAAGGGGTCGGC |
| epikol_01574 | UBE2I_UBC_1 | TCGCCCAGGAGAGGAAAGCA |
| epikol_01575 | UBE2I_UBC_2 | AACTGGGAGTGCGCCATTCC |
| epikol_01576 | UBE2I_UBC_3 | ATGAGGTTCATCGTGCCATC |
| epikol_01577 | UBE2I_UBC_4 | GCACGATGAACCTCATGAAC |
| epikol_01578 | UBE2I_UBC_5 | ACCCGAATGTGTACCCTTCG |
| epikol_01579 | UBR2_GECKO_1 | ACTTAACACCTCTGAAATTG |
| epikol_01580 | UBR2_GECKO_2 | TGCATAACTTGAACTTTGAG |

| | | |
|---|---|---|
| epikol_01581 | UBR2_GECKO_3 | CTCTTCTTCCTCAATTTCAG |
| epikol_01582 | UBR2_GECKO_4 | TACACCATCTCTAAATCTGC |
| epikol_01583 | UBR2_GECKO_5 | ATTGTCGCACATCAGAATTT |
| epikol_01584 | UHRF1_1 | TCCCGTCCATGGTCCGAACC |
| epikol_01585 | UHRF1_2 | GGACAGCGAGTCCACCGTGT |
| epikol_01586 | UHRF1_3 | GTGGATCCAGGTTCGGACCA |
| epikol_01587 | UHRF1_4 | TGGACAGCGAGTCCACCGTG |
| epikol_01588 | UHRF1_5 | GGCGGACCTCGTAGTCGAAG |
| epikol_01589 | USP22_PC19_1 | CTTCATGAACTGCATCGTGC |
| epikol_01590 | USP22_PC19_2 | AGTCCCGCAGAAGTGGCGTG |
| epikol_01591 | USP22_PC19_3 | GCCTGCTAGGTGCCTCGCGT |
| epikol_01592 | USP22_PC19_4 | ACCTGGTGTGGACCCACGCG |
| epikol_01593 | USP22_PC19_5 | GCAACCCGCCTGTGAAGATC |
| epikol_01594 | USP22_UHYD_1 | CTTCATGAACTGCATCGTGC |
| epikol_01595 | USP22_UHYD_2 | CTGCGTGGGCTGATCAACCT |
| epikol_01596 | USP22_UHYD_3 | GCCTGCTAGGTGCCTCGCGT |
| epikol_01597 | USP22_UHYD_4 | ACCTGGTGTGGACCCACGCG |
| epikol_01598 | USP22_UHYD_5 | GCAACCCGCCTGTGAAGATC |
| epikol_01599 | USP44_PC19_1 | TTGTTGGGCGTAACCACGAA |
| epikol_01600 | USP44_PC19_2 | TTACGCCCAACAAGACGCTC |
| epikol_01601 | USP44_PC19_3 | GCTTGATCTGAACCAATGGC |
| epikol_01602 | USP44_PC19_4 | CCACTGCATTGATACCTTTC |
| epikol_01603 | USP44_PC19_5 | TGGGCTTCTGTGAGTACAAC |
| epikol_01604 | USP44_UHYD_1 | TTGTTGGGCGTAACCACGAA |
| epikol_01605 | USP44_UHYD_2 | TTACGCCCAACAAGACGCTC |
| epikol_01606 | USP44_UHYD_3 | AAATTCCTGAGCGTCTTGTT |
| epikol_01607 | USP44_UHYD_4 | CCACTGCATTGATACCTTTC |
| epikol_01608 | USP44_UHYD_5 | TATCTATGACTTGTCCGCGG |
| epikol_01609 | UTX_JMJC_1 | CAAACCATTCACAGTCACCT |
| epikol_01610 | UTX_JMJC_2 | GAATGGTTTGTTGTTCCTGA |
| epikol_01611 | UTX_JMJC_3 | GTTGTTCCTGAAGGTTACTG |
| epikol_01612 | UTX_JMJC_4 | GTTCATTGGGTTCAGGCTAT |
| epikol_01613 | UTX_JMJC_5 | TGGGCCACCAAGAACCCATT |
| epikol_01614 | UTY_JMJC_1 | GTTGTACCTGAAGATTATTG |
| epikol_01615 | UTY_JMJC_2 | TGTTGTACCTGAAGATTATT |
| epikol_01616 | UTY_JMJC_3 | GTGCATTGGGTTCAAGCTGT |
| epikol_01617 | UTY_JMJC_4 | TTGCTTCATAAAGATCTTCA |
| epikol_01618 | UTY_JMJC_5 | TTGTTGTACCTGAAGATTAT |
| epikol_01619 | WDR5_WD40_1 | AAATTTGGGGCGCGTATGAT |
| epikol_01620 | WDR5_WD40_2 | ATCTGACGACCAGGCTACAT |
| epikol_01621 | WDR5_WD40_3 | CAAGACTTTGCCAGCTCACT |
| epikol_01622 | WDR5_WD40_4 | ATAGCTACTTGAAACTATCA |
| epikol_01623 | WDR5_WD40_5 | CAGGATGTATTTGCCGTTCG |
| epikol_01624 | WDR82_WD40_1 | TTTCAGCCCCAACGGCGAGA |
| epikol_01625 | WDR82_WD40_2 | ACGAATGAAGCTGCCGTTGG |
| epikol_01626 | WDR82_WD40_3 | CGTCTGATTGATGCATTCAA |
| epikol_01627 | WDR82_WD40_4 | CGAGATGACCGTCTCGCCGT |
| epikol_01628 | WDR82_WD40_5 | CTGCATGAGTGTATCTGATG |

| | | |
|---|---|---|
| epikol_01629 | ZMYND11_BROMO_1 | ATTCATTGTCTCCCGCATGA |
| epikol_01630 | ZMYND11_BROMO_2 | ACAATAAACACCCGATGTAC |
| epikol_01631 | ZMYND11_BROMO_3 | ACACCCGATGTACAGGAGGC |
| epikol_01632 | ZMYND11_BROMO_4 | CGGTATTGTGGAGAAGCAAT |
| epikol_01633 | ZMYND11_BROMO_5 | GTGAGCAAGCTGACATTGCG |
| epikol_01634 | ZMYND8_BROMO_1 | AATGGCAAACTTGAGCAGGT |
| epikol_01635 | ZMYND8_BROMO_2 | ATTCCAGAAGCCCGTTCCAT |
| epikol_01636 | ZMYND8_BROMO_3 | GGGTGCTGTTCCAATGGAAC |
| epikol_01637 | ZMYND8_BROMO_4 | AAATCCACTTTGCATCAGCC |
| epikol_01638 | ZMYND8_BROMO_5 | TGGCTGCACAGAAGCCTTCC |
| epikol_01639 | NonTargetingControlGuideForHuman_0001 | ACGGAGGCTAAGCGTCGCAA |
| epikol_01640 | NonTargetingControlGuideForHuman_0002 | CGCTTCCGCGGCCCGTTCAA |
| epikol_01641 | NonTargetingControlGuideForHuman_0003 | ATCGTTTCCGCTTAACGGCG |
| epikol_01642 | NonTargetingControlGuideForHuman_0004 | GTAGGCGCGCCGCTCTCTAC |
| epikol_01643 | NonTargetingControlGuideForHuman_0005 | CCATATCGGGGCGAGACATG |
| epikol_01644 | NonTargetingControlGuideForHuman_0006 | TACTAACGCCGCTCCTACAG |
| epikol_01645 | NonTargetingControlGuideForHuman_0007 | TGAGGATCATGTCGAGCGCC |
| epikol_01646 | NonTargetingControlGuideForHuman_0008 | GGGCCCGCATAGGATATCGC |
| epikol_01647 | NonTargetingControlGuideForHuman_0009 | TAGACAACCGCGGAGAATGC |
| epikol_01648 | NonTargetingControlGuideForHuman_0010 | ACGGGCGGCTATCGCTGACT |
| epikol_01649 | NonTargetingControlGuideForHuman_0011 | CGCGGAAATTTTACCGACGA |
| epikol_01650 | NonTargetingControlGuideForHuman_0012 | CTTACAATCGTCGGTCCAAT |
| epikol_01651 | NonTargetingControlGuideForHuman_0013 | GCGTGCGTCCCGGGTTACCC |
| epikol_01652 | NonTargetingControlGuideForHuman_0014 | CGGAGTAACAAGCGGACGGA |
| epikol_01653 | NonTargetingControlGuideForHuman_0015 | CGAGTGTTATACGCACCGTT |
| epikol_01654 | NonTargetingControlGuideForHuman_0016 | CGACTAACCGGAAACTTTTT |
| epikol_01655 | NonTargetingControlGuideForHuman_0017 | CAACGGGTTCTCCCGGCTAC |
| epikol_01656 | NonTargetingControlGuideForHuman_0018 | CAGGAGTCGCCGATACGCGT |
| epikol_01657 | NonTargetingControlGuideForHuman_0019 | TTCACGTCGTCTCGCGACCA |
| epikol_01658 | NonTargetingControlGuideForHuman_0020 | GTGTCGGATTCCGCCGCTTA |
| epikol_01659 | NonTargetingControlGuideForHuman_0021 | CACGAACTCACACCGCGCGA |
| epikol_01660 | NonTargetingControlGuideForHuman_0022 | CGCTAGTACGCTCCTCTATA |
| epikol_01661 | NonTargetingControlGuideForHuman_0023 | TCGCGCTTGGGTTATACGCT |
| epikol_01662 | NonTargetingControlGuideForHuman_0024 | CTATCTCGAGTGGTAATGCG |
| epikol_01663 | NonTargetingControlGuideForHuman_0025 | AATCGACTCGAACTTCGTGT |
| epikol_01664 | NonTargetingControlGuideForHuman_0026 | CCCGATGGACTATACCGAAC |
| epikol_01665 | NonTargetingControlGuideForHuman_0027 | ACGTTCGAGTACGACCAGCT |
| epikol_01666 | NonTargetingControlGuideForHuman_0028 | CGCGACGACTCAACCTAGTC |
| epikol_01667 | NonTargetingControlGuideForHuman_0029 | GGTCACCGATCGAGAGCTAG |
| epikol_01668 | NonTargetingControlGuideForHuman_0030 | CTCAACCGACCGTATGGTCA |
| epikol_01669 | NonTargetingControlGuideForHuman_0031 | CGTATTCGACTCTCAACGCG |
| epikol_01670 | NonTargetingControlGuideForHuman_0032 | CTAGCCGCCCAGATCGAGCC |
| epikol_01671 | NonTargetingControlGuideForHuman_0033 | GAATCGACCGACACTAATGT |
| epikol_01672 | NonTargetingControlGuideForHuman_0034 | ACTTCAGTTCGGCGTAGTCA |
| epikol_01673 | NonTargetingControlGuideForHuman_0035 | GTGCGATGTCGCTTCAACGT |
| epikol_01674 | NonTargetingControlGuideForHuman_0036 | CGCCTAATTTCCGGATCAAT |
| epikol_01675 | NonTargetingControlGuideForHuman_0037 | CGTGGCCGGAACCGTCATAG |
| epikol_01676 | NonTargetingControlGuideForHuman_0038 | ACCCTCCGAATCGTAACGGA |

| | | |
|---|---|---|
| epikol_01677 | NonTargetingControlGuideForHuman_0039 | AAACGGTACGACAGCGTGTG |
| epikol_01678 | NonTargetingControlGuideForHuman_0040 | ACATAGTCGACGGCTCGATT |
| epikol_01679 | NonTargetingControlGuideForHuman_0041 | GATGGCGCTTCAGTCGTCGG |
| epikol_01680 | NonTargetingControlGuideForHuman_0042 | ATAATCCGGAAACGCTCGAC |
| epikol_01681 | NonTargetingControlGuideForHuman_0043 | CGCCGGGCTGACAATTAACG |
| epikol_01682 | NonTargetingControlGuideForHuman_0044 | CGTCGCCATATGCCGGTGGC |
| epikol_01683 | NonTargetingControlGuideForHuman_0045 | CGGGCCTATAACACCATCGA |
| epikol_01684 | NonTargetingControlGuideForHuman_0046 | CGCCGTTCCGAGATACTTGA |
| epikol_01685 | NonTargetingControlGuideForHuman_0047 | CGGGACGTCGCGAAAATGTA |
| epikol_01686 | NonTargetingControlGuideForHuman_0048 | TCGGCATACGGGACACACGC |
| epikol_01687 | NonTargetingControlGuideForHuman_0049 | AGCTCCATCGCCGCGATAAT |
| epikol_01688 | NonTargetingControlGuideForHuman_0050 | ATCGTATCATCAGCTAGCGC |
| epikol_01689 | NonTargetingControlGuideForHuman_0051 | TCGATCGAGGTTGCATTCGG |
| epikol_01690 | NonTargetingControlGuideForHuman_0052 | CTCGACAGTTCGTCCCGAGC |
| epikol_01691 | NonTargetingControlGuideForHuman_0053 | CGGTAGTATTAATCGCTGAC |
| epikol_01692 | NonTargetingControlGuideForHuman_0054 | TGAACGCGTGTTTCCTTGCA |
| epikol_01693 | NonTargetingControlGuideForHuman_0055 | CGACGCTAGGTAACGTAGAG |
| epikol_01694 | NonTargetingControlGuideForHuman_0056 | CATTGTTGAGCGGGCGCGCT |
| epikol_01695 | NonTargetingControlGuideForHuman_0057 | CCGCTATTGAAACCGCCCAC |
| epikol_01696 | NonTargetingControlGuideForHuman_0058 | AGACACGTCACCGGTCAAAA |
| epikol_01697 | NonTargetingControlGuideForHuman_0059 | TTTACGATCTAGCGGCGTAG |
| epikol_01698 | NonTargetingControlGuideForHuman_0060 | TTCGCACGATTGCACCTTGG |
| epikol_01699 | NonTargetingControlGuideForHuman_0061 | GGTTAGAGACTAGGCGCGCG |
| epikol_01700 | NonTargetingControlGuideForHuman_0062 | CCTCCGTGCTAACGCGGACG |
| epikol_01701 | NonTargetingControlGuideForHuman_0063 | TTATCGCGTAGTGCTGACGT |
| epikol_01702 | NonTargetingControlGuideForHuman_0064 | TACGCTTGCGTTTAGCGTCC |
| epikol_01703 | NonTargetingControlGuideForHuman_0065 | CGCGGCCCACGCGTCATCGC |
| epikol_01704 | NonTargetingControlGuideForHuman_0066 | AGCTCGCCATGTCGGTTCTC |
| epikol_01705 | NonTargetingControlGuideForHuman_0067 | AACTAGCCCGAGCAGCTTCG |
| epikol_01706 | NonTargetingControlGuideForHuman_0068 | CGCAAGGTGTCGGTAACCCT |
| epikol_01707 | NonTargetingControlGuideForHuman_0069 | CTTCGACGCCATCGTGCTCA |
| epikol_01708 | NonTargetingControlGuideForHuman_0070 | TCCTGGATACCGCGTGGTTA |
| epikol_01709 | NonTargetingControlGuideForHuman_0071 | ATAGCCGCCGCTCATTACTT |
| epikol_01710 | NonTargetingControlGuideForHuman_0072 | GTCGTCCGGGATTACAAAAT |
| epikol_01711 | NonTargetingControlGuideForHuman_0073 | TAATGCTGCACACGCCGAAT |
| epikol_01712 | NonTargetingControlGuideForHuman_0074 | TATCGCTTCCGATTAGTCCG |
| epikol_01713 | NonTargetingControlGuideForHuman_0075 | GTACCATACCGCGTACCCTT |
| epikol_01714 | NonTargetingControlGuideForHuman_0076 | TAAGATCCGCGGGTGGCAAC |
| epikol_01715 | NonTargetingControlGuideForHuman_0077 | GTAGACGTCGTGAGCTTCAC |
| epikol_01716 | NonTargetingControlGuideForHuman_0078 | TCGCGGACATAGGGCTCTAA |
| epikol_01717 | NonTargetingControlGuideForHuman_0079 | AGCGCAGATAGCGCGTATCA |
| epikol_01718 | NonTargetingControlGuideForHuman_0080 | GTTCGCTTCGTAACGAGGAA |

**Supplementary Table1 Sequences and targets of sgRNAs of EpiDoKOL**

| gRNA ID | Sequence |
|---|---|
| g-T1 | TGAACCGCATCGAGCTGAA |
| g-T2 | GAGCGCACCATCTTCTTCA |
| g-NT1 | ACGGAGGCTAAGCGTCGCAA |
| g-NT2 | CGCTTCCGCGGCCCGTTCAA |
| g-ASH2L | CACCGGCCTCTCATGGAGTACGGAA |
| g-RBX1 | CACCGACTTCCACTGCATCTCTCGC |
| g-SSRP1 | CACCGGACTGCTCTACCCGCTGGAG |

**Supplementary Table2 All sgRNA sequences used for EpiDoKOL Cas9 activity and essentiality validation experiments.**

| Internal PCR Primers | Sequences |
|---|---|
| **Forward Mix Primers** | |
| Forward_stag1 | AATGATACGGCGACCACCGAGATCTACACTCTTTCCCTACACGACGCTCTTCCGATCTATCTTGTGGAAAGGACGA[illegible] |
| Forward_stag2 | AATGATACGGCGACCACCGAGATCTACACTCTTTCCCTACACGACGCTCTTCCGATCTCTTCTTGTGGAAAGGACG[illegible] |
| Forward_stag3 | AATGATACGGCGACCACCGAGATCTACACTCTTTCCCTACACGACGCTCTTCCGATCTGACTCTTGTGGAAAGGAC[illegible] |
| Forward_stag4 | AATGATACGGCGACCACCGAGATCTACACTCTTTCCCTACACGACGCTCTTCCGATCTTCGATCTTGTGGAAAGGA[illegible] |
| Forward_stag5 | AATGATACGGCGACCACCGAGATCTACACTCTTTCCCTACACGACGCTCTTCCGATCTATCGCTCTTGTGGAAAGG[illegible] |
| Forward_stag6 | AATGATACGGCGACCACCGAGATCTACACTCTTTCCCTACACGACGCTCTTCCGATCTGTCAGATCTTGTGGAAAG[illegible] |
| Forward_stag7 | AATGATACGGCGACCACCGAGATCTACACTCTTTCCCTACACGACGCTCTTCCGATCTTGCATCGTCTTGTGGAAAG |
| Forward_stag8 | AATGATACGGCGACCACCGAGATCTACACTCTTTCCCTACACGACGCTCTTCCGATCTCTACGTGATCTTGTGGAAA |
| Forward_stag9 | AATGATACGGCGACCACCGAGATCTACACTCTTTCCCTACACGACGCTCTTCCGATCTACTCGTGATTCTTGTGGAA[illegible] |
| **Reverse Index Primers** | |
| rev_index1 | CAAGCAGAAGACGGCATACGAGATCGTGATGTGACTGGAGTTCAGACGTGTGCTCTTCCGATCTTCTACTATTCTTT |
| rev_index3 | CAAGCAGAAGACGGCATACGAGATGCCTAAGTGACTGGAGTTCAGACGTGTGCTCTTCCGATCTTCTACTATTCTTT |
| rev_index4 | CAAGCAGAAGACGGCATACGAGATTGGTCAGTGACTGGAGTTCAGACGTGTGCTCTTCCGATCTTCTACTATTCTTT |
| rev_index7 | CAAGCAGAAGACGGCATACGAGATGATCTGGTGACTGGAGTTCAGACGTGTGCTCTTCCGATCTTCTACTATTCTTT |
| rev_index9 | CAAGCAGAAGACGGCATACGAGATCTGATCGTGACTGGAGTTCAGACGTGTGCTCTTCCGATCTTCTACTATTCTTT[illegible] |
| rev_index10 | CAAGCAGAAGACGGCATACGAGATAAGCTAGTGACTGGAGTTCAGACGTGTGCTCTTCCGATCTTCTACTATTCTTT |
| **External PCR Primers** | |
| **External Forward Primer** | AATGGACTATCATATGCTTACCGTAACTTGAAAGTATTTCG |
| **External Reverse Primer** | TCTACTATTCTTTCCCCTGCACTGTTGTGGGCGATGTGCGCTCTG |

**Supplementary Table3 Sequences of external and internal PCR primers used for Nested-PCR**

| | |
|---|---|
| **BARD1-qPCR-F** | TGCAGCCAAGAATGGGCATGTG |
| **BARD1-qPCR-R** | CTTCTCTGGTAGCAGCAATAGCG |
| **PDS5B-qPCR-F** | CTCTTGGAGAGGATAGCACCTG |
| **PDS5B-qPCR-R** | GACCTGCTCTGATGGCTTGATC |
| **RAD21-qPCR-F** | GTGGAAAGAGACAGGAGGAGTAG |
| **RAD21-qPCR-R** | AGGTCTTCTGGTACAAGCGGTG |
| **ASH2L-qPCR-F** | AGAATGGCCGACAGTTGGG |
| **ASH2L-qPCR-R** | CCTTCAAGTTTGCTTGCTTCC |

**Supplementary Table4 Sequences qPCR primers used for RNAseq validation**
